# A Cysteine-Less and Ultra-Fast Split Intein Rationally Engineered from Being Aggregation-Prone to Highly Efficient in Protein trans-Splicing

**DOI:** 10.1101/2025.01.22.634254

**Authors:** Christoph Humberg, Zahide Yilmaz, Katharina Fitzian, Wolfgang Dörner, Daniel Kümmel, Henning D. Mootz

## Abstract

Split inteins ligate their fused extein protein sequences while undergoing self-removal. This unique protein *trans*-splicing reaction has been harnessed for numerous applications in protein modification. However, several purified split intein precursors splice only partially or are entirely inactive for unknown reasons. We have studied the split Aes123 PolB1 intein, which splices only to about 30%. As a rare representative of cysteine-less split inteins, the Aes intein is attractive due to its resistance to oxidative conditions and orthogonality to thiol chemistries. We revealed that the reduced splicing efficiency is caused by the formation of soluble, β-sheet dominated aggregates of the N-terminal precursor. We computationally, biochemically and biophysically characterized the fully active monomeric fraction to identify sequence regions important for aggregation. Guided by a crystal structure we designed stably monomeric mutants with virtually complete splicing activity. The triple mutant CLm intein (Cysteine-Less and monomeric) retained the ultra-fast rate discovered for the wildtype and exhibits superior utility as a thiol-independent protein modification tool. Characterization of two other benchmark split inteins suggests that the discovered aggregation propensity reflects an inherent challenge to keep the split intein precursor in the partially disordered form required to fold with its complementary partner into the active complex.

## Introduction

Inteins are autoprocessing domains that mediate the virtually traceless ligation of their flanking extein polypeptide sequences in a precursor protein with their own concomitant removal.^1, 2^ This process is termed protein splicing and encompasses a coordinated multistep rearrangement of the polypeptide backbone (Supplementary Fig. 1).^3–7^ It typically involves two thioester or oxyester intermediates on Cys, Ser or Thr side chains at the two splice junctions between exteins and intein. In split inteins two separate precursor proteins with Int^N^ and Int^C^ fragments, P_N_ and P_C_, first associate and fold into the active intein domain before undergoing a mechanistically analogous reaction termed protein *trans*-splicing (Fig. 1A and Supplementary Fig. 1).^4–6, 8^

**Figure 1.**
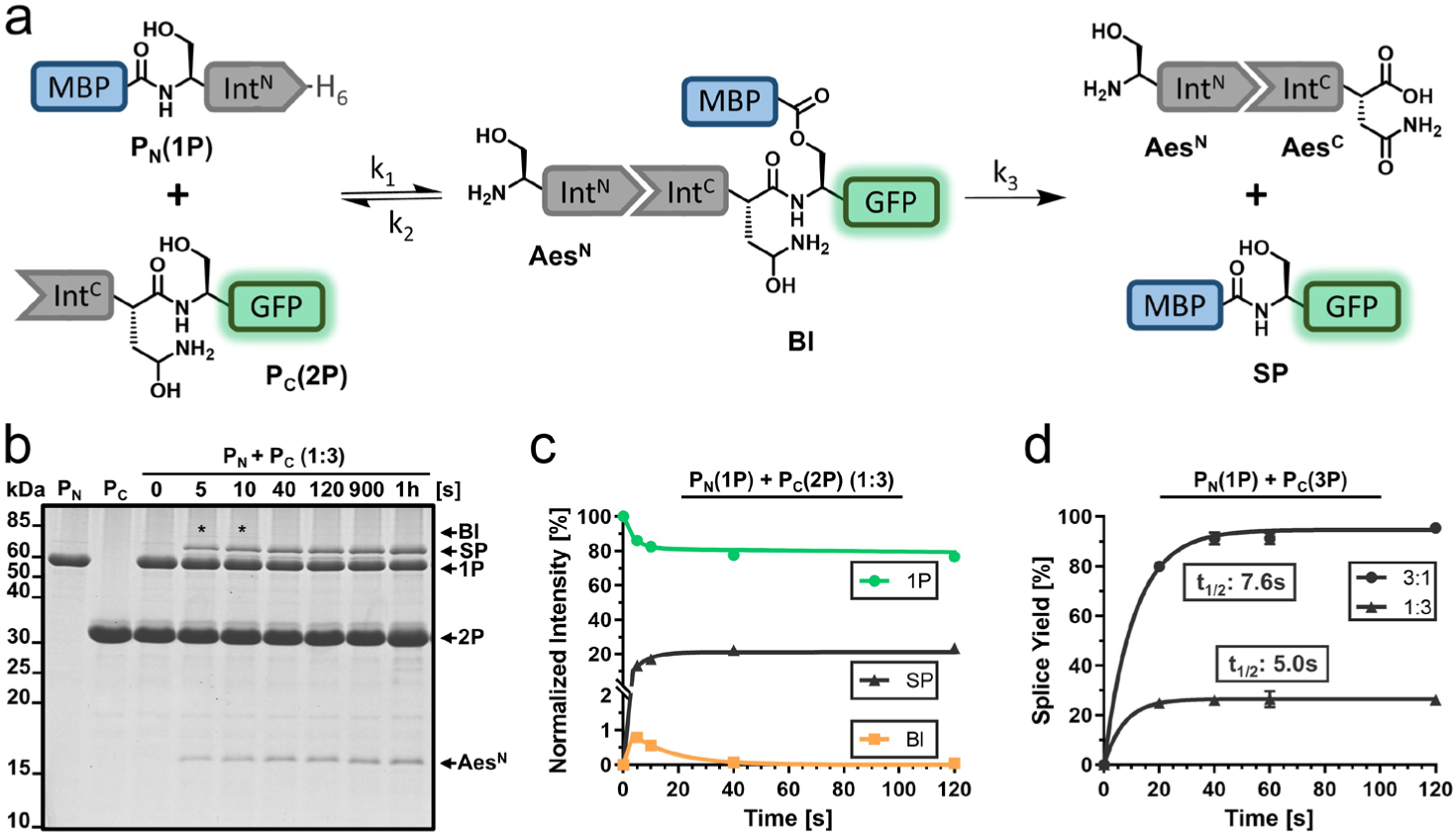
Protein *trans*-splicing of the wildtype Aes123 PolB1 intein. **a** Scheme of the PTS reaction in a simplified three-state kinetic model. Not shown are the linear intermediate resulting from the *N-O* acyl shift and the C-terminal succinimide prior to hydrolysis. **b** Analysis of the PTS reaction at 37 °C using precursors **1P** and **2P** at 10 µM and 30 µM, respectively, and in absence of any reducing agents. Shown is a Coomassie-stained SDS-PAGE gel. **c** Time-course of the PTS reaction obtained by densitometric analysis of data as shown in **b** and fitted to the three-state kinetic model. **d** Splice product formation plotted against the time applying different molar ratios of the precursors **1P** and **3P**. Calculated molecular masses are: P_N_(1P) = 58.2 kDa, P_C_(2P) = 31.3 kDa, BI = 74.8 kDa, SP = 70.1 kDa, Aes^N^ = 14.7 kDa, Aes^C^ = 4.7 kDa; BI = branched intermediate, SP = splice product. For panel **c** and **d**, *n* = 3 technical replicates. Data are presented as mean ± s.d. normalized to the molecular weight of each protein species.

Inteins have been enabled many protein engineering technologies that rely on their unique making and breaking of peptide bonds.^7, 9–11^ Split inteins are of particular potential for protein labeling, semi-synthesis and reconstitution applications as they assemble a peptide backbone from two separate proteins that are individually producible and modifiable. A high *trans*-splicing efficiency, a high rate and high affinity are the most important activity traits of a split intein to be useful for effective manipulation of both purified proteins and proteins in living cells. However, these properties are exhibited by only very few split inteins. Predicting these properties from the sequence is currently impossible and hence each newly identified intein requires an elaborate biochemical and biophysical characterization.

We have recently introduced the first two cysteine-less split inteins as a new tool with powerful expanded protein engineering capabilities.^12, 13^ Cysteine-less split inteins employ only oxyester intermediates, i.e., they form a rare subgroup that exhibit only serine or threonine residues at the splice junctions.^14^ Ideally, also the remainder of the intein sequences contains no or only non-essential cysteines. Therefore, cysteine-less split inteins are fully active without the addition of reducing agents typically applied for cysteine-dependent inteins to prevent oxidative inactivation. This way, disulfide bonds in the exteins remain protected, for example. By being inert against oxidative conditions and orthogonal to cysteine-directed bioconjugation reagents, cysteine-less inteins also enable attractive new schemes that exploit cysteine chemistry in the extein sequences, or involve protein *trans*-splicing in the oxidative extracellular milieu, for example.^12^

Importantly, however, a cysteine-less split intein with the desired features of high-efficiency, high-rate and high-affinity splicing is still lacking so far. The previously reported cysteine-less Aes123 PolB1 intein (in short: Aes intein), naturally split at the endonuclease position, showed only about 30% splicing efficiency with regard to its P_N_.^12^ By shifting the split site to an atypical position the efficiency could be significantly increased, however, at the cost of a strongly reduced rate of the PTS reaction.^12^ A cysteine-less split intein with the important favorable features is thus still urgently sought after.

Split inteins occur either naturally or are artificially generated from *cis*-splicing inteins by genetically breaking the polypeptide chain of the conserved intein fold of about 130-150 aa.^8, 15–17^ The most common split site coincides with a position where most *cis*-inteins (the so-called maxi-inteins) harbor an insertion of a homing endonuclease domain (Supplementary Fig. 1A).^18^ *Cis*-inteins without this insertion are called mini-inteins.^19^ This split position yields Int^N^ and Int^C^ fragments of about 100 aa and 40 aa and is also typically found in naturally split inteins, such as the widely used *Npu* DnaE and Gp41-1 inteins.^20–22^ Atypically split inteins with a short Int^N^ and long Int^C^ fragment have later been identified.^23–25^

Naturally split inteins are generally considered to be more favorable in protein *trans*-splicing due to their evolutionary adaptation for this function. However, poor efficiencies^21, 26^ and a wide range of rates with half-life times from seconds to hours^21, 22, 27^ can also be found in this group.

Furthermore, for the same intein the efficiency may be complete or only partial depending on different exteins, however, irrespective of the immediately flanking residues, as observed for the *Npu* DnaE intein,^21^ for example. Artificially splitting at the endonuclease position has yielded many active split inteins.^16, 17^ However, these showed good efficiency only when the two artificially split fragments are then co-expressed inside cells.^17^ In contrast, when the P_N_ and P_C_ are separately purified before their combination, the efficiencies were typically limited to 40-50% at best.^28–30^ In other cases, such artificially split intein precursors were even nearly inactive, but were capable of splicing when collectively refolded from denaturing conditions.^15, 31–33^ This latter procedure also denatures the extein proteins, therefore limiting the practical utility, also of an earlier reported cysteine-less split intein.^31^ In contrast, artificially splitting inteins at atypical positions has yielded a few examples with very high splicing efficiency of the purified precursors, first reported for the *Ssp* DnaB intein with an 11 aa Int^N^ fragment.^12, 34^ Overall, the molecular origins for incomplete protein *trans*-splicing, typically correlated with incomplete consumption of the P_N_, remain entirely unknown.

Initial association of the precursor proteins split at the endonuclease position is electrostatically driven by opposite charges in disordered regions of each fragment (‘capture’),^35, 36^ followed by folding into the conserved and intertwined horseshoe structure of inteins that triggers protein *trans*-splicing. The transition from disordered to ordered (‘collapse’) becomes thermodynamically favorable through the formation of a structure rich in β-strands.^35, 36^ For the *Npu* DnaE intein, the opposite charges have been mapped to disordered regions in the C-terminal of the two symmetry-related lobes (N_2_) within the Int^N^ fragment, and to the Int^C^ fragment. In contrast, the N-terminal lobe of the Int^N^ fragment (N_1_) exhibits residual, molten-globule-like structure before fragment association.^36^

In this work, we revisited the Aes intein to understand the molecular origins for its limited efficiency. We discovered that a highly aggregated, yet soluble P_N_ caused the partial inactivity. We identified regions in the disordered part of the Int^N^ that are prone to aggregation and obtained by rational mutagenesis a fully monomeric variant with excellent efficiency. Combined with its ultra-fast rate, the engineered Aes intein is a highly valuable addition to the protein chemistry toolbox. We further show that the formation of soluble aggregates is a more widely occurring feature of both artificially and naturally split inteins, thus providing an explanation for a long-standing bottleneck in intein research.

## Results

### The active fraction of the cysteine-less Aes split intein splices at an ultra-fast rate

We revisited the naturally occurring Aes split intein as a detailed biochemical characterization was abandoned in a previous study due to its poor activity.^12^ We prepared recombinant model N- and C-terminal precursor proteins MBP-Aes^N^-H_6_ (**1P**; P_N_), Aes^C^-GFP (**2P**; P_C_) and SBP-Aes^C^-SBP (**3P**; P_C_) with maltose-binding protein (MBP), green fluorescent protein (GFP) or streptavidin-binding peptide (SBP) as exteins, respectively (Fig. 1A). Proteins were produced in *E. coli* and purified using Ni-NTA or streptactin affinity chromatography. In the protein samples and assays no reducing agents were applied. Upon mixing complementary constructs **1P** and **2P** in a 1:3 ratio at 37°C, we observed splice product formation of about 30%, relative to the limiting Aes^N^ precursor **1P** (Fig. 1B). These results confirmed the inteins poor efficiency,^12^ which significantly limits its utility.

Interestingly, however, the reaction showed an ultra-fast rate of 138.7 ± 18.6 x 10^-3^ s^-1^ (t_½_ = 5.0 s) at 37°C (Fig. 1C). A three-fold molar excess of the Aes^N^ precursor led to virtually quantitative splicing of the Aes^C^ precursor (**1P** + **3P**) with a rate of 91.3 ± 6.2 x 10^-3^ s^-1^ (t_½_ = 7.6 s) (Fig. 1D, Supplementary Fig. 2). N- or C-cleavage as potential side reactions were below the detection limit. This ultra-fast splicing rate of the Aes intein was previously overlooked and ranks this intein among the fastest known, e.g. the Gp41-1 intein (t_½_ = 5 s at 37°C)^22, 37^ and the NrdJ-1 intein (t_½_ = 7 s),^22^ and more than 10-fold faster than the *Npu* DnaE intein (t_½_ = 63 s),^21^ all of which are cysteine-dependent.

### Concentration-dependent aggregate formation of the Aes^N^ precursor is yield limiting

To address the large inactivity of the Aes^N^ precursor we hypothesized it partially misfolds in the absence of its P_C_ partner. To test co-folding of both P_N_ and P_C_, we denatured separately purified **1P** and **3P** with 8 M urea and then mixed both proteins in a 1:3 molar ratio (10 μM **1P** + 30 μM **3P**), followed by removal of the denaturant by dialysis. Indeed, this procedure led to virtually complete conversion of **1P** into splice product (Supplementary Fig. 3). To investigate P_N_ conformation in solution, we analyzed **1P** (m_calc_ = 58.2 kDa) by size exclusion chromatography (SEC) (Fig. 2A,B). The protein appeared in two well-separated fractions.

**Figure 2.**
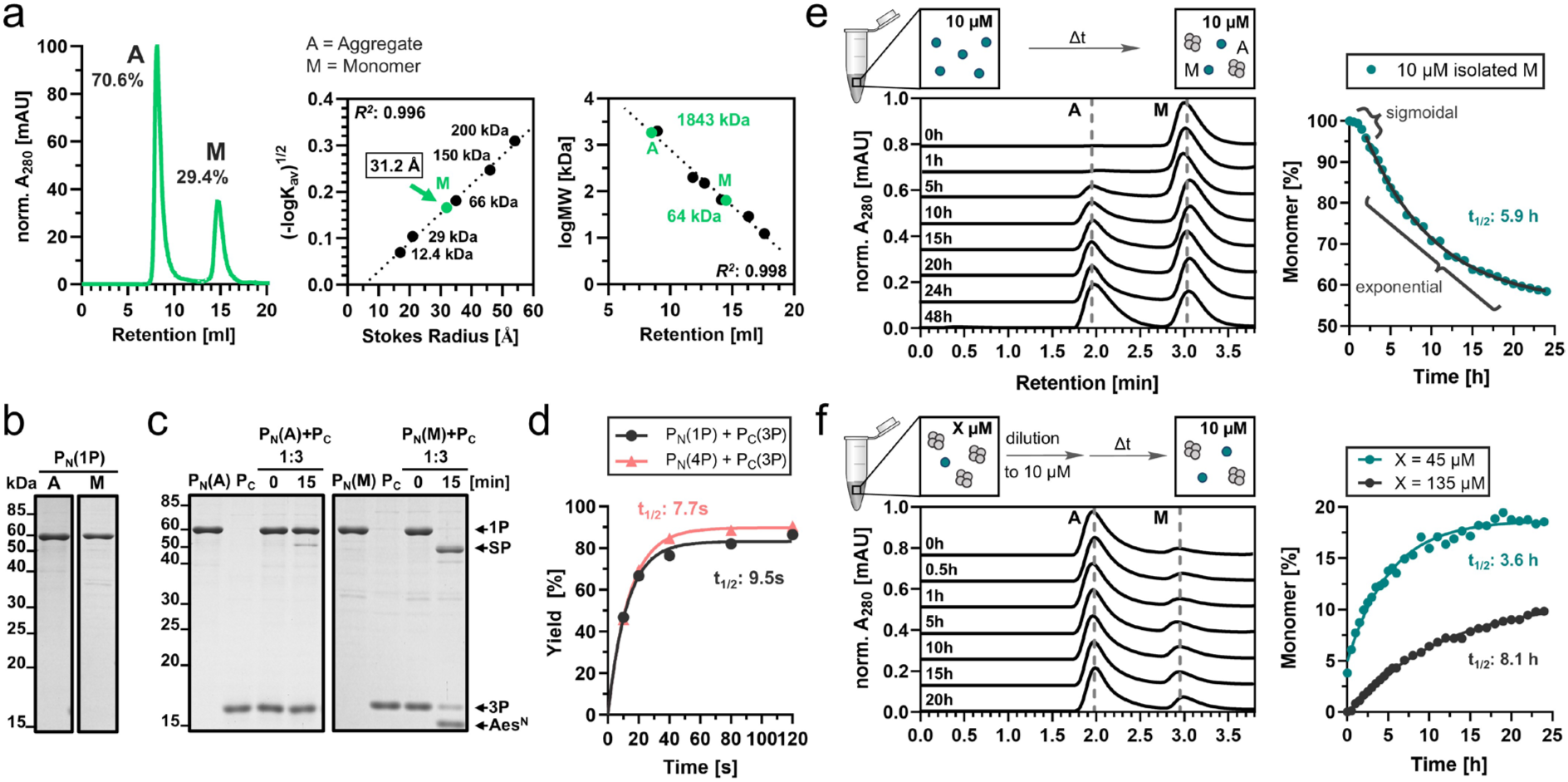
The Aes^N^ precursor **1P** exhibits two folding states as revealed by SEC. **a** SEC analysis of **1P** (left panel). This data and a calibration curve generated with globular protein standards were used to determine the Stokes radius (middle panel) and the apparent molecular weight of the aggregated (A) and monomeric (M) species. Because the aggregated species eluted shortly before dextran as the void volume marker, we could only estimate its hydrodynamic radius to be >270 Å. **b** Analysis of aggregated and monomeric fractions on a Coomassie-stained SDS-PAGE gel. **c** Analysis of PTS reactions using the aggregated and monomeric species of **1P** (10 µM) with three-fold excess of **3P** (30 µM) at 37 °C. Shown are Coomassie-stained SDS-PAGE gels. **d** Time-courses of two PTS reactions using monomeric fractions of the indicated Aes^N^ precursors **1P** and **4P** with a three-fold excess of **3P** (see also Supplementary Fig. 4). **e** Analysis of concentration-dependent re-aggregation of **1P** over time. The left panel shows representative analytical SEC profiles recorded at indicated time points following the isolation of its monomeric fraction adjusted to 10 µM. The right panel shows the plot of the monomeric fraction. Note that only the second, exponential phase was used for fitting. The sigmoidal appearance of the curve at early time points is expected as a typical feature of amyloid aggregate formation due to a slow nucleation (lag phase) before aggregation proceeds as a concentration-dependent polymerization reaction (exponential phase) until reaching saturation (equilibrium phase).^46^ **f** Analysis of concentration-dependent resolution of **1P** aggregate. The left panel shows representative analytical SEC profiles recorded at indicated time points after dilution of an aggregate fraction from 45 µM to 10 µM. The right panel shows the plotted aggregate resolution to the monomeric species from starting concentrations 135 µM and 45 µM, each fitted to a one-phase exponential equation. For panel **a** (middle and right inlets), *n* = 3 technical replicates. For panel **d**, n = 2 replicates. Data are presented as mean band intensity in SDS-PAGE analysis ± s.d. normalized to the molecular weight. For the plots shown in panel **e** and **f**, *n* = 1.

Fraction A (about 71% of the protein) eluted at the void volume of the column, thus corresponding to an apparent molecular weight (MW) of at least ca. 2,000 kDa, indicative of a highly oligomeric or aggregated form of **1P**. The retention time of fraction B (about 29 %) translated into a Stoke radius of 31.2 Å, which reflects an apparent MW of about 64 kDa, in good consistency with a monomeric and partially disordered form. Most importantly, the aggregated fraction of **1P** showed only traces of PTS activity with an excess of P_C_ **3P**, while the monomeric fraction spliced virtually quantitatively under these conditions (Fig. 2C). These findings explained the partial activity of the Aes^N^ precursor. Variation of the N-extein to GFP in construct GFP-Aes^N^-H_6_ (**4P**) did not change the occurrence of the aggregated fraction, suggesting that aggregation is an inherent property of the Aes^N^ fragment (Supplementary Fig. 4).

Attempts to prevent the aggregated fraction by varying the expression conditions or applying different salt concentrations failed (Supplementary Fig. 4). We therefore developed a refolding protocol to obtain monomeric P_N_, for further analysis. Following Ni-NTA purification under denaturing conditions, **1P** and **4P** were step-wisely refolded by dialysis in presence of sucrose as a stabilizer. Both proteins prepared this way exhibited nearly quantitative PTS activity and retained ultra-fast rates with t_1/2_ = 7.7 and 9.5 s with partner protein SBP-Aes^C^-SBP (**3P**) (Fig. 2D, Supplementary Fig. 4). Using biolayer interferometry, we investigated the binding affinity of the monomeric split precursors inactivated by mutations to block splicing (Supplementary Fig. 5). A high affinity with a *K*_d_ value of 12 nM with a *k*_on_ of 4.9 ± 1.8 x 10^4^ M^-1^s^-1^ and *k*_off_ of 7.5 ± 1.1 x 10^-4^ s^-1^ revealed an additional favorable property of the split Aes intein next to its rate and cysteine-less nature. Assuming splice competence of the intein fragments, simulation of the underlying binding kinetics even indicated a *K*_d_ of 0.5 nM (Supplementary Fig. 6, Supplementary Table 5). Furthermore, the binding data revealed a biphasic behavior consistent with the assumed split intein assembly mechanism (see Supplementary Note 1).

However, we then discovered that the elaborate refolding protocol of the active, monomeric P_N_ fractions was foiled as a useful preparative procedure by the tendency of the P_N_ to re-aggregate and form a monomer-aggregate ratio in a concentration-dependent manner (Fig. 2E). The ∼30% of monomeric form of **1P** described so far was observed for an isolated protein concentration of ∼10-15 µM, while at 45 µM and 135 µM only about 4% and no detectable monomeric species were present (Fig. 2F). Consequently, protein dilution induced disaggregation, however, at inconvenient concentrations and time scales. Further work with the wildtype intein was thus impractical and engineered Aes split intein variants resistant to aggregation were required.

### Disordered regions of the Int^N^ N_2_ and N_1_ lobes are drivers for β-sheet dependent aggregate formation

To better understand the origins of the Aes^N^ aggregation, we carried out a computational, biophysical and biochemical analysis. Sequence analysis confirmed the typical opposite charges of the Aes^N^ (120 aa) and Aes^C^ (39 aa) pieces. The charge-hydrophobicity distribution indicated a partial folding of the first lobe (N_1_) of the Aes^N^ as well as both order promoting and disorder promoting characteristics in the second lobe N_2_ of the Aes^N^ and in the Aes^C^ fragment (Supplementary Fig. 7). Circular dichroism (CD) spectroscopy and thermal shift analysis using an inactivated Aes^N^ precursor with a minimal N-extein (residues DTD) corroborated the overall only partially folded and largely disordered structure of the Aes^N^ monomer (Fig. 3A, Supplementary Fig. 8,9). Importantly, CD spectroscopy also revealed that the aggregated form of P_N_ is dominated by β-sheets. To closer pinpoint disordered regions within the Aes^N^ monomer we performed carbene-footprinting (Fig. 3B, Supplementary Fig. 10). A folded or partially folded protein is assumed to be better shielded against labeling with a reactive carbene species than a disordered region. Carbene-labeling of a monomeric Aes^N^ fraction, followed by trypsinization and quantitative ESI-MS analysis showed that three trypsin-peptides (Pep4 to Pep6) identified from the second half of the Aes^N^ sequence were more prone to labeling than the three peptides identified from the first half (Pep1 to Pep3) (Fig. 3C). Notably, however, with respect to the two symmetry-related N_1_ and N_2_ lobes, Pep4 (aa61-67) belongs to the N-terminal lobe N_1_ because the symmetry-axis between N_1_ and N_2_ is at aa68-69. All six peptides were significantly more protected against carbene-labeling in a folded Aes^N^-Aes^C^ complex, as expected (Fig. 3C, Supplementary Fig. 10).

**Figure 3.**
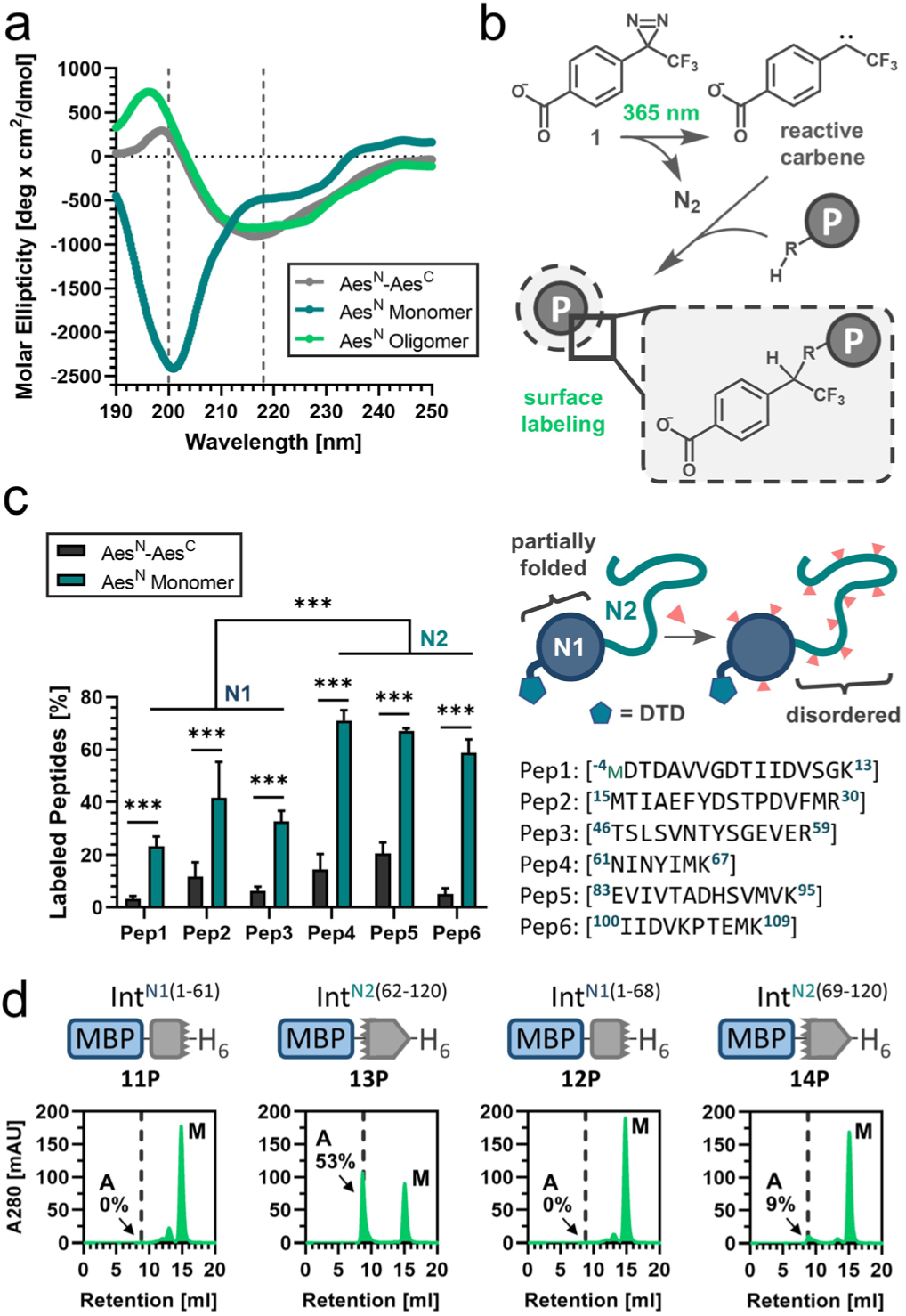
Mapping of folding states within the Aes^N^ precursor. **a** Far UV circular dichroism spectroscopy of the monomeric (blue) and the aggregated (green) species of DTD-(S1A)Aes^N^ (**8P**) in comparison to the fused complex structure MYIDTD-Aes^N^(S1A)-GSH-Aes^C^(N159A)-SVYLN (**9P**) (gray). **b** Scheme of the carbene-footprinting reaction to label exposed protein regions using trifluoromethylaryl diazirine (**1**). **c** MS-analysis of carbene labeling in Aes^N^ tryptic peptides (Pep1-Pep6) in the monomeric form of **8P** and the complex structure of MDTD-Aes^N^(S1A)-GSH-Aes^C^(N159A)-SVYLN (**10P**; 10 µM each) after 2 s of UV irradiation at 77 K using 10 mM aryldiazirine. **d** SEC-analysis of aggregate and monomer species content of protein constructs (30 µM each) containing partial Aes^N^ fragments. For panel **a**, *n* = 3-10 technical replicates. Data are presented as mean molar ellipticity corrected for concentration. For panel **c**, *n* = 3-5 technical replicates. Data are presented as mean ± s.d. normalized to the total peptide intensity. *P*-values are derived form a two-way ANOVA Sidak’s multiple comparison test (*P* = ≤0.0001).

To further investigate which parts of the Aes^N^ were the drivers for the aggregation we separately produced two N-terminal (aa1-61 (**11P**) and aa1-68 (**12P**)) and two C-terminal (aa62-120 (**13P**) and aa69-120 (**14P**)) segments according to the carbene-labeling results. We used SEC to analyze these segments as fusion proteins with MBP for their aggregation tendency (Fig. 3D).

Only **13P**, the only construct encompassing all three peptides Pep4, Pep5 and Pep6, showed a significant aggregated fraction (53%). Segment **14P**, encompassing the two C-terminal Pep5 and Pep6, showed little aggregated species that increased at higher concentration (9% at 30 µM and 15% at 150 µM; Supplementary Fig. 11). Together, not only the N_2_ lobe, but also parts of the N_1_ lobe seemed to exhibit a more disordered structure than the remainder of the Aes^N^ precursor, and these regions seemed to be responsible for the formation of soluble aggregates.

### Design of aggregation-reducing mutations that retain intein activity

We noted that the behavior of the Aes^N^ fragment was reminiscent of β-sheet-rich amyloid-like fibrils that require an at least partially disordered region with aggregation-prone sites to form the initial fibril nucleus.^38, 39^ We hypothesized that the identification of such nucleation sites within the disordered part of the Aes^N^ precursor should provide amino acid candidates for a mutational strategy to increase the monomeric and active fraction (Fig. 4A). Importantly, however, such mutations would have to maintain the association, cooperative folding and splicing activity with the Aes^C^ counterpart.

**Figure 4.**
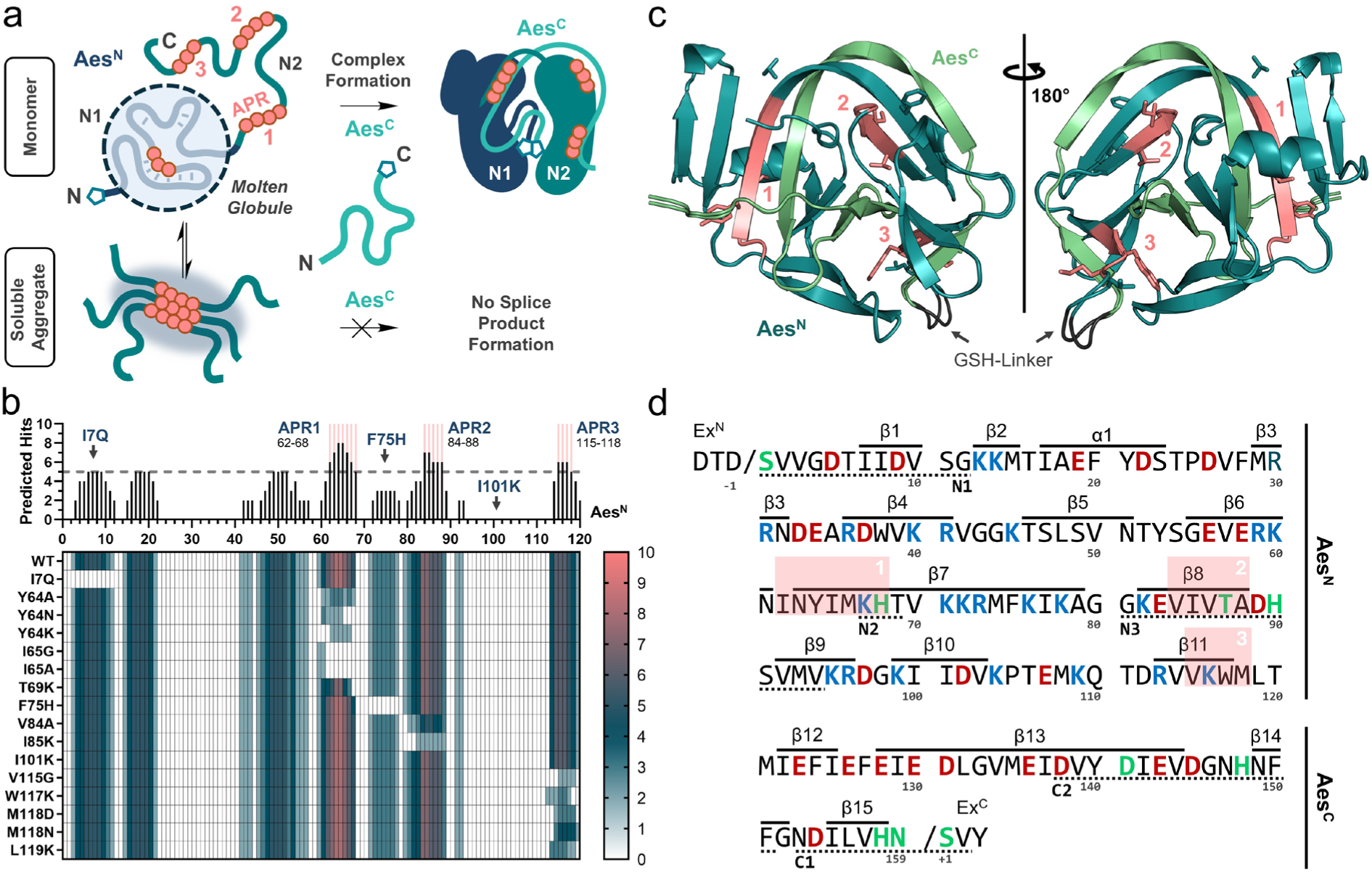
Selection of mutations for Aes^N^ engineering strategy. **a** Model of Aes^N^ aggregation. Unsaturated aggregation-prone regions (APR) in disordered regions of Aes^N^ serve as nuclei for soluble aggregate formation. **b** Sequence-based consensus prediction of amyloidogenic pattern formation in the Aes^N^ fragment using the web tool AMYLPRED2 including 10 different prediction algorithms.^40^ The upper graph indicates the number of positive hits of the different algorithms plotted against the native Aes^N^ fragment sequence. APRs labeled in red are defined by ≥5 positive hits (dashed line indicates threshold). The heat map below shows the effect of the selected single mutations on the same aggregation-prone site prediction. **c** X-ray structure of the Aes intein, obtained with the fused protein MYIDTD-Aes^N^(S1A)-GSH-Aes^C^(N159A)-SVYLN (**9P**). Shown is an overlay of chain A and B in the unit cell. (for details see Supplementary Table 7). The three strongest APRs are marked in the structure (red). The artificial GSH inker is shown in dark gray. **d** Primary sequence of the Aes^N^ (aa1-120) and the Aes^C^ fragment (aa121-139), each with 3 extein residues. Indicated are positively (red) and negatively (blue) charged residues, conserved motifs (underlined), key catalytic amino acids (green) and the location of secondary structure elements.

We used the bioinformatic web tool AMYLPRED2 to analyze the Aes^N^ sequence. This program integrates ten sequence-based prediction tools for amyloidogenic determinants that operate on different algorithms.^40^ Figure 4B shows a plot of the number of positive predictions along the Aes^N^ sequence. The more of these tools predicted the same aggregation-prone region (APR), the more probable we assumed this region to be an aggregation hotspot. Interestingly, the three strongest APRs predicted were all located in the C-terminal segment of Aes^N^ (aa60-120) consistent with our biophysical analysis.

We then designed mutations within the predicted regions of the C-terminal half of Aes^N^. To this end, we also solved the crystal structure of an Aes^N^-Aes^C^ fusion at 1.38 Å resolution to confirm that selected residues indeed formed β-sheet structures and to choose sterically fitting substitutions. See Supplementary Fig. 12 and Supplementary Note 2 for further insights from the crystal structure, which also confirmed the presence of the recently discovered motif NX histidine as a unique residue conserved in cysteine-less inteins (Supplementary Fig. 13).^13^ Interestingly, two of the three strongest APRs within Aes^N^ (APR1 & APR3) are directly in contact with Aes^C^ (Fig. 4C). Amino acids were changed to ones with a lower propensity for forming β-sheets.^41^ Mutations towards polar and in particular positively charged side chains were preferred to avoid counteracting the electrostatically driven association with the Aes^C^ precursor.

Catalytically important and highly conserved residues were not altered (Fig. 4D). This way we manually designed 14 single mutations at 10 positions. Figure 4B shows how each of these impacted the aggregation profile of the Aes^N^ sequence according to a re-analysis with AMYLPRED2.

To experimentally test these mutations, we introduced them individually into MBP-Aes^N^-H_6_ (**1P**). To exclude concentration-dependent artefacts, each mutant protein was set at 10 µM and incubated for 24 h at 15 °C prior to its analysis for aggregate formation by SEC (Fig. 5A). To our delight, the majority of the mutants showed an increased percentage of monomeric fraction, ranging from 24 to 71% compared to 19% measured for the WT (**1P**) under these conditions. Two control mutations outside the APRs of the C-terminal segment of Aes^N^, otherwise designed by similar principles (I7Q, I101K), did not have a positive influence (Fig. 4B).

**Figure 5.**
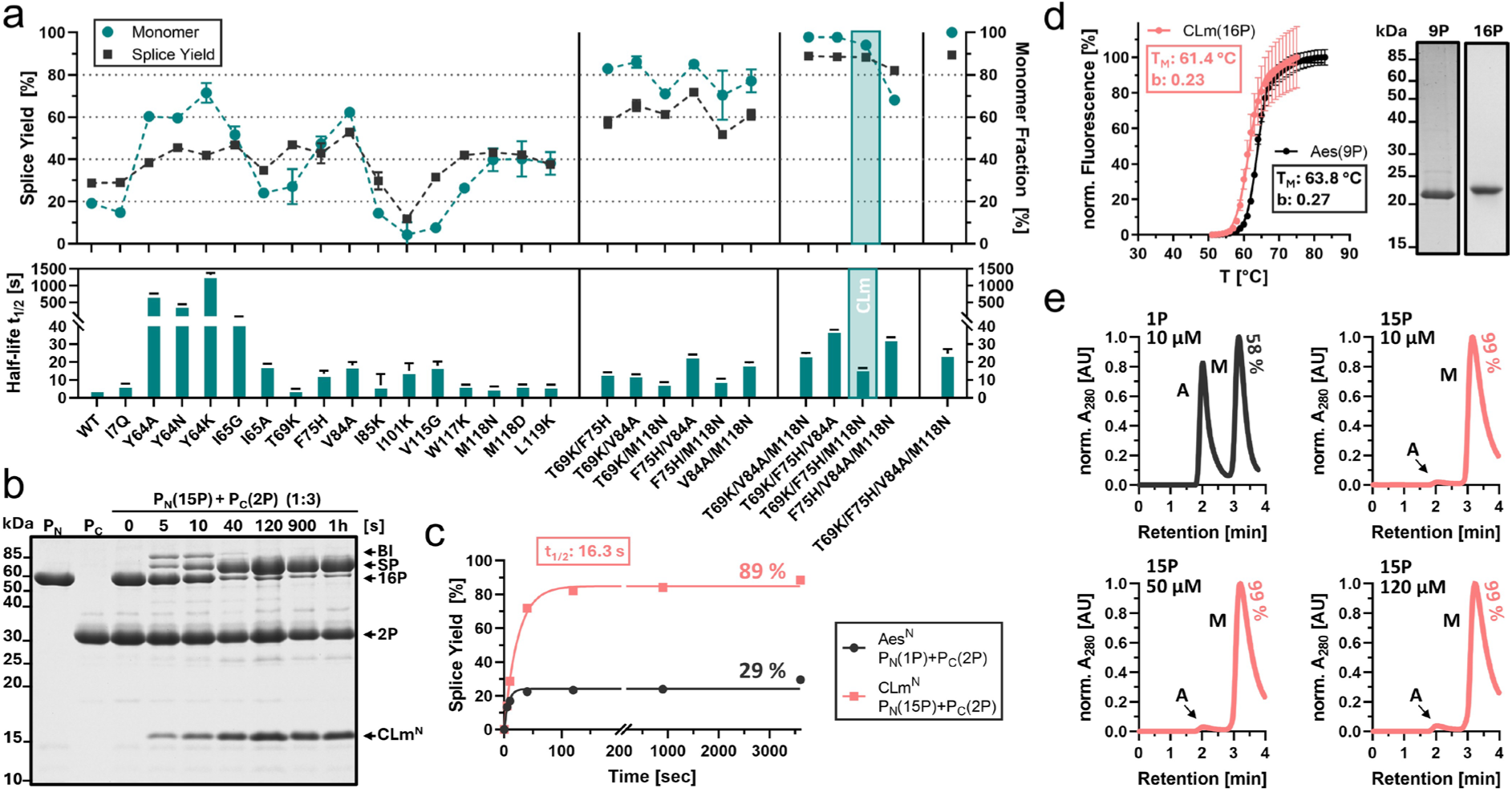
Mutational engineering of monomeric and active Aes^N^ precursors. **a** Effects of the single mutations and selected mutual combinations introduced into **1P**. The top panel shows splice yields (black) when mixed with three-fold excess of the Aes^C^ partner **2P** (30 µM) for 1 h at 37 °C (see also Supplementary Fig. 15) and the monomer proportion (blue) analyzed by SEC after incubation at 10 µM for 24h at 15 °C. The lower panel shows the determined half-life times (t_1/2_) of the PTS reaction. **b** Representative Coomassie-stained SDS-PAGE of the PTS reaction using the CLm^N^ triple mutant **15P** (T69K/F75H/M118N) with three-fold molar excess of **2P** (30 µM) at 37°C. **c** Time-courses of the PTS reactions of the native MBP-Aes^N^-H_6_ (**1P**) and the corresponding CLm^N^ mutant **15P** with three-fold molar excess of **2P** (30 µM) at 37°C. **d** Thermal stability of the fused CLm^N^-Aes^C^ complex structure **16P** (red) compared to the fused Aes^N^-Aes^C^ **9P**. Indicated are the melting temperatures (T_M_) and the Hill coefficient (b) of the folding cooperativities. The Coomassie-stained SDS-PAGE gel (right panel) shows the used constructs. **e** SEC analysis of re-aggregation of the SEC-purified monomeric Aes^N^ (**1P**) and CLm^N^ (**15P**) precursors after 24 h incubation at 15°C at the indicated concentrations. Aggregated (A) and monomeric (M) fractions are indicated. BI = branched intermediate; SP = splice product. For panel **a** and **c**, *n* = 2 technical replicates. Data are presented as normalized mean peak area ± s.d or as band intensity in SDS-PAGE analysis ± s.d. normalized to the molecular weight, respectively. For panel **d**, *n* = 6. Data are presented as mean ± s.d. normalized to the highest fluorescence intensity.

We then performed PTS assays with Aes^C^-GFP (**2P**) using all 16 mutants of **1P**, however, without SEC separation of the aggregated fraction. Figure 5A shows that indeed splice yields increased for most mutants in an overall good correlation with the increased monomeric fraction (Pearson correlation coefficient ρ = 0.89; *P* = <0.0001). Their splicing rates remained favorable and dropped at most only about 4-fold. Some mutants relatively underperformed in their splicing ability (e.g. mutations at Y64) or showed increased levels of cleavage reactions (e.g., **1P**(I65G)) and were not pursued further (Supplementary Fig. 14).

Encouraged by these findings, we created double mutants of the 4 most promising single mutations (T69K, F75H, V84A and M118N) in all six possible combinations. All double mutants showed further increased percentages of the monomeric fraction and of the splice yields, suggesting the individual mutations could act synergistically (Fig. 5A). Furthermore, in terms of splicing rates they all remained in the ultra-fast range with rate reductions of only 1.7 to 4.8-fold compared to the WT. Double mutant **1P**(T69K/M118N) showed the best performance in this regard with a t_½_ = 8.7s. We found the cleavage side reactions slightly increased but still low in most cases (up to 1% and 3% for C- and N-cleavage, respectively).

Subsequently, based on the same best single mutations, we prepared all 4 possible triple mutants of MBP-Aes^N^-H_6_ (**1P**). Impressively, these mutants formed monomeric species at >95%, except for **1P**(F75H/V84A/M118N), and they showed virtually complete splicing (about 90%). In the following, we focused on the mutant **1P**(T69K/F75H/M118N), from now on termed CLm intein (cysteine-less monomeric; **15P**), as it retained the best ultra-fast splicing activity with t_½_ = 16.3 s, only about 3.3-fold slower than the WT (Fig. 5B,C), and reached 89% splicing efficiency.

We observed 3% of C-terminal cleavage side-product, but no detectable N-terminal cleavage (Supplementary Fig. 14C). Notably, the negative impact of the T69K and F75H mutations on the thermal stability of the fused Int^N^-Int^C^ complex (Supplementary Fig. 14A) was rescued in the CLm triple mutant, as the melting temperature of the CLm^N^-Aes^C^ complex **16P** (61.4 ± 0.2 °C) was only slightly lower than for the WT Aes^N^-Aes^C^ complex **9P** (Fig. 5D). Most importantly, virtually no aggregated species could be detected any more for the CLm^N^ precursor **15P**, consistent with its high splicing efficiency. We also found that **15P** was resistant against re-aggregation over a wide concentration range (Fig. 5E). Thus, this engineered CLm^N^ precursor could be used for nearly quantitative and ultra-fast splicing without any preceding SEC separation of the monomeric fraction.

Finally, we prepared the quadruple mutant **1P**(T69K/F75H/V84A/M118N). While this mutant appeared even slightly more improved in terms of monomeric behavior and splicing yield, its rate in protein *trans*-splicing was lower (24.7 s). For these reasons, we proceeded with the new CLm mutant to explore splicing applications.

### The engineered CLm^N^ fragment shows extein generality and enhances expression yields

Unexpectedly, when testing the general utility of the CLm^N^ mutant for various exteins, we noticed a positive influence on protein expression yields. Compared to the WT Aes^N^ constructs, yields were improved 1.5 to 5.9-fold for different exteins and in different *E. coli* strains (Supplementary Fig. 16A). For example, with MBP as N-extein the yield of purified **15P** over **1P** was enhanced 1.6-fold. A construct with the ALFA nanobody^42^ (ALFAnb) as the N-extein, ALFAnb-CLm^N^-H_6_ (**17**), yielded nearly 6-fold higher levels of purified protein. Furthermore, **17P** exhibited virtually only monomeric and no aggregated species, compared to only 45% monomeric species for **18P** containing the WT Aes^N^ fragment (Supplementary Fig. 16B). Therefore, using the CLm^N^ fragment for the intein precursor increased the yield of the isolated monomeric species by 13-fold. Importantly, this construct was also fully active in PTS, as demonstrated by a ligation with H_6_-Smt3-Aes^C^-GFP (**19P**) to furnish the ALFAnb-GFP chromobody as a splice product in nearly quantitative yields and with ultra-fast kinetics of t_1/2_ = 14.3 s (Fig. 6A,B; Supplementary Fig. 16). Thus, the favorable properties of the CLm^N^ precursor fragment were found to be general also in other extein contexts.

**Figure 6.**
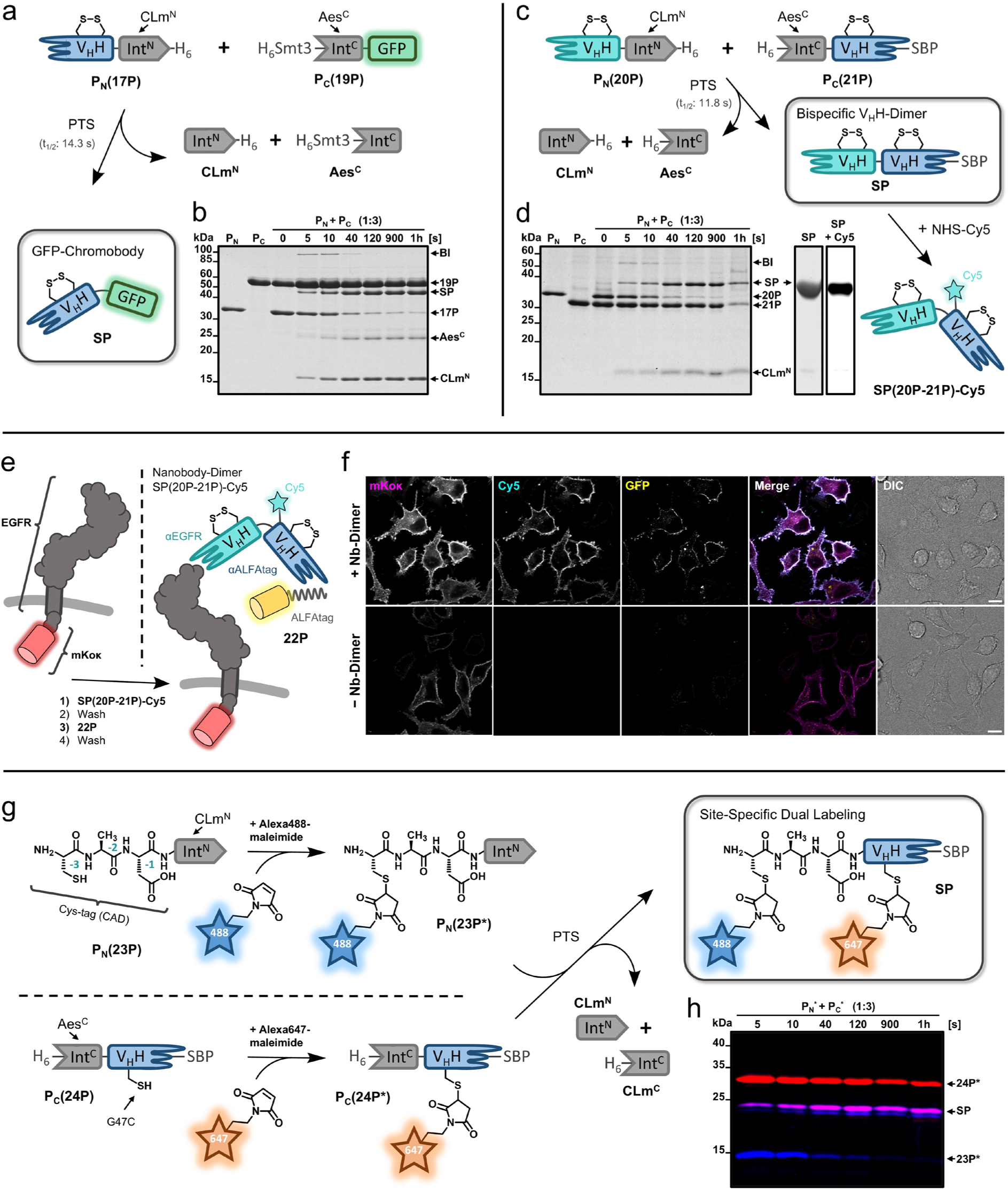
Protein modification reactions using the engineered CLm^N^ intein fragment. **a** Scheme of the ALFAnb-GFP chromobody generation by protein *trans*-splicing. **b** Analysis of the PTS reaction shown in **a**, at 37°C, using the Aes^C^ precursor at three-fold molar excess (15 µM) (see also Supplementary Fig. 16). Shown is a Coomassie-stained SDS-PAGE gel. **c** Scheme of the bispecific nanobody dimer generation by protein *trans*-splicing and its fluorophore bioconjugation. Afterwards the generated V_H_H-dimer **SP(20P-21P)** was purified and labeled with NHS-Cy5 to provide **SP(20P-21P)-Cy5**. **d** Analysis of the PTS reaction shown in **b**, at 37°C, using the Aes^C^ precursor at three-fold molar excess (15 µM) (see also Supplementary Fig. 17). Shown is a Coomassie-stained SDS-PAGE gel. The gel slices (right panel) show the purified splice product (SP) by Coomassie-staining and the Cy5-labeled SP with a fluorescence scan at 647 nm. **e** Scheme of the cell surface labeling by nanobody-dimer-mediated formation of a ternary complex. **f** Fluorescence-microscopy (CLSM) analysis of labeled cells (scale bar: 10 µm). **g** Scheme to prepare a nanobody using two separated thiol bioconjugation steps prior to protein *trans*-splicing. **h** Analysis of formation of the dually labeled ALFAnb splice product (SP). Shown is a fluorescence scan of an SDS-PAGE analysis of the PTS reaction at 37°C using the Aes^C^ precursor **24P\*** in three-fold molar excess (15 µM) (see also Supplementary Fig. 18). Calculated molecular weights are: P_N_(**17P**) = 30.7 kDa, P_C_(**19P**) = 45.0 kDa, CLm^N^-H_6_ = 14.7 kDa, H_6_-Smt3-Aes^C^ = 18.4 kDa, SP(**17P-19P**) = 42.6 kDa, P_N_(**20P**) = 30.7 kDa, P_C_(**21P**) = 26.0 kDa, H_6_-Aes^C^ = 4.7 kDa, SP(20P-21P) = 36.4 kDa, P_N_(**23P**) = 14.7 kDa, P_N_(**24P**) = 26.8 kDa, CLm^N^ = 13.7 kDa, SP(**23P*-24P\***) = 22.5 kDa. BI = branched intermediate; SP = splice product; V_H_H = variable domain heavy chain (nanobody).

### CLm split intein-mediated protein modifications that highlight its cysteine-less nature

To further demonstrate the utility of the engineered CLm mutant we aimed for the ligation of two disulfide-containing proteins (Fig. 6). Biparatopic nanobody dimers are useful reagents to dimerize proteins but their expression can be problematic and less efficient compared to single nanobodies. To generate a dimer of the anti-EGFR nanobody EgA1 and the ALFA nanobody, we prepared, purified and incubated constructs EgA1nb-CLm^N^-H_6_ (**20P**) and H_6_-CLm^C^-ALFAnb-SBP (**21P**) (Fig. 6C,D). Again, highly efficient and rapid protein *trans*-splicing occurred (t_1/2_ = 11.8 s) (Supplementary Fig. 17A). The splice product EgA1nb-ALFAnb-SBP was purified, labeled with NHS-Cy5 and tested for biparatopic binding in a cellular assay (Fig. 6E,F). To this end, HeLa cells were transiently transfected to express the extracellular and *trans*-membrane domains of EGFR, while the intracellular kinase domain was displaced with the fluorescent protein mKoκ. Cy5-labeled EgA1nb-ALFAnb-SBP specifically bound only to transfected cells. We then added ALFA-tagged GFP (**22P**, GFP-ALFAtag-H_6_) to the growth medium and observed the formation of the dimerized, three-color protein complex on the cell surface, dependent on the presence of the nanobody dimer, thus confirming its activity (Fig. 6F; Supplementary Fig. 17B). Notably, the utilized nanobody-containing split intein precursors were expressed in *E. coli* T7 shuffle cells to introduce their disulfide bonds and no reducing agents were added in the subsequent process to avoid their reduction. Both these operations would not have been suitable with cysteine-dependent split inteins.

To underline the orthogonality of the engineered CLm intein to thiol chemistries, we then aimed for site-specific dual protein labeling. We developed a new cysteine-tag^43^ reagent for minimal chemical labeling based on the engineered CLm^N^ fragment. The construct CAD-CLm^N^ (**23P**) with a minimal N-extein composed of a three-residue cysteine-tag (Cys-Ala-Asp) was obtained by cleavage of a H_6_-Smt3-CAD-CLm^N^ protein with Ulp1 protease. We bioconjugated AlexaFluor488(AF488)-maleimide to the single N-terminal cysteine (Fig. 6G; Supplementary Fig. 18). Before transferring this labeled 3aa-tag to the protein of interest by PTS, this time using the ALFA nanobody as the C-extein, we introduced a surface-exposed^42^ cysteine at position 47 to give H_6_-CLm^C^-ALFAnb(G47C)-SBP (**24P**). This single cysteine in the C-extein was bioconjugated with AF647-maleimide. Protein *trans*-splicing with both labeled intein precursor proteins then furnished the dually labeled ALFA nanobody virtually quantitatively and with ultra-fast splice kinetics (t_1/2_ = 15.5) (Fig. 6H). Of note, the second single labeled species visible below the splice product in Figure 6H is an artefact from partial thioether cleavage during sample preparation (Supplementary Fig. 18). Again, this labeling scheme could only be done with a cysteine-less split intein.

### Int^N^ precursor aggregation is widespread in split inteins

We then applied the AMYLPRED2 prediction algorithm to other intein sequences. Interestingly, high prediction scores for amyloidogenic patterns were returned for many of the commonly used split inteins (Fig. 7A; Supplementary Fig. 19). Artificially split inteins even exhibited on average a higher score for such patterns in the second segment of their Int^N^ fragment (N_2_), compared to naturally split inteins (Fig. 7B).

**Figure 7.**
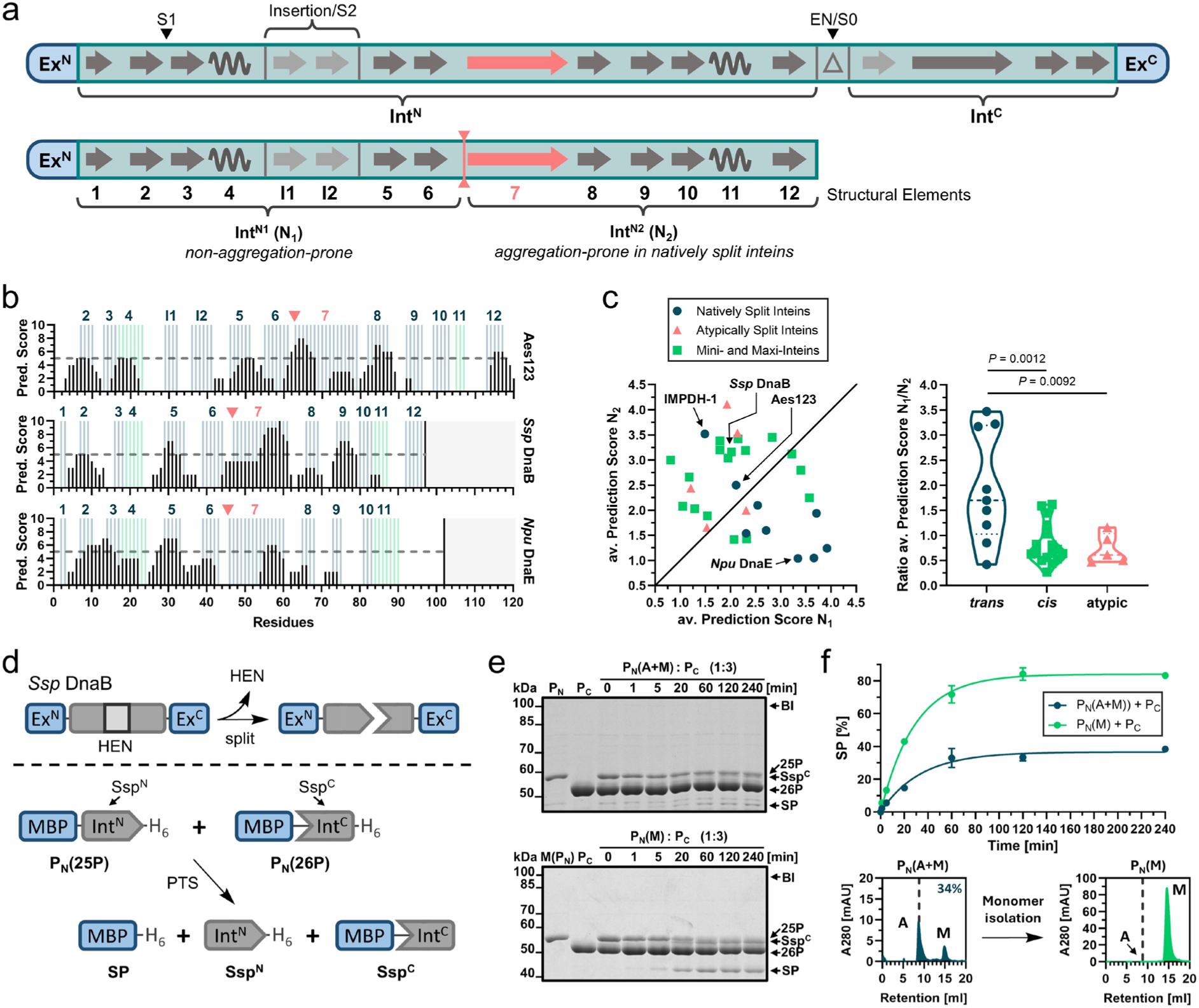
Tendency to form inactive soluble aggregates is an inherent challenge for split inteins. **a** Illustration of conserved secondary structure elements in the minimal intein horseshoe fold for *cis*-inteins (top) and the Int^N^ fragment of split inteins (bottom). Additionally, one optional insertion of β-strands I1 & I2 is shown that is also present in the Aes split intein. The split positions corresponding to typically split (EN/S0) and atypically split (S1) inteins are indicated (EN = intervening homing endonuclease domain).^23, 51^ The long β-sheet in the Int^N^ (structural element 7; marked in red) stretches across an axis of two-fold symmetry in the folded intein structure that has previously been used to define the Int^N^ N_1_ and N_2_ lobes.^36^ Here, however, the border between N_1_ and N_2_ segments was defined at the beginning of β-sheet 5, indicated by the vertical red line (see main text). **b** Sequence-based consensus predictions of amyloidogenic pattern formation using AMYLPRED2 (Ref^40^) of the Int^N^ parts of the Aes, artificially split *Ssp* DnaB and *Npu* DnaE inteins (top to bottom). Secondary structure elements are marked in blue (β-sheets) and green (α-helices). **c** Plot of the average (av.) prediction scores of the Int^N^(N_2_) segments against those of the Int^N^(N_1_) segments of commonly used inteins grouped in natively split inteins, *cis*-inteins and atypically split inteins (left panel). Distribution of the ratio of the average scores (N_1_/N_2_) for the same three groups of inteins (right panel). **d** History of the artificially split *Ssp* DnaB intein obtained from a maxi intein (top).^29, 52^ Scheme of the PTS reaction with investigated model proteins (bottom). **e** Coomassie-stained SDS-PAGE analysis of the PTS reactions (under reducing conditions) using purified proteins as illustrated in **d** using a three-fold molar excess of the *Ssp*^C^ precursor **26P** (15 µM) at 37 °C. In the upper panel the *Ssp*^N^ precursor **25P** was used without prior SEC-purification, in the lower panel the isolated monomer of **25P** was used. Plot of the PTS reactions fitted to a one-phase exponential equation. **f** SEC-analysis of **25P** before (left panel) and after (right panel) monomer isolation. These two preparations were used for the reactions in **e**. The lower graph shows the time-course of the PTS reactions in **e** fitted to a one-phase exponential equation. Calculated molecular masses are: P_N_(**25P**) = 56.4 kDa, P_C_(**26P**) = 50.1 kDa, *Ssp*^N^ = 13.3 kDa, *Ssp*^C^ = 48.6 kDa, SP = 44.6 kDa. BI = branched intermediate, SP = splice product. For panel **c**, *n* = 9 for natively split inteins, *n* = 17 for cis-inteins, *n* = 5 for atypically split inteins. *P*-values are derived from an ordinary one-way ANOVA Tukeýs multiple comparison test. For panel **f**, *n* = 2 technical replicates. Data are presented as mean ± s.d. normalized to the molecular weight.

We selected the artificially split *Ssp* DnaB intein because this intein was previously described to consume its Int^N^ precursor to only about 40 % in the protein *trans*-splicing reaction.^29, 43^ Using the model constructs MBP-Ssp^N^-H_6_ (**25P**) and MBP-Ssp^C^-H_6_ (**26P**)^29^ we could reproduce the incomplete splicing of the Int^N^ precursor (ca. 38%) (Fig. 7C,D). Indeed, analysis of **25P** by SEC revealed an inactive, high-molecular weight aggregate fraction (ca. 66%) next to a monomeric fraction (ca. 34%) (Fig. 7E). A larger quantity of the monomeric sample, prepared by our denaturation-refolding protocol, led to virtually complete splicing of the Int^N^ precursor (Fig. 7D,E). This is the first time that full activity was described for this split intein.

As a second case, we investigated the naturally split *Npu* DnaE intein, known for its rapid and efficient splicing. However, certain precursor constructs have been reported to exhibit poor splicing yields of unknown origin.^21^ We have revisited the constructs GFP-Npu^N^-H_6_ (**27P**) and Npu^C^-GFP (**28P**),^21^ which yielded only about 45% splice product formation, again due to an inactive fraction of the Int^N^ precursor **27P** (Supplementary Fig. 20). Indeed, SEC analysis revealed that this precursor existed to only 43% as a monomer, while the remaining fraction behaved as a high-molecular weight soluble aggregate. Using an isolated monomeric species of **27P** the splicing efficiency with **28P** increased to over 80%. Notably, the partially inactive **27P** was isolated at significant higher concentrations than other Npu^N^ precursors, suggesting that in this case the concentration dependency of the aggregation caused the splicing inefficiency.

## Discussion

In this work, we addressed the poor splicing efficiency of the Int^N^ precursor of the cysteine-less Aes split intein, which is a widespread and unexplained phenomenon observed for many split inteins. We carried out an in-depth biochemical and biophysical characterization of the intein combined with a structural and computational analysis. We revealed that the Aes^N^ fragment is prone to form soluble high-molecular weight aggregates with high β-sheet content that are inactive when combined with an Aes^C^ precursor, thus explaining its poor efficiency. By mutating residues in disordered regions that were computationally predicted for high propensity to serve as aggregation nuclei we could generate several triple and one quadruple mutant of the Aes^N^ fragment that showed stable monomeric behavior and nearly complete splicing activity.

Notably, the CLm^N^ triple mutant (Cysteine-Less and monomeric) combined the high splicing efficiency with a retained ultra-fast splicing rate that places it among the fastest known split inteins, about 4-fold faster than the *Npu* DnaE intein.^21^ The CLm intein (CLm^N^ + Aes^C^) thus holds great potential for expanded protein modification applications as the first cysteine-less and high-activity split intein.

Importantly, we further showed that the explanation for the incomplete splicing efficiency seems generalizable for many other split inteins. We computationally predicted similar aggregation features in several other split inteins and verified soluble aggregates to cause incompletely splicing Int^N^ precursors of the artificially split *Ssp* DnaB and the naturally split *Npu* DnaE inteins.

How can the general tendency of split inteins to form soluble aggregates be explained? Previous studies and our present work have revealed that a disordered-to-ordered transition is required to access the conserved intein horseshoe structure with its interwoven Int^N^ and Int^C^ fragments.^35, 36^ The conserved charge segregation, reflected by high and opposing local charges in the Int^N^ and Int^C^ pieces, helps both to keep the intein fragments in disordered conformations and to increase their association rate through electrostatic interactions. The disordered nature further enhances their association rate by enlarged capture radii. It also ensures the inactivity of the fragments prior to complex formation. In contrast to typical intrinsically disordered proteins,^44^ however, the disordered regions in split inteins have to exhibit a high content of β-sheet promoting residues (thus order-favoring residues) to finally stabilize the intein horseshoe fold. This propensity to form β-sheets creates an intrinsic conflict to balance out different possible folding pathways. While essential to adopt the stable horseshoe fold following association with the complementary Int^C^ precursor, it also favors the formation of β-sheet-based aggregates with itself.

In the natural cellular context, the association with the complementary Int^C^ precursor can kinetically outcompete the pathway towards aggregation. We therefore propose that the evolutionary pressure against the aggregation tendency has been insufficient for many split intein fragments. While it is challenging to predict the precise threshold at which aggregation tendency outweighs maintaining the disordered region in a folding-competent state, this notion explains the aggregation propensity of certain natural split intein precursors when produced in isolated form. It also accounts for the aggregation dependence on protein concentration, as higher concentrations promote the nucleation and elongation processes.^45, 46^

Importantly, these conclusions also align with the higher aggregation tendencies of artificially split inteins and the higher resistance to do so observed for atypically split inteins. Artificially split inteins are derived from *cis*-inteins in which the Int^N^ is linked with the Int^C^ fragment on the same polypeptide and hence was less exposed during evolution to form aggregates with itself. A similar effect is obtained by the generation of caged split inteins through fusion of parts of their complementary sequence.^47, 48^ In atypically split inteins, on the other hand, the split site is shifted to give a short Int^N^ fragment.^23, 24^ Thereby, the regions of opposing charges and disordered nature similarly become linked to a *cis*-arrangement in the large Int^C^ fragment, hence reducing the aggregation propensity by intramolecular saturation. Consequently, atypically split inteins also show an altered association mechanism that relies more on hydrophobic interactions.^49, 50^ These considerations explain why artificially generated inteins with an atypical split site showed virtually complete splicing, in contrast to their parent constructs split at the endonuclease position, but lost their favorable association and splicing kinetics.^12, 29, 34^

Finally, given that the molecular origins for the tendency of a split intein to form β-sheet aggregates are predictable from its own sequence, our rational design strategy, as demonstrated here for the Aes intein, is likely generalizable to other split inteins suffering from low splicing yields due to aggregation. Our work provides both the explanation and the cure for a folding flaw that has hampered split intein tools for more than 25 years.

## Acknowledgement

This project was financially supported by the Deutsche Forschungsgemeinschaft (grants DFG MO1073/6-2 and MO1073/9-1 to H.D.M.). We thank Dr. Jedd Bellamy-Carter (Loughborough University) for kindly providing his PepFoot software.

## Author Contributions

C. H. conceptualized the study, performed all experiments except for protein crystal structure determination and wrote the manuscript. Z. Y. performed DNA cloning and protein preparation for crystallography. K. F. performed crystallographic data collection and refinement. W. D. methodologically established carbene footprinting and supported data analysis. D. K. assisted in crystallographic data collection and refinement and reviewed the manuscript. H. D. M. conceptualized the study, supervised experiments and wrote the manuscript. All authors read and approved the final manuscript.

