## Supplementary Material for "A Cysteine-Less and Ultra-Fast Split Intein Rationally Engineered from Being Aggregation-Prone to Highly Efficient in Protein trans-Splicing"

for

### Table of Contents Page

**Supplementary Methods**


Recombinant gene expression and protein purification S4
Protein *trans*-splicing assay S4
Preparation of bispecific nanobody by protein *trans*-splicing S5
Thiol-bioconjugation of CysTag proteins S5
Densitometric analysis and determination of protein *trans*-splicing rate constants S5
Analytical RP-HPLC and ESI-MS S5
Analytical size exclusion chromatography S6
Stoke radius determination S6
Bioinformatic analysis S6
Biolayer interferometry (BLI) S7
BLI data analysis S7
Kinetical simulation S8
Circular dichroism spectroscopy S9
Thermal shift assay S9
Carbene footprinting S9
Protein crystallization and structure determination S10
Cell culture S10
Confocal laser scanning microscopy S10

#### Table of Contents Page

##### Supplementary Tables, Figures and Notes


| | | |
|---|---|---|
| **Note S1**: | Biophysical and kinetical investigation of the underlying assembly mechanism by biolayer interferometry. | S22 |
| **Note S2**: | Crystal structure of the Aes123 PolB1 intein. | S30 |

| | | |
|---|---|---|
| **Table S1**: | Reaction and rate equations of the biphasic two-step conformational change model. | S7 |
| **Table S2**: | Reaction and rate equations of the biphasic two-step conformational change model combined with a simplified one-step splice kinetic. | S8 |
| **Table S3**: | List of purified recombinant protein constructs and their expression plasmids. | S11 |
| **Table S4**: | List of plasmids used for mammalian cell culture. | S12 |
| **Table S5**: | List of sequences of recombinantly produced proteins. | S13 |
| **Table S6**: | Kinetical binding parameters of the Aes123 PolB1 intein. | S21 |
| **Table S7**: | Crystallographic data collection and structure refinement. | S28 |
| **Table S8**: | Inteins and the corresponding canonical $Int^N$ sequences used to compare the predicted aggregation tendency. | S40 |

| | | |
|---|---|---|
| **Figure S1**: | Intein sequence features and protein splicing mechanism. | S15 |
| **Figure S2**: | The C-terminal precursor is completely consumed when the N-terminal precursor is given in excess. | S16 |
| **Figure S3**: | Collective refolding of both split intein precursor proteins leads to quantitative splicing. | S17 |
| **Figure S4**: | $Aes^N$ fragment aggregation is independent of the extein context, the expression system and ionic strength. | S18 |
| **Figure S5**: | Investigation of the Aes split intein assembly mechanism by biolayer interferometry (BLI). | S19 |
| **Figure S6**: | Investigation of the underlying relationship between the Aes assembly and splice mechanism. | S20 |
| **Figure S7**: | Sequence-based prediction of the folding state by charge-hydrophobicity plot and amino acid composition profiling. | S23 |
| **Figure S8**: | Splice activity of the $Aes^N$ precursor with a short N-terminal extein of only 3 residues. | S24 |
| **Figure S9**: | Biophysical investigation of the $Aes^N$ precursor. | S25 |
| **Figure S10**: | Principle of the photoreactive carbene labeling and LC-MS based footprinting. | S26 |
| **Figure S11**: | SEC analysis of the single $Aes^N$ fragment segments fused to MBP | S27 |
| **Figure S12**: | Structural analysis of the catalytic center and the charge/hydrophobicity distribution among the intein fragments using the crystal structure of the Aes123 PolB1 intein | S29 |
| **Figure S13**: | Structural comparison between the cysteine-less Aes123 PolB1 and the PolB16 inteins. | S31 |
| **Figure S14**: | Influence of the aggregation-reducing mutations on the extent of side reactions and on the folding cooperativity to the canonical intein complex structure. | S32 |
| **Figure S15**: | Analysis of the aggregation-reducing mutants of MBP-$Aes^N$-$H_6$. | S33 |
| **Figure S16**: | The Aes123 PolB1 triple mutant increases the expression yield and protein purity. | S35 |
| **Figure S17**: | Intein-mediated generation of a bispecific nanobody-dimer for cell surface labelling. | S36 |
| **Figure S18**: | Nanobody functionalization using intein-mediated dual thiol-bioconjugation | S37 |
| **Figure S19**: | *In silico* aggregation-prone site prediction of commonly used inteins. | S38 |
| **Figure S20**: | $Int^N$ aggregation into inactive species in the natively split *Npu* DnaE intein | S41 |

**Figure S21**: SEC analysis of the purified and isolated monomeric or aggregated proteins used throughout this study. S42
**Figure S22**: Analytics to the used constructs. S43
**Figure S23**: Unprocessed SDS-PAGE images of figures shown in the main text. S45
**Figure S24**: Unprocessed SDS-PAGE images of the supplementary figures. S46

#### Supplementary Methods

**Recombinant gene expression and protein purification**. Plasmid-encoded constructs were expressed in *Escherichia coli* LOBSTR BL21 (DE3) Gold or Shuffle T7 cells grown in LB medium at 37°C or 30°C, respectively. Protein expression was induced at 28°C for 4 h or 18°C overnight by addition of either IPTG (0.4 mM) or L-arabinose (0.2% (w/v)), depending on the plasmid. Cells were ruptured using an Emulsiflex C5 (Avestin) or by sonication (10 s on/ 15 s off) for 12 min. The proteins **1P** (and its mutated variants), **4P**, **6P**, **11P** – **15P**, **17P** – **20P**, **22P**, **25P** – **26P** fused to a hexahistidine tag were purified by Ni-NTA affinity chromatography at 4°C using gravity flow columns (Cube Biotech) in Ni-NTA buffer (50 mM Tris, 300 mM NaCl, 20 mM imidazole, pH 8.0) and eluted with the same buffer containing 250 mM imidazole. Monomer isolation of **1P** and **4P** was achieved by cell rupture under denaturing conditions in Ni-NTA buffer with 8 M urea. The proteins were purified using a Ni-NTA gravity-flow column as described above and slowly refolded by stepwise dialysis into Ni-NTA buffer with 20% (w/v) sucrose as stabilizing osmolyte. The refolded protein was applied to size exclusion chromatography (SEC) performed on a Superdex 200 10/300 column (GE Healthcare) at 4°C and a flow rate of 0.75 mL/min using a FPLC Äkta Purifier system (GE Healthcare). The eluted monomeric protein was directly used for subsequent assays.

The proteins **2P**, **5P** and **28P** were expressed bearing an N-terminal $H_6$-Smt3 tag in *E. coli* LOBSTR BL21 (DE3) Gold cells grown in LB medium from an IPTG-inducible protein expression vector overnight at 18°C. The cells were lysed by sonication and purified using a Ni-NTA gravity-flow column as described above. After further purification by SEC using a Superdex 200 10/300 column (GE Healthcare) at 4°C and a flow rate of 0.75 mL/min the eluted protein was treated with 500 nM His6-tagged Ulp1 for 30 min at 8°C. After passing the protein mixture over Ni-NTA resin to remove Ulp1 and the cleaved Smt3 tag the desired product was obtained in the flowthrough.

**7P**, **8P** and **23P** were generated with a DTD or CAD tripeptide N-extein, respectively. The $H_6$-Smt3-tagged proteins were expressed and purified as described above. In the case of purification under denaturing conditions (**7P**, **8P**), the proteins were slowly refolded by stepwise dialysis in Ni-NTA buffer (50 mM Tris, 300 mM NaCl, pH 8) with 20% (w/v) sucrose as stabilizing osmolyte. The refolded protein was applied to SEC performed on a Superdex 200 10/300 column (GE Healthcare) at 4°C and a flow rate of 0.75 mL/min. The monomeric fractions were pooled and treated with 500 nM $His_6$-tagged Ulp1 for 30 min at 8°C. After passing the protein mixture over Ni-NTA resin to remove Ulp1 and the cleaved Smt3 tag the desired product was obtained in the flowthrough.

Proteins **3P**, **21P**, **24P** and **27P** bearing a streptavidin binding protein (SBP) tag were expressed in LOBSTR BL21 (DE3) Gold or Shuffle T7 cells (**24P**). Purification was achieved by a Strep-Tactin gravity-flow column (IBA) in buffer W (100 mM Tris, 150 mM NaCl, 1 mM EDTA, pH 8) and elution was induced by addition of buffer W with 2.5 mM desthiobiotin.

Tag-less protein purification of **9P** (and its mutated variants), **10P** and **16P** was achieved via chitin-binding domain (CBD) pulldown using the IMPACT™ kit (New England Biolabs) in CBD buffer (20 mM Tris, 500 mM NaCl, 1 mM EDTA, pH 8). The supernatant of the centrifuged cell lysate was transferred to a gravity flow column with chitin-agarose. On-column thiolysis was induced by adding 50 mM β-mercaptoethanol in CBD buffer to cleave of the fused *Ssp* $GyrB^N$ intein.[1] To achieve complete cleavage the column was left at 4°C shaking for 48 h. Subsequently, the eluted protein was further purified by size exclusion chromatography (SEC) using Superdex 75 10/30 column (GE Healthcare) at 4°C.

Protein concentrations were determined using the calculated extinction coefficient at 280 nm. The identity of the products was confirmed by ESI-MS and the purity was assessed by SDS-PAGE or analytical RP-HPLC.

**Protein *trans*-splicing assay**. Reactions were started by mixing the N- and C-terminal intein precursor proteins at indicated concentrations at 37°C and in absence of reducing agents (unless otherwise stated). At indicated time points aliquots were removed and the reaction was stopped by adding 4x SDS-PAGE loading buffer (500 mM Tris/HCl, 8% (w/v) SDS, 40% (v/v) glycerine, 20% (v/v) β-mercaptoethanol, 5 mg/L bromophenol blue, pH 6.8) and boiling (95°C, 5 min) or by adding 1% formic acid (final conc.). Splice product formation was analyzed by SDS-PAGE or ESI-MS, respectively.

**Preparation of bispecific nanobody by protein *trans*-splicing**. Constructs **20P** and **21P** were purified as described, mixed in Ni-NTA buffer (20 mM Tris, 300 mM NaCl, pH 8.0) and incubated at 37°C for 60 min. The reaction mixture was centrifuged (4400 rpm, 3 min), loaded onto a Ni-NTA column and the nanobody dimer was collected in the flow through. Afterwards the nanobody dimer was enriched by strep-tactin affinity purification and eluted with buffer W + 2.5 mM desthiobiotin. The purified nanobody dimer was dialyzed against PBS buffer.

**Thiol-bioconjugation of CysTag proteins**. The proteins **23P** (25 µM) and **24P** (35 µM) were purified as described above described and reduced with 5 eq. TCEP for 15 min at 4°C in PBS buffer (pH 7.4). Subsequently, Alexa Fluor 488 and Alexa Fluor 647 maleimide (Jena Bioscience), respectively, were added to each of the proteins in three steps starting with 2 eq. (60 min) followed by 1.5 eq. (30 min) twice at 4°C. The reaction with **23P** was quenched by 8 eq. DTT and purified by a Zeba™ Spin Desalting Columns (ThermoFisher). The labeled **24P** was further purified by Ni-NTA affinity chromatography.

**Densitometric analysis and determination of protein *trans*-splicing rate constants.** Coomassie-stained bands were analyzed using Gel Analyzer 2010a (gelanalyzer.com) and normalized to the corresponding molecular weight. Normalized intensities were used to calculate the ratio x of splice product to the precursor protein used in deficit to determine the splice yield as follows.

$$P(\%) = \frac{(100 \times x)}{100 + x} \tag{1}$$

To determine the overall splice rate ($k_{\text{total}}$) the splice product formation was treated as a pseudo-first-order reaction with one of the precursors given in three-fold molar excess and plotted against the time. The plot was fitted to the following singe exponential function using GraphPad Prism 8.0.

$$[P]_t = P_{max}(1 - e^{-k_{total}t}) \tag{2}$$

where P is the normalized intensity of the splice product and $k_{\text{total}}$ describes the pseudo-first-order rate equation of the protein *trans*-splice reaction. The variable t is the reaction time in seconds and $P_{\text{max}}$ is a normalization factor which represents the fraction of active precursor protein.

In order to fit the experimental data to a simplified three-state kinetic model as described by Shah *et al.*.[2] the normalized intensities of the precursor protein which was used in deficit $[\text{A}]$, the normalized intensity of the branched intermediate $[\text{BI}]$ and the normalized intensity of the splice product $[\text{P}]$ were globally fitted using GraphPad Prism 8.0 to a system of equations which are the analytical solution to the coupled differential rate equations for those species.

$$\begin{aligned}
p &= k_1 + k_2 + k_3 \\
q &= \sqrt{p^2 - 4(k_1 k_3} \\
a &= \tfrac{1}{2}(p + q) \\
b &= \tfrac{1}{2}(p - q) \\
[A]_t &= P_{max}\left[\left(\frac{k_1(a-k_3)}{a(a-b)}\right)e^{-at} + \left(\frac{k_1(k_3-b)}{b(a-b)}\right)e^{-bt}\right] \\
[BI]_t &= P_{max}\left[\left(\frac{-k_1 a}{a(a-b)}\right)e^{-at} + \left(\frac{k_1 b}{b(a-b)}\right)e^{-bt}\right] \\
[P]_t &= P_{max}\left[\left(\frac{k_1 k_3}{ab}\right) + \left(\frac{k_1 k_3}{a(a-b)}\right)e^{-at} - \left(\frac{k_1 k_3}{b(a-b)}\right)e^{-bt}\right]
\end{aligned} \tag{3}$$

In these equations, p, q, a, and b are algebraic combinations of rate constants $k_1$, $k_2$, and $k_3$. $P_{\text{max}}$ is analogous to the normalization factor described above.

**Analytical RP-HPLC and ESI-MS**. RP-HPLC analysis was done using an Agilent 1260 Infinity series system (Agilent Technologies) with a multiple wavelength detector SL and a single quadrupole mass spectrometer (Agilent). Samples were diluted with 95% $H_2O$, 5% acetonitrile and 0.1% TFA and centrifuged (14000 rpm, 2 min). According to the sample concentration, an appropriate volume was loaded on an analytical C18 column (ZORBAX SB-C18 RR HT, 3 x 50 mm, 1.8 µm, Agilent) at a flow rate of 0.4 mL/min. After a desalting step for 3 min in 5 % buffer B (eluent A: 0.1% formic acid in water; eluent B: 0.1% formic acid in acetonitrile) proteins were separated by gradual elution with 20 - 80% B in 11 min. Absorbance was recorded at 280 nm and subsequently the column was washed for 4 min with 100% B.

Mass analysis of intact proteins was performed using an UltiMate™ 3000 RS system (Thermo Fisher Scientific GmbH) connected to a maXis II UHR-qTOF mass spectrometer (Bruker Daltonik GmbH) with a standard ESI source (Apollo, Bruker Daltonik GmbH). When necessary, proteins were reduced with 2 mM TCEP at 4°C for 10 minutes to avoid inhomogeneity S15 issues. Then, samples were acidified using a 10% formic acid solution to reach a pH 2-3 and centrifuged (14000 rpm, 3 min). According to the protein concentration, an appropriate volume of the supernatant was loaded on a C4 column (Advance Bio RP-mAb C4, 2.1 mm x 50 mm, 3.5 µm, Agilent Technologies) at a flow rate of 0.6 mL/min in 5% eluent B (eluent A: 0.1% formic acid in water; eluent B: 0.1% formic acid in acetonitrile). After a desalting period of 7 minutes at 5% B, a steep gradient was applied (5-60% B in 2 min). MS settings: capillary voltage 4500 V, endplate offset 500 V, nebulizer 5.0 bar, dry gas 9.0 L/min, dry T = 200°C, mass range m/z 300-3000. Data were analyzed with DataAnalysis 4.4 (Bruker Daltonik GmbH) and deconvolution was performed using the MaxEnt algorithm implemented in the software.

**Analytical size exclusion chromatography**. Analytical gel filtration was done using a 1260 infinity LC system (Agilent) equipped with an AdvanceBio SEC 120Å 1.9 µm, 2.1 × 150 mm PEEK (Agilent), AdvanceBio SEC 200Å 1.9 µm, 2.1 × 150 mm PEEK (Agilent) or AdvanceBio SEC 200Å 1.9 µm, 4.6 × 300 mm (Agilent) column at flowrates of 0.1 and 0.35 mL/min, respectively. Prior analysis, the proteins were diluted to 10 µM and incubated at 15°C for 24 h in assay buffer (50 mM Tris, 300 mM NaCl, pH 7) to prevent the measurement of artefacts due to the concentration dependence and dynamics in aggregation. Then, the solution was directly measured at 15°C by injecting 5 µl of the sample. The obtained UV profiles at 280 nm were normalized to the highest peak to determine the aggregate/monomer ratio by the integrated peak area using GraphPad Prism 8.0.

For the time-resolved measurements to monitor the kinetics of de- and re-aggregation, the samples were directly injected from the same probe after concentration adjustment or monomer isolation (as described above) at the indicated time-points. The aggregate or monomer ratio was fitted to a hyperbolic function using GraphPad Prism 8.0. Note that the start of the re-aggregation process was better fitted by a sigmoidal function. Proteins **17P**, **18P**, **25P** and **27P** were measured without prior concentration adjustment. Here, the cysteine-containing proteins were reduced with 0.5 mM TCEP prior SEC analysis to prevent dimer formation.

**Stoke radius determination.** The Stoke radius of **1P** was determined by size exclusion chromatography on a Superdex 200 10/300 pre-packed column (GE Healthcare) at 4°C and a flow rate of 0.75 mL/min in assay buffer. The elution volume of **1P** ($V_e$) was converted into the mobility-factor parameter ($K_{av}$) using the following equation.

$$K_{av} = \frac{V_e - V_0}{V_t - V_0} \quad (4)$$

Where $V_0$ is the column void volume determined with blue dextran and $V_t$ is the total column bed volume determined with acetone at 280 nm. The Stoke radius ($R_{ST}$) of **1P** was estimated using a linear calibration plot of $R_{ST}$ vs. $(-\log K_{av})^{1/2}$, obtained with the standard globular molecular weight markers β-amylase ($R_{ST}$ = 54 Å), alcohol dehydrogenase ($R_{ST}$ = 46 Å), bovine serum albumin ($R_{ST}$ = 35 Å), carbonic anhydrase ($R_{ST}$ = 21 Å) and cytochrome c ($R_{ST}$ = 17 Å) according to the method described elsewhere.[3]

**Bioinformatic analysis**. The charge hydrophobicity plot was done using the PONDR® software (http://www.pondr.com/).[4] The mean hydrophobicity was determined using the Kyte-Doolittle hydrophobicity scale. In the plot, intrinsically disordered and native proteins are separated by a solid line which represents an empirically defined border as R = 2.785 H – 1.151, where R describes the mean net charge and H the hydrophobicity.[5] The amino acid composition of $Aes^N$ was analysed using ProtParam[6] and compared to the Disprot 9.6[7] and UniProtKB/TrEMBL 2024_04 protein database.[8]

For the prediction of aggregate-forming regions the sequence-based web tool AMYLPRED2 (http://thalis.biol.uoa.gr/AMYLPRED2/)[9] was used which employs a consensus of different methods to predict amyloid fibril formation. Note that the method AmyloidMutants usually integrated in the web tool was not used due to connection errors.

Protein structure prediction was done using Phyre2[10] and AlphaFold3[11].

**Biolayer interferometry (BLI).** Binding kinetics were measured using the Octet R8 instrument (Sartorius) and streptavidin conjugated biosensors (Sartorius). Sensor hydration was done in the final assay buffer (PBS, 0.02% Tween-20) for 10 min. Purified **5P** was biotinylated with 20 eq. biotin-X-N-hydroxy succinimide (Calbiochem) for 2h at 4°C. **6P** was diluted in assay puffer to final concentrations of 200 nM, 100 nM, 50 nM, 25 nM, 12.5 nM, 6.25 nM and 3.125 nM. The BLI assay was performed by: **(1)** sensor equilibration: sensors immersed in assay buffer for 30 s. **(2)** Ligand loading: sensor immobilization with 2 µg/mL biotinylated **5P** for 300 s. **(3)** Baseline: sensors immersed in assay buffer for 60 s. **(4)** Association: sensors immersed with the analyte **6P** at different concentrations respectively for 500 s. **(5)** Dissociation: sensors immersed in assay buffer for 500 s.

**BLI data analysis.** Due to the biphasic behavior in the binding response, the BLI data were analyzed based on analytical solutions of linear rate equations as described by Tiwari *et al.*.[12] BLI data fitting was done by double exponential functions according to the two-step conformational change model combining a bimolecular and a unimolecular equilibriums reaction using GraphPad Prism 8.0 (Supplementary Table 1). In general, the BLI response (Y) is a linear combination of the two variables

$$Y = \alpha X_1 + \beta X_2 \qquad (5)$$

where $X_1$ is $[N{\cdot}C]$ and $X_2$ is $[NC]$. Here, $[N{\cdot}C]$ describes the associated complex and $[NC]$ describes the folded complex. The association profiles were fitted using the equation:

$$Y = D + Ee^{-\sigma_1 x} + Fe^{-\sigma_2 x} \qquad (6)$$

$$D = -(E + F)$$

where D, E and F are all constants. The exponents $\sigma_1$ and $\sigma_2$ are eigenvalues and dependent on the underling matrix (see Tiwari *et al.*[12]). The dissociation profiles were fitted using the following equation with two additional parameters:

$$Y = Ee^{-\gamma_1(x-t_0)} + Fe^{-\gamma_2(x-t_0)} \qquad (7)$$

where $t_0$ is the time at the start of the dissociation phase. Since the $\alpha$ *to* $\beta$ ratio is unknown, both E and F are free fitting parameters. The exponents $\gamma_1$ and $\gamma_2$ can be obtained from the dissociation profile while $\gamma$ is $\sigma$ at the analyte concentration (C) C = 0. The dependencies of the individual fitted eigenvalues and the sums and products of these eigenvalues on the concentration C were used to verify the correct binding model as described by Tiwari *et al.*.[12]

**Supplementary Table 1**. Reaction and rate equations of the biphasic two-step conformational change model.

| *Reaction* | *Rate equation* |
|---|---|
| $[A] + [B] \underset{k_d}{\overset{k_a}{\rightleftharpoons}} [A \cdot B] \underset{k_u}{\overset{k_f}{\rightleftharpoons}} [AB]$ <br> $[B] = [B_0] - [A \cdot B] - [AB]$ | $\frac{d[A \cdot B]}{dt} = k_a C[B_0] - (k_a C - k_d - k_f)[A \cdot B] - (k_a C - k_u)[AB]$ <br> $\frac{d[AB]}{dt} = k_f[A \cdot B] - k_u[AB]$ |

To determine the rate constants, the slope of $\sigma_1+\sigma_2$ plotted against C was used to get $k_a$. The slope of $\sigma_1\sigma_2/k_a$ plotted against C gives the sum $k_f + k_u$, $\gamma_1 + \gamma_2 - (k_f + k_u)$ gives $k_d$, $\gamma_1\gamma_2/k_d$ gives $k_u$, and finally, $\gamma_1 + \gamma_2 - (k_d + k_u)$ provides $k_f$. Here, $k_a$ is the concentration dependent association rate, $k_d$ is the dissociation rate, $k_f$ describes the folding rate and $k_u$ describes the rate of unfolding. The equilibrium association constants were calculated as described below to get the overall association constant ($K_a$). Finally, the dissociation constant ($K_d$) was calculated as the inverse of $K_a$.

$$K_d = \frac{1}{K_a} \tag{8}$$

$$K_a = K_{a1}(1 + K_{a2})$$

$$K_{a1} = \frac{k_a}{k_d}$$

$$K_{a2} = \frac{k_f}{k_u}$$

The overall association and dissociation rate were calculated as described below:

$$k_{on} = \frac{k_a k_f}{k_d + k_f} \tag{9}$$

$$k_{off} = \frac{k_d k_u}{k_d + k_f}$$

For the steady state analysis, the average response (Y) from 490 to 500 s was used and the dissociation constant ($K_d$) was extracted from the following equation:

$$Y = \frac{Y_{max}\, X}{(K_D + X)} \tag{10}$$

**Kinetic simulation.** The two-step intein assembly mechanism consisting of a combined bimolecular equilibrium reaction and unimolecular equilibrium reaction was simulated by a numerical solution of the underlying rate equations (Supplementary Table 1) using the built-in function ´NDSolve´ of Wolfram Mathematica 14.1 (Wolfram Research, Inc., Mathematica, Version 14.1, Champaign, IL (2024)). Here, [A] corresponds to the analyte concentration (C) and [B] describes the ligand concentration. $[B_0]$ is the initial ligand concentration bound to the sensor. [N·C] is the concentration of the associated complex and [NC] describes the concentration of the folded complex structure. The rate equations $k_a$, $k_d$, $k_f$ and $k_u$ are described in the BLI data analysis section (see above) and were determined experimentally.

The simplified three-step intein assembly and kinetic mechanism consisting of an additional and subsequent irreversible unimolecular reaction from [AB] to the splice product [P] was similarly simulated by a numerical solution of the underlying rate equations (Supplementary Table 2).

**Supplementary Table 2**. Reaction and rate equations of the biphasic two-step conformational change model combined with a simplified one-step splice kinetic.

| *Reaction* | *Rate equation* |
|---|---|
| $[A] + [B] \underset{k_d}{\overset{k_a}{\rightleftharpoons}} [A \cdot B] \underset{k_u}{\overset{k_f}{\rightleftharpoons}} [AB] \xrightarrow{k_{splice}} [P]$ | $\frac{d[A]}{dt} = -k_a C[A][B] + k_d[A \cdot B]$ |
| | $\frac{d[B]}{dt} = -k_a C[A][B] + k_d[A \cdot B]$ |
| | $\frac{d[A \cdot B]}{dt} = k_a C[A][B] - k_d[A \cdot B] - k_f[A \cdot B] + k_u[AB]$ |
| | $\frac{d[AB]}{dt} = k_f[A \cdot B] - k_u[AB] - k_{splice}[AB]$ |
| | $\frac{d[P]}{dt} = k_{splice}[AB]$ |

In the final simulation the rate determining step of folding was equated to the overall splice rate determined by Equ. (2) ($k_f$ = $k_{total}$) to account for the competition between $k_u$ and $k_{splice}$ which is not considered in the biolayer interferometry measurements due to construct inactivation. $k_3$ was determined as described by Equ. (3) and equated with $k_{\text{splice}}$ as rate-determining step of the splice mechanism alone ($k_3$ = $k_{\text{splice}}$). To determine $k_3$ only the active precursor protein was considered to calculate the splice product formation.

**Circular dichroism spectroscopy**. The isolated proteins **9P** (40 µM) and **8P** in its monomeric (16 µM) or aggregated (23 µM) form were shortly dialyzed into CD buffer (7.5 mM $K_2HPO_4$, 2.5 mM $KH_2PO_4$, 100 mM $(NH_4)_2SO_4$, pH 7.4). Far-UV circular dichroism spectra were obtained with a Jasco J-810 apparatus equipped with a 150 W air-cooled xenon lamp using a quartz cuvette with a 0.1 mm pathlength at 25°C from 190 to 250 nm in CD buffer. Spectra were recorded at the indicated concentrations and CD buffer solution was used as blank. All measurements were repeated for at least three times. The protein denaturation experiments were performed with **8P** by adding 4 M or 6 M urea to the protein sample and CD spectra were obtained from 205 to 250 nm. The molar ellipticity was determined using the build-in spectrum analysis software.

**Thermal shift assay**. Proteins **8P** and **9P** (and its mutated variants including **16P**) were adjusted to a final concentration of 0.2 g/L and 0.5 g/L, respectively, mixed with 5x SYPRO™ Orange (Invitrogen) as fluorescent dye and added to a 96-well PCR plate. The PCR plate was sealed and placed into CFX96 Touch Real-Time PCR Detection System (Bio-Rad). Samples where heated from 10 to 90°C in increments of 1°C while fluorescence was measured. The measurements for each protein were performed 5-8 times. The data profile was cut at the highest value to account for post-peak aggregation of protein-dye complexes leading to quenching of the fluorescence signal. The truncated fluorescence imaging data was normalized and fitted to a sigmoidal four-parameter logistic equation using GraphPad Prism 8.0.

$$Y = Y_{min} + \frac{Y_{max} - Y_{min}}{1 + 10^{(T_M - X) \times b}} \quad (11)$$

where $Y_{min}$ is the minimal normalized fluorescence intensity, $Y_{max}$ is the maximal intensity, $T_M$ describes the melting temperature, and b describes the Hill coefficient.

**Carbene footprinting.** The photochemical probe sodium 4-(3-(trifluoromethyl)-3H-diazirin-3-yl)-benzoatearyldiazirine (**1**) was prepared by treating 4-(3-(trifluoromethyl)-3H-diazirin-3-yl)-benzoic acid (TCI) with sodium hydroxide as described by Manzi *et al.*.[13] Also, the photochemical labeling approach was based on Manzi *et al.*. Here, 10 µM **8P** in its monomeric form was mixed with 1 mM aryldiazirine in a total volume of 30 µL as drop placed on the lid of a microreaction tube in 50 mM Tris, 300 mM NaCl, pH 8. The mixture was left equilibrating for 5 min at RT in the dark. Then, the solution was snap-frozen in liquid $N_2$ (77 K) and placed on dry ice. The labeling reaction was initiated by 2 s of irradiation with 365 nm at 1400 mA using a LED lamp M365LPI (Thorlabs Inc.). The distance to the probe was kept constant at 5 cm. After irradiation, the sample was thawed at RT.

To prepare for typtic protein digestions, the labeled mixture was then mixed with 4x SDS loading dye (250 mM Tris/HCl, pH=6.8, 8% (w/v) SDS, 40% (v/v) glycerine, 20% (v/v) β-mercaptoethanol, 0.2% (w/v) bromophenol blue) and heated to 95°C for 5 min. Following separation by SDS-polyacrylamide gel electrophoresis and coomassie brilliant blue staining, protein bands were excised, destained by 50% (v/v) EtOH in $H_2O$, 0.1% (v/v) TFA at 60°C overnight, washed and dried with acetonitrile and by vacuum concentration. Trypsin digestion was performed at 37°C using 400 ng pre-warmed trypsin (Promega) in 50 mM ammonium carbonate supplemented with 0.01% ProteaseMax (Promega) for 6h. Afterwards, the supernatant (20 µL) was collected and acidified with 0.1% formic acid as final concnetration and analyzed by LC/MS without further dilution.

For the LC-MS analysis of the tryptic peptides an UltiMate™ 3000 RS system (Thermo Fisher Scientific GmbH, Dreieich, Germany) connected to a maXis II UHR-qTOF mass spectrometer (Bruker Daltonik GmbH, Bremen, Germany) with a standard ESI source (Apollo, Bruker Daltonik GmbH, Bremen, Germany) was used. An appropriate amount of the peptide solution was loaded on a C18 column (ZORBAX SB-C18 RR HT, 80 Å, 1.8 µm, 50 mm x 3 mm, Agilent Technologies, Waldbronn, Germany) at a flow rate of 0.6 mL/min in 5% eluent B (eluent A: 0.1% formic acid in water; eluent B: 0.1% formic acid in acetonitrile). After 5 min at 5% B, a gradient was applied (5 - 35% B in 20 min, followed by 35 - 100% B in 2 min, 1 min at 100% B and 2 min at 5% B). The following MS settings were applied: Positive polarity, capillary voltage 4500 V, endplate offset -500 V, nebulizer 1.5 bar, dry gas 8.0 L/min, dry T = 180°C, mass range m/z 150-2200.

To analyze the data of the carbene-labeled peptides, MSconvert GUI (64-bit, version 3.0.23046-c1e4e67) was used to convert Bruker file folders containing .baf files to mz5 files. Settings: Binary encoding precision 64-bit, Write index, Use zlib compression, TPP compatibility were ticked. Filter: “Zero samples” with Parameters: “removeExtra 1-“. The mz5 files were used to identify labeled and unlabeled peptides and to determine the fractional modifications by means of the PepFoot software as described elsewhere.[14] PepFoot settings: Aryldiazirin-TDBA modification of trypsin-digested peptides with a length of 5-20 amino acids and charge states of 1-8. Tolerance: 15 mmu. Miss-cleavage of peptides was not considered.

Tandem MS analysis of tryptic peptides (LC/$MS^2$) was performed using an UltiMate™ 3000 RS LC nano system (Thermo Fisher Scientific GmbH, Dreieich, Germany) connected to a maXis II UHR-qTOF mass spectrometer with a nano-ESI source (CaptiveSpray with nanoBooster, Bruker Daltonik GmbH, Bremen, Germany). 3.5 µL of each sample were loaded on a C18 trapping column (Acclaim PepMap™ 100, 5 µm, 100 Å, ID 100 µm x L 20 mm, Thermo Fisher Scientific GmbH, Dreieich, Dreieich, Germany) at a flow rate of 20 µL/min in 2% eluent B (eluent A: 0.1% formic acid in water; eluent B: 0.1% formic acid in acetonitrile). After 10 minutes of washing at 2% B, a 45-minute gradient (5 to 50% B, flow rate 500 nL/min) was applied for the separation on a C18 nano column (PepSep TWENTY-FIVE C18, ID 150 µm x L 250 mm, 1,5 µm, Bruker Daltonik GmbH, Germany). MS settings: capillary voltage 1600 V, mass range: m/z 150-2200. MS survey scans were performed with a cycle time of 2.5 s. After each survey scan, the 10 to 20 most abundant precursor ions with $z > 1$ were selected for fragmentation using collision-induced dissociation. MS/MS summation time was adjusted depending on the precursor intensity, the precursor isolation window and the collision energy were depending on the precursor m/z and charge. DataAnalysis 5.3 (Bruker Daltonik GmbH, Bremen, Germany) was used for chromatogram processing and fragment spectra isolation. The resulting mgf files were analyzed using ProteinScape 4.2 (Bruker Daltonik GmbH, Germany) as a front-end for searches against the SWISS-PROT databases on a Mascot server (Mascot 2.5, Matrix Science Ltd., London, UK).

**Protein crystallization and structure determination**. Purified **9P** was crystallized using the hanging drop method and microseeds streak seeding with a cat whisker. Best crystals were grown at 20°C with a protein concentration of 7 mg/ml in 50% precipitant mix (40% v/v PEG 500* MME; 20% w/v PEG 20000), 0.1 M buffer system (1 M Tris base, BICINE) pH 8.5, and 0.1 M monosaccharides as additive. X-ray diffraction data were collected at beamline P13 (EBML, DESY, Hamburg) to 1.38 Å resolution (Supplementary Table 7). Molecular replacement was done with a Phyre2 model[10] using the Aes123 PolB1 sequence and an initial model was obtained using Phenix AutoSol.[15] The structure was further built with Coot[16] and refined with Refmac5.[17] Four terminal N-extein residues were resolved in the electron density as well as seven residues of the C-extein. The coordinates and structure factors have been deposited in the Protein Data Bank under the accession codes 9HTH.

**Cell culture**. HeLa cells were cultured in EMEM (supplemented with 10% fetal calf serum, 1% non-essential amino acids and 1% L-glutamine) at 37°C and 5% $CO_2$. 1.8 x $10^5$ confluent cells were seeded on 16 mm coverslips in 12-well plates and used for transient transfection with the respective pDisplay plasmid, encoding EGFR-mKoκ using Metafectene (Biontex) as recommended by the manufacturer. After 36 h of incubation, binding studies and microscopy analysis were performed. To this end, cells were incubated with 250 nM of Cy5-labelled EgA1-ALFA nanobody-dimer in fresh medium for 10 min at 37°C. Afterwards, cells were incubated with 50 nM of GFP-ALFAtag-$H_6$ (**22P**) for additional 5 min, washed twice with PBS prior to fixation. Cy5-labeling of the nanobody dimer was done with 20 eq. biotin-X-N-hydroxy succinimide (Calbiochem) for 2h at 4°C, quenched with 10 mM Tris and purified by Strep-Tactin affinity chromatography before applied to the cells.

**Confocal laser scanning microscopy**. Cells were fixed with 4% paraformaldehyde in PBS for 20 min at 25°C, washed with $ddH_2O$ and mounted on coverslips using Aqua/Poly-Mount mounting solution (Polysciences). Confocal microscopy was carried out using a 63X water-immersion objective lens on a Leica DMi8 system.

#### Supplementary Tables, Figures and Notes

**Supplementary Table 3** List of purified recombinant protein constructs and their expression plasmids.

| Protein | Construct | Encoding Plasmid | Vector System | Reference |
|---|---|---|---|---|
| **1P** | MBP-YIDTD-Aes$^N$-$H_6$ | pTT04 | pMAL-c2x | Ref[18] |
| **1P**(I7Q) | MBP-Aes$^N$(I7Q)-$H_6$ | pCH96 | pMAL-c2x | this work |
| **1P**(Y64A) | MBP-Aes$^N$(Y64A)-$H_6$ | pCH141 | pMAL-c2x | this work |
| **1P**(Y64N) | MBP-Aes$^N$(Y64N)-$H_6$ | pCH127 | pMAL-c2x | this work |
| **1P**(Y64K) | MBP-Aes$^N$(Y64K)-$H_6$ | pCH126 | pMAL-c2x | this work |
| **1P**(I65G) | MBP-Aes$^N$(I65G)-$H_6$ | pCH87 | pMAL-c2x | this work |
| **1P**(I65A) | MBP-Aes$^N$(I65A)-$H_6$ | pCH95 | pMAL-c2x | this work |
| **1P**(T69K) | MBP-Aes$^N$(T69K)-$H_6$ | pCH18 | pMAL-c2x | this work |
| **1P**(F75H) | MBP-Aes$^N$(F75H)-$H_6$ | pCH128 | pMAL-c2x | this work |
| **1P**(V84A) | MBP-Aes$^N$(V84A)-$H_6$ | pCH121 | pMAL-c2x | this work |
| **1P**(I85K) | MBP-Aes$^N$(I85K)-$H_6$ | pCH125 | pMAL-c2x | this work |
| **1P**(I101K) | MBP-Aes$^N$(I101K)-$H_6$ | pCH124 | pMAL-c2x | this work |
| **1P**(V115G) | MBP-Aes$^N$(V115G)-$H_6$ | pCH90 | pMAL-c2x | this work |
| **1P**(W117K) | MBP-Aes$^N$(W117K)-$H_6$ | pCH91 | pMAL-c2x | this work |
| **1P**(M118D) | MBP-Aes$^N$(M118D)-$H_6$ | pCH92 | pMAL-c2x | this work |
| **1P**(M118N) | MBP-Aes$^N$(M118N)-$H_6$ | pCH93 | pMAL-c2x | this work |
| **1P**(L119K) | MBP-Aes$^N$(L119K)-$H_6$ | pCH94 | pMAL-c2x | this work |
| **1P**(T69K/F75H) | MBP-Aes$^N$(T69K/F75H)-$H_6$ | pCH138 | pMAL-c2x | this work |
| **1P**(T69K/V84A) | MBP-Aes$^N$(T69K/V84A)-$H_6$ | pCH137 | pMAL-c2x | this work |
| **1P**(T69K/M118N) | MBP-Aes$^N$(T69K/M118N)-$H_6$ | pCH100 | pMAL-c2x | this work |
| **1P**(F75H/V84A) | MBP-Aes$^N$(F75H/V84A)-$H_6$ | pCH134 | pMAL-c2x | this work |
| **1P**(F75H/M118N) | MBP-Aes$^N$(F75H/M118N)-$H_6$ | pCH136 | pMAL-c2x | this work |
| **1P**(V84A/M118N) | MBP-Aes$^N$(V84A/M118N)-$H_6$ | pCH135 | pMAL-c2x | this work |
| **1P**(T69K/F75H/V84A) | MBP-Aes$^N$(T69K/F75H/V84A)-$H_6$ | pCH143 | pMAL-c2x | this work |
| **1P**(T69K/F75H/M118N) (**15P**) | MBP-Aes$^N$(T69K/F75H/M118N)-$H_6$ (MBP-YIDTD-CLm$^N$-$H_6$) | pCH145 | pMAL-c2x | this work |
| **1P**(T69K/V84A/M118N) | MBP-Aes$^N$(T69K/V84A/M118N)-$H_6$ | pCH142 | pMAL-c2x | this work |
| **1P**(F75H/V84A/M118N) | MBP-Aes$^N$(F75H/V84A/M118N)-$H_6$ | pCH144 | pMAL-c2x | this work |
| **1P**(T69K/F75H/V84A/M118N) | MBP-Aes$^N$(T69K/F75H/V84A/M118N)-$H_6$ | pCH424 | pMAL-c2x | this work |
| **2P** | Aes$^C$-SVYLN-sfGFP (**Precursor**: $H_6$-smt3-Aes$^C$-SVYLN-sfGFP) | pCH196 | pET28b | this work |
| **3P** | SBP-Aes$^C$-SVYLN-SBP | pTT22 | pET16b | Ref[18] |
| **4P** | sfGFP-YIDTD-Aes$^N$-$H_6$ | pCH84 | pMAL-c2x | this work |
| **5P** | Aes$^C$(N159A)-SVYLN-sfGFP (**Precursor**: $H_6$-smt3-Aes$^C$(N159A)-SVYLN-sfGFP) | pCH188 | pET28b | this work |
| **6P** | $H_6$-Smt3-DTD-(S1A)Aes$^N$ | pCH148 | pET28b | this work |
| **7P** | DTD-Aes$^N$ (**Precursor**: $H_6$-Smt3-DTD-Aes$^N$) | pCH182 | pET28b | this work |
| **8P** | DTD-(S1A)Aes$^N$ (**Precursor**: $H_6$-Smt3-DTD-(S1A)Aes$^N$) | pCH148 | pET28b | this work |
| **9P** | MYIDTD-(S1A)-Aes$^N$-GSH-Aes$^C$(N159A)-SVYLN (**Precursor**: MYIDTD-Aes$^N$(S1A)-GSH-Aes$^C$(N159A)-SVYLN-GyrA-CBD) | pZY49 | pTWIN1 | this work |
| **9P**(I7Q) | (S1A)-Aes$^N$(I7Q)-GSH-Aes$^C$(N159A) | pCH112 | pTWIN1 | this work |
| **9P**(T69K) | (S1A)-Aes$^N$(T69K)-GSH-Aes$^C$(N159A) | pCH39 | pTWIN1 | this work |
| **9P**(F75H) | (S1A)-Aes$^N$(F75H)-GSH-Aes$^C$(N159A) | pCH140 | pTWIN1 | this work |
| **9P**(V84A) | (S1A)-Aes$^N$(V84A)-GSH-Aes$^C$(N159A) | pCH139 | pTWIN1 | this work |

| Protein | Construct | Encoding Plasmid | Vector System | Reference |
|---|---|---|---|---|
| **9P**(M118N) | (S1A)-Aes$^N$(M118N)-GSH-Aes$^C$(N159A) | pCH151 | pTWIN1 | this work |
| **10P** | MDTD-(S1A)-Aes$^N$-GSH-Aes$^C$(N159A)-SVYLN (**Precursor**: MDTD-Aes$^N$(S1A)-GSH-Aes$^C$(N159A)-SVYLN-GyrA-CBD) | pCH271 | pTWIN1 | this work |
| **11P** | MBP-(S1A)Aes$^{N1}$(S1-N61)-H$_6$ | pCH129 | pMAL-c2x | this work |
| **12P** | MBP-(S1A)Aes$^{N1}$(S1-H68)-H$_6$ | pCH118 | pMAL-c2x | this work |
| **13P** | MBP-Aes$^{N2}$(I62-T120)-H$_6$ | pCH130 | pMAL-c2x | this work |
| **14P** | MBP-Aes$^{N2}$(T69-T120)-H$_6$ | pCH117 | pMAL-c2x | this work |
| **15P** | MBP-CLm$^N$-H$_6$ | pCH145 | pMAL-c2x | this work |
| **16P** | MYIDTD-(S1A)-CLm$^N$-GSH-Aes$^C$(N159A)-SVYLN (**Precursor**: MYIDTD-CLm$^N$(S1A)-GSH-Aes$^C$(N159A)-SVYLN-GyrA-CBD) | pCH163 | pTWIN1 | this work |
| **17P** | ALFAnb-CLm$^N$-H$_6$ | pCH152 | pET16b | this work |
| **18P** | ALFAnb-Aes$^N$-H$_6$ | pCH222 | pET16b | this work |
| **19P** | H$_6$-smt3-Aes$^C$-SVYLN-sfGFP | pCH196 | pET28b | this work |
| | H$_6$-Smt3-DTD-Aes$^N$ | pCH182 | pET28b | this work |
| | EgA1nb-Aes$^N$-H$_6$ | pCH223 | pET16b | this work |
| **20P** | EgA1nb-CLm$^N$-H$_6$ | pCH200 | pET16b | this work |
| **21P** | H$_6$-Aes$^C$-ALFAnb-SBP | pCH204 | pET16b | this work |
| **22P** | sfGFP-ALFAtag-H$_6$ | pBJ278 | pET22a | Ref[19] |
| **23P** | CAD-CLm$^N$ (**Precursor**: H$_6$-smt3-CTD-Aes$^N$) | pCH291 | pET28b | this work |
| **24P** | H$_6$-Aes$^C$-ALFAnb(G47C)-SBP | pCH294 | pET16b | this work |
| **25P** | MBP-Ssp$^N$-H$_6$ | pTK56 | pTrc99a | Ref[20] |
| **26P** | MBP-Ssp$^C$-H$_6$ | pTK55 | pMAL-c2x | Ref[20] |
| **27P** | Strep-eGFP-Npu$^N$ | pVS07 | pRSFDuet1 | Ref[21] |
| **28P** | Npu$^C$-eGFP (**Precursor**: H$_6$-Smt3-Npu$^C$-eGFP) | pCH451 | pET28b | this work |

**Supplementary Table 4** List of plasmids used for mammalian cell culture.

| Plasmid | Construct | Encoding Plasmid | Vector System | Reference |
|---|---|---|---|---|
| **1** | Ig-kappa-HA-EGFR-mKoκ | pAA147 | pDisplay | this work |

**Supplementary Table 5** List of sequences of recombinantly produced proteins.

| Protein | Sequence* |
|---|---|
| **1P** | MKTEEGKLVIWINGDKGYNGLAEVGKKFEKDTGIKVTVEHPDKLEEKFPQVAATGDGPDIIFWAHDRFGGYAQSGLLAEITP<br>DKAFQDKLYPFTWDAVRYNGKLIAYPIAVEALSLIYNKDLLPNPPKTWEEIPALDKELKAKGKSALMFNLQEPYFTWPLIAA<br>DGGYAFKYENGKYDIKDVGVDNAGAKAGLTFLVDLIKNKHMNADTDYSIAEAAFNKGETAMTINGPWAWSNIDTSKVNYG<br>VTVLPTFKGQPSKPFVGVLSAGINAASPNKELAKEFLENYLLTDEGLEAVNKDKPLGAVALKSYEEELAKDPRIAATMENAQK<br>GEIMPNIPQMSAFWYAVRTAVINAASGRQTVDEALKDAQTNSSSNNNNNNNNNNLGIEGRISEFYIDTDSVVGDTIIDVSGKK<br>MTIAEFYDSTPDVFMRRNDEARDWVKRVGGKTSLSVNTYSGEVERKNINYIMKHTVKKRMFKIKAGGKEVIVTADHSVMVK<br>RDGKIIDVKPTEMKQTDRVVKWMLTGSHHHHHH |
| **2P/19P** | MGSSHHHHHHSSGLVPRGSHMASMSDSEVNQEAKPEVKPEVKPETHINLKVSDGSSEIFFKIKKTTPLRRLMEAFAKRQGKE<br>MDSLRFLYDGIRIQADQTPEDLDMEDNDIIEAHREQIGGSMIEFIEFEIEDLGVMEIDVYDIEVDGNHNFFGNDILVHNSVYLN<br>GTSKGEELFTGVVPILVELDGDVNGHKFSVRGEGEGDATNGKLTLKFICTTGKLPVPWPTLVTTLTYGVQCFSRYPDHMKRH<br>DFFKSAMPEGYVQERTISFKDDGTYKTRAEVKFEGDTLVNRIELKGIDFKEDGNILGHKLEYNFNSHNVYITADKQKNGIKAN<br>FKIRHNVEDGSVQLADHYQQNTPIGDGPVLLPDNHYLSTQSVLSKDPNEKRDHMVLLEFVTAAGITHG |
| **3P** | MDEKTTGWRGGHVVEGLAGELEQLRARLEHHPQGQREPGASGGGGSSSMIEFIEFEIEDLGVMEIDVYDIEVDGNHNFFGND<br>ILVHNSVYLNGTMDEKTTGWRGGHVVEGLAGELEQLRARLEHHPQGQREP |
| **4P** | MSKGEELFTGVVPILVELDGDVNGHKFSVRGEGEGDATNGKLTLKFICTTGKLPVPWPTLVTTLTYGVQCFSRYPDHMKRHD<br>FFKSAMPEGYVQERTISFKDDGTYKTRAEVKFEGDTLVNRIELKGIDFKEDGNILGHKLEYNFNSHNVYITADKQKNGIKANF<br>KIRHNVEDGSVQLADHYQQNTPIGDGPVLLPDNHYLSTQSVLSKDPNEKRDHMVLLEFVTAAGITHGGEFYIDTDSVVGDTII<br>DVSGKKMTIAEFYDSTPDVFMRRNDEARDWVKRVGGKTSLSVNTYSGEVERKNINYIMKHTVKKRMFKIKAGGKEVIVTAD<br>HSVMVKRDGKIIDVKPTEMKQTDRVVKWMLTGSHHHHHH |
| **5P** | MGSSHHHHHHSSGLVPRGSHMASMSDSEVNQEAKPEVKPEVKPETHINLKVSDGSSEIFFKIKKTTPLRRLMEAFAKRQGKE<br>MDSLRFLYDGIRIQADQTPEDLDMEDNDIIEAHREQIGGSRIEFIEFEIEDLGVMEIDVYDIEVDGNHNFFGNDILVHASVYLNG<br>TSKGEELFTGVVPILVELDGDVNGHKFSVRGEGEGDATNGKLTLKFICTTGKLPVPWPTLVTTLTYGVQCFSRYPDHMKRHD<br>FFKSAMPEGYVQERTISFKDDGTYKTRAEVKFEGDTLVNRIELKGIDFKEDGNILGHKLEYNFNSHNVYITADKQKNGIKANF<br>KIRHNVEDGSVQLADHYQQNTPIGDGPVLLPDNHYLSTQSVLSKDPNEKRDHMVLLEFVTAAGITHG |
| **6P/8P** | MGSSHHHHHHSSGLVPRGSHMASMSDSEVNQEAKPEVKPEVKPETHINLKVSDGSSEIFFKIKKTTPLRRLMEAFAKRQGKE<br>MDSLRFLYDGIRIQADQTPEDLDMEDNDIIEAHREQIGGDTDAVVGDTIIDVSGKKMTIAEFYDSTPDVFMRRNDEARDWVK<br>RVGGKTSLSVNTYSGEVERKNINYIMKHTVKKRMFKIKAGGKEVIVTADHSVMVKRDGKIIDVKPTEMKQTDRVVKWMLT |
| **7P** | MGSSHHHHHHSSGLVPRGSHMASMSDSEVNQEAKPEVKPEVKPETHINLKVSDGSSEIFFKIKKTTPLRRLMEAFAKRQGKE<br>MDSLRFLYDGIRIQADQTPEDLDMEDNDIIEAHREQIGGDTDSVVGDTIIDVSGKKMTIAEFYDSTPDVFMRRNDEARDWVK<br>RVGGKTSLSVNTYSGEVERKNINYIMKHTVKKRMFKIKAGGKEVIVTADHSVMVKRDGKIIDVKPTEMKQTDRVVKWMLT |
| **9P** | MYIDTDAVVGDTIIDVSGKKMTIAEFYDSTPDVFMRRNDEARDWVKRVGGKTSLSVNTYSGEVERKNINYIMKHTVKKRMF<br>KIKAGGKEVIVTADHSVMVKRDGKIIDVKPTEMKQTDRVVKWMLTGSHMIEFIEFEIEDLGVMEIDVYDIEVDGNHNFFGNDI<br>LVHASVYLNKLGGCFSGDTLVALTDGRSVSFEQLVEEEKQGKQNFCYTIRHDGSIGVEKIINARKTKTNAKVIKVTLDNGESIIC<br>TPDHKFMLRDGSYKCAMDLTLDDSLMPLHRKISTTEDSGHMEAVLNYNHRIVNIEAVSETIDVYDIEVPHTHNFALASTGMK<br>IEEGKLTNPGVSAWQVNTAYTAGQLVTYNGKTYKCLQPHTSLAGWEPSNVPALWQLQ |
| **10P** | MDTDAVVGDTIIDVSGKKMTIAEFYDSTPDVFMRRNDEARDWVKRVGGKTSLSVNTYSGEVERKNINYIMKHTVKKRMFKI<br>KAGGKEVIVTADHSVMVKRDGKIIDVKPTEMKQTDRVVKWMLTGSHMIEFIEFEIEDLGVMEIDVYDIEVDGNHNFFGNDIL<br>VHASVYLNKLGGCFSGDTLVALTDGRSVSFEQLVEEEKQGKQNFCYTIRHDGSIGVEKIINARKTKTNAKVIKVTLDNGESIICT<br>PDHKFMLRDGSYKCAMDLTLDDSLMPLHRKISTTEDSGHMEAVLNYNHRIVNIEAVSETIDVYDIEVPHTHNFALASTGMKI<br>EEGKLTNPGVSAWQVNTAYTAGQLVTYNGKTYKCLQPHTSLAGWEPSNVPALWQLQ |
| **11P** | MKTEEGKLVIWINGDKGYNGLAEVGKKFEKDTGIKVTVEHPDKLEEKFPQVAATGDGPDIIFWAHDRFGGYAQSGLLAEITP<br>DKAFQDKLYPFTWDAVRYNGKLIAYPIAVEALSLIYNKDLLPNPPKTWEEIPALDKELKAKGKSALMFNLQEPYFTWPLIAA<br>DGGYAFKYENGKYDIKDVGVDNAGAKAGLTFLVDLIKNKHMNADTDYSIAEAAFNKGETAMTINGPWAWSNIDTSKVNYG<br>VTVLPTFKGQPSKPFVGVLSAGINAASPNKELAKEFLENYLLTDEGLEAVNKDKPLGAVALKSYEEELAKDPRIAATMENAQK<br>GEIMPNIPQMSAFWYAVRTAVINAASGRQTVDEALKDAQTNSSSNNNNNNNNNNLGIEGRGENLYFQGYIDTDAVVGDTIID<br>VSGKKMTIAEFYDSTPDVFMRRNDEARDWVKRVGGKTSLSVNTYSGEVERKNGSHHHHHH |
| **12P** | MKTEEGKLVIWINGDKGYNGLAEVGKKFEKDTGIKVTVEHPDKLEEKFPQVAATGDGPDIIFWAHDRFGGYAQSGLLAEITP<br>DKAFQDKLYPFTWDAVRYNGKLIAYPIAVEALSLIYNKDLLPNPPKTWEEIPALDKELKAKGKSALMFNLQEPYFTWPLIAA<br>DGGYAFKYENGKYDIKDVGVDNAGAKAGLTFLVDLIKNKHMNADTDYSIAEAAFNKGETAMTINGPWAWSNIDTSKVNYG<br>VTVLPTFKGQPSKPFVGVLSAGINAASPNKELAKEFLENYLLTDEGLEAVNKDKPLGAVALKSYEEELAKDPRIAATMENAQK<br>GEIMPNIPQMSAFWYAVRTAVINAASGRQTVDEALKDAQTNSSSNNNNNNNNNNLGIEGRGENLYFQGYIDTDAVVGDTIID<br>VSGKKMTIAEFYDSTPDVFMRRNDEARDWVKRVGGKTSLSVNTYSGEVERKNINYIMKHGSHHHHHH |
| **13P** | MKTEEGKLVIWINGDKGYNGLAEVGKKFEKDTGIKVTVEHPDKLEEKFPQVAATGDGPDIIFWAHDRFGGYAQSGLLAEITP<br>DKAFQDKLYPFTWDAVRYNGKLIAYPIAVEALSLIYNKDLLPNPPKTWEEIPALDKELKAKGKSALMFNLQEPYFTWPLIAA<br>DGGYAFKYENGKYDIKDVGVDNAGAKAGLTFLVDLIKNKHMNADTDYSIAEAAFNKGETAMTINGPWAWSNIDTSKVNYG<br>VTVLPTFKGQPSKPFVGVLSAGINAASPNKELAKEFLENYLLTDEGLEAVNKDKPLGAVALKSYEEELAKDPRIAATMENAQK<br>GEIMPNIPQMSAFWYAVRTAVINAASGRQTVDEALKDAQTNSSSNNNNNNNNNNLGIEGRGENLYFQGINYIMKHTVKKRM<br>FKIKAGGKEVIVTADHSVMVKRDGKIIDVKPTEMKQTDRVVKWMLTGSHHHHHH |
| **14P** | MKTEEGKLVIWINGDKGYNGLAEVGKKFEKDTGIKVTVEHPDKLEEKFPQVAATGDGPDIIFWAHDRFGGYAQSGLLAEITP<br>DKAFQDKLYPFTWDAVRYNGKLIAYPIAVEALSLIYNKDLLPNPPKTWEEIPALDKELKAKGKSALMFNLQEPYFTWPLIAA<br>DGGYAFKYENGKYDIKDVGVDNAGAKAGLTFLVDLIKNKHMNADTDYSIAEAAFNKGETAMTINGPWAWSNIDTSKVNYG<br>VTVLPTFKGQPSKPFVGVLSAGINAASPNKELAKEFLENYLLTDEGLEAVNKDKPLGAVALKSYEEELAKDPRIAATMENAQK<br>GEIMPNIPQMSAFWYAVRTAVINAASGRQTVDEALKDAQTNSSSNNNNNNNNNNLGIEGRGENLYFQGTVKKRMFKIKAGG<br>KEVIVTADHSVMVKRDGKIIDVKPTEMKQTDRVVKWMLTGSHHHHHH |

| Protein | Sequence* |
|---|---|
| **15P** | MKTEEGKLVIWINGDKGYNGLAEVGKKFEKDTGIKVTVEHPDKLEEKFPQVAATGDGPDIIFWAHDRFGGYAQSGLLAEITP<br>DKAFQDKLYPFTWDAVRYNGKLIAYPIAVEALSLIYNKDLLPNPPKTWEEIPALDKELKAKGKSALMFNLQEPYFTWPLIAA<br>DGGYAFKYENGKYDIKDVGVDNAGAKAGLTFLVDLIKNKHMNADTDYSIAEAAFNKGETAMTINGPWAWSNIDTSKVNYG<br>VTVLPTFKGQPSKPFVGVLSAGINAASPNKELAKEFLENYLLTDEGLEAVNKDKPLGAVALKSYEEELAKDPRIAATMENAQK<br>GEIMPNIPQMSAFWYAVRTAVINAASGRQTVDEALKDAQTNSSSNNNNNNNNNNLGIEGRISEFYIDTDSVVGDTIIDVSGKK<br>MTIAEFYDSTPDVFMRRNDEARDWVKRVGGKTSLSVNTYSGEVERKNINYIMKHKVKKRMHKIKAGGKEVIVTADHSVMV<br>KRDGKIIDVKPTEMKQTDRVVKWNLTGSHHHHHH |
| **16P** | MYIDTDAVVGDTIIDVSGKKMTIAEFYDSTPDVFMRRNDEARDWVKRVGGKTSLSVNTYSGEVERKNINYIMKHKVKKRMH<br>KIKAGGKEVIVTADHSVMVKRDGKIIDVKPTEMKQTDRVVKWNLTGSHMIEFIEFEIEDLGVMEIDVYDIEVDGNHNFFGNDI<br>LVHASVYLNKLGGCFSGDTLVALTDGRSVSFEQLVEEEKQGKQNFCYTIRHDGSIGVEKIINARKTKTNAKVIKVTLDNGESIIC<br>TPDHKFMLRDGSYKCAMDLTLDDSLMPLHRKISTTEDSGHMEAVLNYNHRIVNIEAVSETIDVYDIEVPHTHNFALASTGMK<br>IEEGKLTNPGVSAWQVNTAYTAGQLVTYNGKTYKCLQPHTSLAGWEPSNVPALWQLQ |
| **17P** | MSGEVQLQESGGGLVQPGGSLRLSCTASGVTISALNAMAMGWYRQAPGERRVMVAAVSERGNAMYRESVQGRFTVTRDFT<br>NKMVSLQMDNLKPEDTAVYYCHVLEDRVDSFHDYWGQGTQVTVSSEPKTPKPQTGGSGTGSGSGYIDTDSVVGDTIIDVSGK<br>KMTIAEFYDSTPDVFMRRNDEARDWVKRVGGKTSLSVNTYSGEVERKNINYIMKHTVKKRMFKIKAGGKEVIVTADHSVMV<br>KRDGKIIDVKPTEMKQTDRVVKWMLTGSHHHHHH |
| **18P** | MSGEVQLQESGGGLVQPGGSLRLSCTASGVTISALNAMAMGWYRQAPGERRVMVAAVSERGNAMYRESVQGRFTVTRDFT<br>NKMVSLQMDNLKPEDTAVYYCHVLEDRVDSFHDYWGQGTQVTVSSEPKTPKPQTGGSGTGSGSGYIDTDSVVGDTIIDVSGK<br>KMTIAEFYDSTPDVFMRRNDEARDWVKRVGGKTSLSVNTYSGEVERKNINYIMKHKVKKRMHKIKAGGKEVIVTADHSVM<br>VKRDGKIIDVKPTEMKQTDRVVKWNLTGSHHHHHH |
| **20P** | MGQVQLQESGGGLVQPGGSLRLSCAASGRTFSSYAMGWFRQAPGKQREFVAAIRWSGGYTYYTDSVKGRFTISRDNAKTTVY<br>LQMNSLKPEDTAVYYCAATYLSSDYSRYALPQRPLDYDYWGQGTQVTVSSLEGTGSGSGYIDTDSVVGDTIIDVSGKKMTIAE<br>FYDSTPDVFMRRNDEARDWVKRVGGKTSLSVNTYSGEVERKNINYIMKHKVKKRMHKIKAGGKEVIVTADHSVMVKRDGKI<br>IDVKPTEMKQTDRVVKWNLTGSHHHHHH |
| **21P** | MGHHHHHHMIEFIEFEIEDLGVMEIDVYDIEVDGNHNFFGNDILVHNSVYLNGSGSHMSGEVQLQESGGGLVQPGGSLRLSCT<br>ASGVTISALNAMAMGWYRQAPGERRVMVAAVSERGNAMYRESVQGRFTVTRDFTNKMVSLQMDNLKPEDTAVYYCHVLE<br>DRVDSFHDYWGQGTQVTVSSEPKTPKPQTGGSLEMDEKTTGWRGGHVVEGLAGELEQLRARLEHHPQGQREP |
| **22P** | MSKGEELFTGVVPILVELDGDVNGHKFSVRGEGEGDATNGKLTLKFICTTGKLPVPWPTLVTTLTYGVQCFSRYPDHMKRHD<br>FFKSAMPEGYVQERTISFKDDGTYKTRAEVKFEGDTLVNRIELKGIDFKEDGNILGHKLEYNFNSHNVYITADKQKNGIKANF<br>KIRHNVEDGSVQLADHYQQNTPIGDGPVLLPDNHYLSTQSVLSKDPNEKRDHMVLLEFVTAAGITHGGSGGGSSRLEEELRR<br>RLTEGGGSLEHHHHHH |
| **23P** | MGSSHHHHHHSSGLVPRGSHMASMSDSEVNQEAKPEVKPEVKPETHINLKVSDGSSEIFFKIKKTTPLRRLMEAFAKRQGKE<br>MDSLRFLYDGIRIQADQTPEDLDMEDNDIIEAHREQIGGCADSVVGDTIIDVSGKKMTIAEFYDSTPDVFMRRNDEARDWVK<br>RVGGKTSLSVNTYSGEVERKNINYIMKHKVKKRMHKIKAGGKEVIVTADHSVMVKRDGKIIDVKPTEMKQTDRVVKWNLT |
| **24P** | MGHHHHHHMIEFIEFEIEDLGVMEIDVYDIEVDGNHNFFGNDILVHNSVYLNGSGSHMSGEVQLQESGGGLVQPGGSLRLSCT<br>ASGVTISALNAMAMGWYRQAPCERRVMVAAVSERGNAMYRESVQGRFTVTRDFTNKMVSLQMDNLKPEDTAVYYCHVLE<br>DRVDSFHDYWGQGTQVTVSSEPKTPKPQTGGSLEMDEKTTGWRGGHVVEGLAGELEQLRARLEHHPQGQREP |
| **25P** | MEIEEGKLVIWINGDKGYNGLAEVGKKFEKDTGIKVTVEHPDKLEEKFPQVAATGDGPDIIFWAHDRFGGYAQSGLLAEITPD<br>KAFQDKLYPFTWDAVRYNGKLIAYPIAVEALSLIYNKDLLPNPPKTWEEIPALDKELKAKGKSALMFNLQEPYFTWPLIAAD<br>GGYAFKYENGKYDIKDVGVDNAGAKAGLTFLVDLIKNKHMNADTDYSIAEAAFNKGETAMTINGPWAWSNIDTSKVNYGVT<br>VLPTFKGQPSKPFVGVLSAGINAASPNKELAKEFLENYLLTDEGLEAVNKDKPLGAVALKSYEEELAKDPRIAATMENAQKGE<br>IMPNIPQMSAFWYAVRTAVINAASGRQTVDEALKDAQTNSSSNNNNNNNNNNLGIEGRISEFSGCISGDSLISLASTGKRVSIK<br>DLLDEKDFEIWAINEQTMKLESAKVSRVFCTGKKLVYILKTRLGRTIKATANHRFLTIDGWKRLDELSLKEHIALPRKLESSSL<br>SSYGSRSHHHHHH |
| **26P** | MKIEEGKLVIWINGDKGYNGLAEVGKKFEKDTGIKVTVEHPDKLEEKFPQVAATGDGPDIIFWAHDRFGGYAQSGLLAEITPD<br>KAFQDKLYPFTWDAVRYNGKLIAYPIAVEALSLIYNKDLLPNPPKTWEEIPALDKELKAKGKSALMFNLQEPYFTWPLIAAD<br>GGYAFKYENGKYDIKDVGVDNAGAKAGLTFLVDLIKNKHMNADTDYSIAEAAFNKGETAMTINGPWAWSNIDTSKVNYGVT<br>VLPTFKGQPSKPFVGVLSAGINAASPNKELAKEFLENYLLTDEGLEAVNKDKPLGAVALKSYEEELAKDPRIAATMENAQKGE<br>IMPNIPQMSAFWYAVRTAVINAASGRQTVDEALKDAQTNSSSNNNNNNNNNNLGIEGRISEFGSSSSPEIEKLSQSDIYWDSI<br>VSITETGVEEVFDLTVPGPHNFVANDIIVHNSIRSRSHHHHHH |
| **27P** | MASWSHPQFEKASGTVSKGEELFTGVVPILVELDGDVNGHKFSVSGEGEGDATYGKLTLKFICTTGKLPVPWPTLVTTLTYG<br>VQCFSRYPDHMKQHDFFKSAMPEGYVQERTIFFKDDGNYKTRAEVKFEGDTLVNRIELKGIDFKEDGNILGHKLEYNYNSHN<br>VYIMADKQKNGIKVNFKIRHNIEDGSVQLADHYQQNTPIGDGPVLLPDNHYLSTQSALSKDPNEKRDHMVLLEFVTAAGITL<br>GMDELYKGSRSGSCLSYETEILTVEYGLLPIGKIVEKRIECTVYSVDNNGNIYTQPVAQWHDRGEQEVFEYCLEDGSLIRATKD<br>HKFMTVDGQMLPIDEIFERELDLMRVDNLPN |
| **28P** | MGSSHHHHHHSSGLVPRGSHMASMSDSEVNQEAKPEVKPEVKPETHINLKVSDGSSEIFFKIKKTTPLRRLMEAFAKRQGKE<br>MDSLRFLYDGIRIQADQTPEDLDMEDNDIIEAHREQIGGMIKIATRKYLGKQNVYDIGVERDHNFALKNGFIASNCFNGTVSK<br>GEELFTGVVPILVELDGDVNGHKFSVSGEGEGDATYGKLTLKFICTTGKLPVPWPTLVTTLTYGVQCFSRYPDHMKQHDFFKS<br>AMPEGYVQERTIFFKDDGNYKTRAEVKFEGDTLVNRIELKGIDFKEDGNILGHKLEYNYNSHNVYIMADKQKNGIKVNFKIR<br>HNIEDGSVQLADHYQQNTPIGDGPVLLPDNHYLSTQSALSKDPNEKRDHMVLLEFVTAAGITLGMDELYK |

* The marked sequences either indicate the N-terminal proteolytically cleavable $H_6$-Smt3 tag or the C-terminal chemically cleavable GyrA-CBD tag.

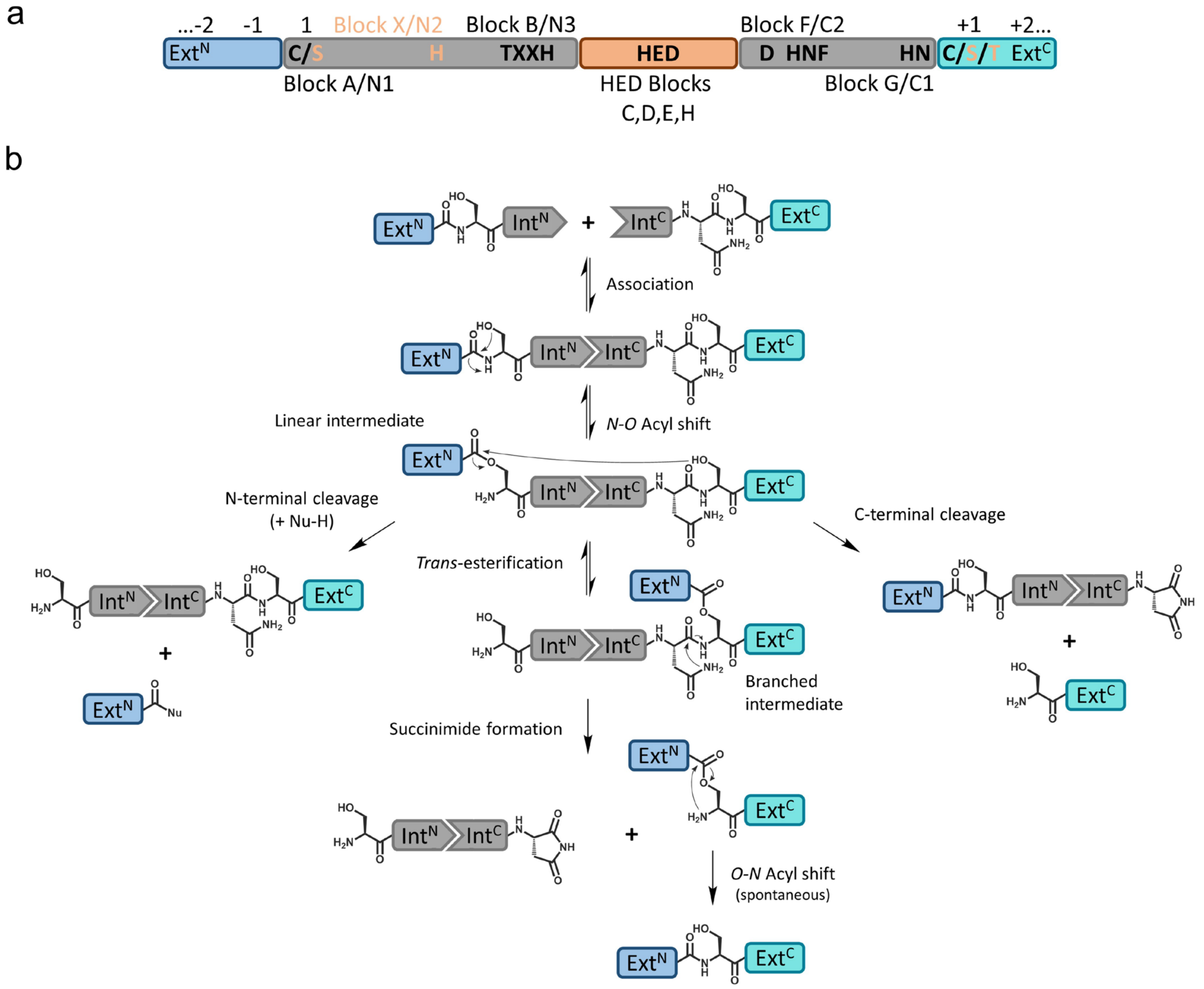


**Supplementary Figure 1** Intein sequence features and protein splicing mechanism. **a** Conserved sequence motifs of class 1 inteins relative to the intervening homing endonuclease domain. **b** General protein splicing mechanism of cysteine-less split inteins. Note that cysteine-less inteins operate with Ser1 and Ser+1 (Thr+1) residues at the two splice junctions to form the linear and branched ester intermediates.

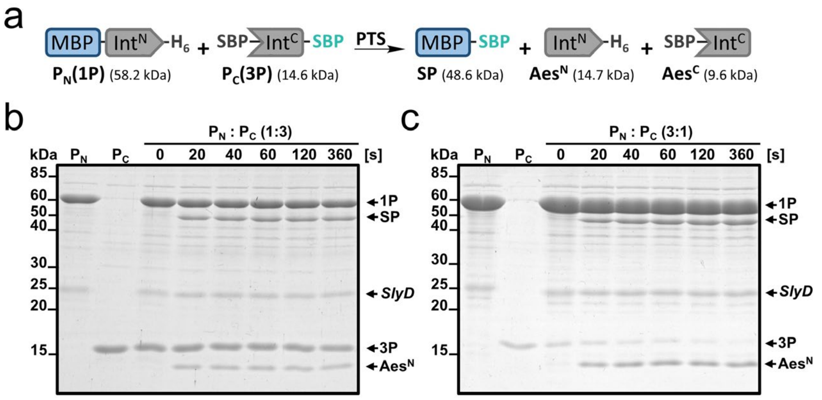


**Supplementary Figure 2** The C-terminal precursor is completely consumed when the N-terminal precursor is given in excess (additional data to Fig. 1d). **a** Scheme of the PTS reaction. **b** SDS-PAGE analysis of the PTS reaction using **1P** with **3P** in three-fold molar excess (10 µM to 30 µM at 37°C). **c** SDS-PAGE analysis of the PTS reaction using **1P** in three-fold molar excess to **3P** (30 µM to 10 µM at 37°C). SlyD is indicated as a known contamination of Ni-NTA-purified proteins.[22]

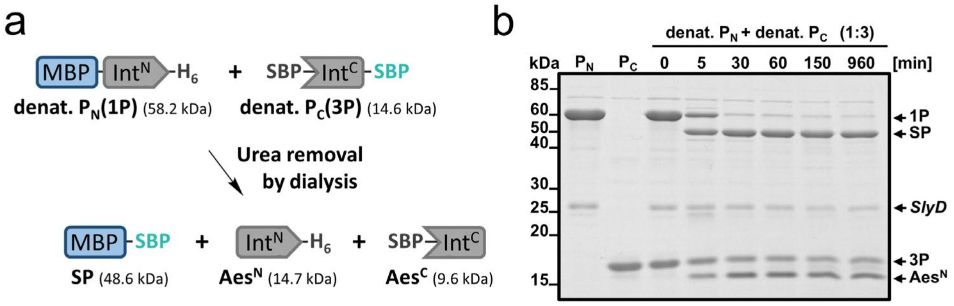


**Supplementary Figure 3** Collective refolding of both split intein precursor proteins leads to quantitative splicing. **a** Scheme of the experiment. Individual precursor proteins were under denaturing conditions with 8 M urea. After mixing, the urea was removed by dialysis and spontaneous protein *trans*-splicing was observed. **b** SDS-PAGE analysis of the reaction using denatured **1P** and denatured **3P** at 10 µM and 30 µM, respectively. The reaction was performed at 25°C. Aliquots were removed during the dialysis at the indicated time points, quenched with SDS sample buffer and analyzed by SDS-PAGE. Note that the N-terminal precursor **1P** is nearly completely consumed. A contaminating band was identified as SlyD as indicated.[22] Shown is a gel stained with Coomassie brilliant blue.

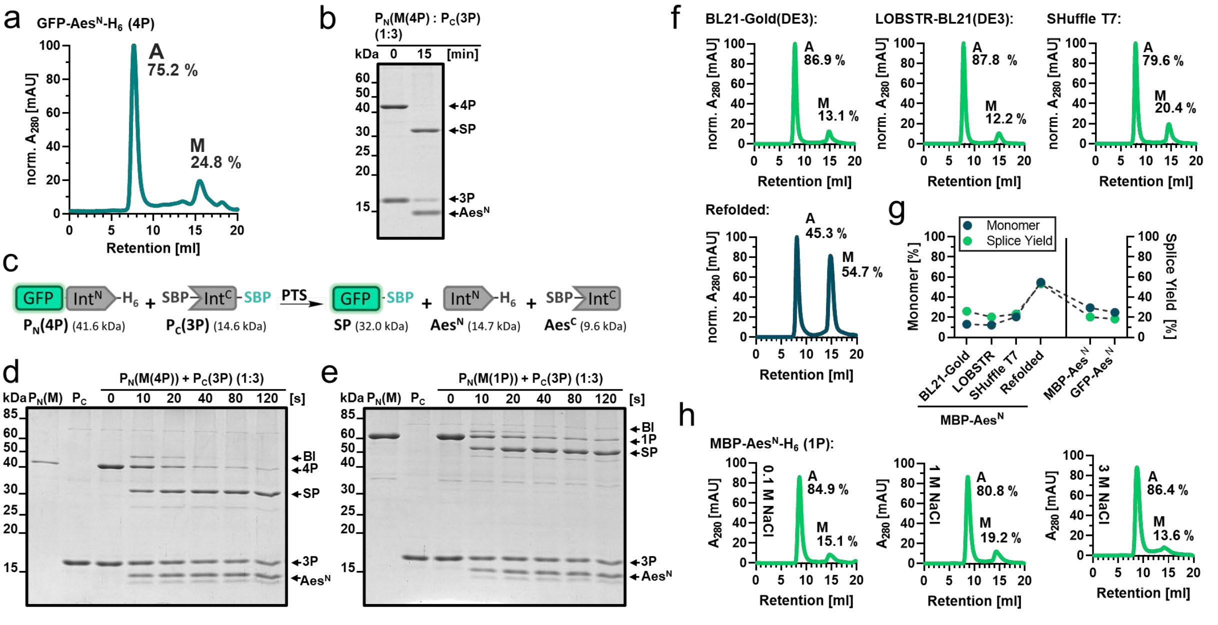


**Supplementary Figure 4** Aes$^{N}$ fragment aggregation is independent of the extein context, the expression system and ionic strength (additional data to Fig. 2). **a** Normalized UV elution profile of **4P** separated into two fractions on a Superdex 200 Increase 10/300 GL prepacked column (GE Healthcare) at a flow rate of 0.5 mL/min. **b** SDS-PAGE of the PTS reaction using the monomeric species of **4P** (10 µM) with three-fold excess of **3P** (30 µM) at 37°C. Shown is a gel stained with Coomassie brilliant blue. **c** Scheme of the PTS reaction using construct **4P** and **3P** (SBP-Aes$^{C}$-SBP) as performed in panel D and quantified in Figure 2D. **d** SDS-PAGE analysis of the time-resolved PTS reaction using the monomeric species of **4P** (10 µM) (purified with the monomer isolation protocol) with three-fold excess of **3P** (30 µM) at 37°C. Shown is a gel stained with Coomassie brilliant blue. **e** SDS-PAGE analysis of the time-resolved PTS reaction using the monomeric species of **1P** (10 µM) (purified with the monomer isolation protocol) with three-fold excess of **3P** (30 µM) at 37°C. **f** SEC-chromatograms of different **1P** samples (all at 1.5 mg/mL) purified by Ni-NTA chromatography, but produced by using different *E. coli* expression systems or an additional step-wise refolding step as indicated. SEC was performed on a Superdex 200 Increase 10/300 GL prepacked column (GE Healthcare) at a flow rate of 0.75 mL/min. **g** Correlation between the proportion of monomer and splice product formation. Shown are the monomer proportion after recombinant Aes$^{N}$ precursor production in different expression systems and with different N-exteins compared to the final splice yield reached with these precursors. The monomer proportion (blue) of **1P** and **4P** was analyzed by SEC using a Superdex 200 Increase 10/300 GL prepacked column (GE Healthcare) after incubation at 10 µM and 15°C for 24h. The splice efficiency (green) was analyzed by SDS-PAGE using **3P** in three-fold molar excess (10 µM to 30 µM, 37°C) to the Aes$^{N}$ precursor after 1 h incubation. **h** SEC-chromatograms of different samples of **1P** (1 mg/mL) after incubation with the indicated salt concentrations for 12 h. SEC was performed on a Superdex 200 Increase 10/300 GL prepacked column (GE Healthcare) at a flow rate of 0.75 mL/min. For panel **g**, *n* = 2 technical replicates regarding the splice yield determination. Data are presented as mean ± s.d normalized to the molecular weight.

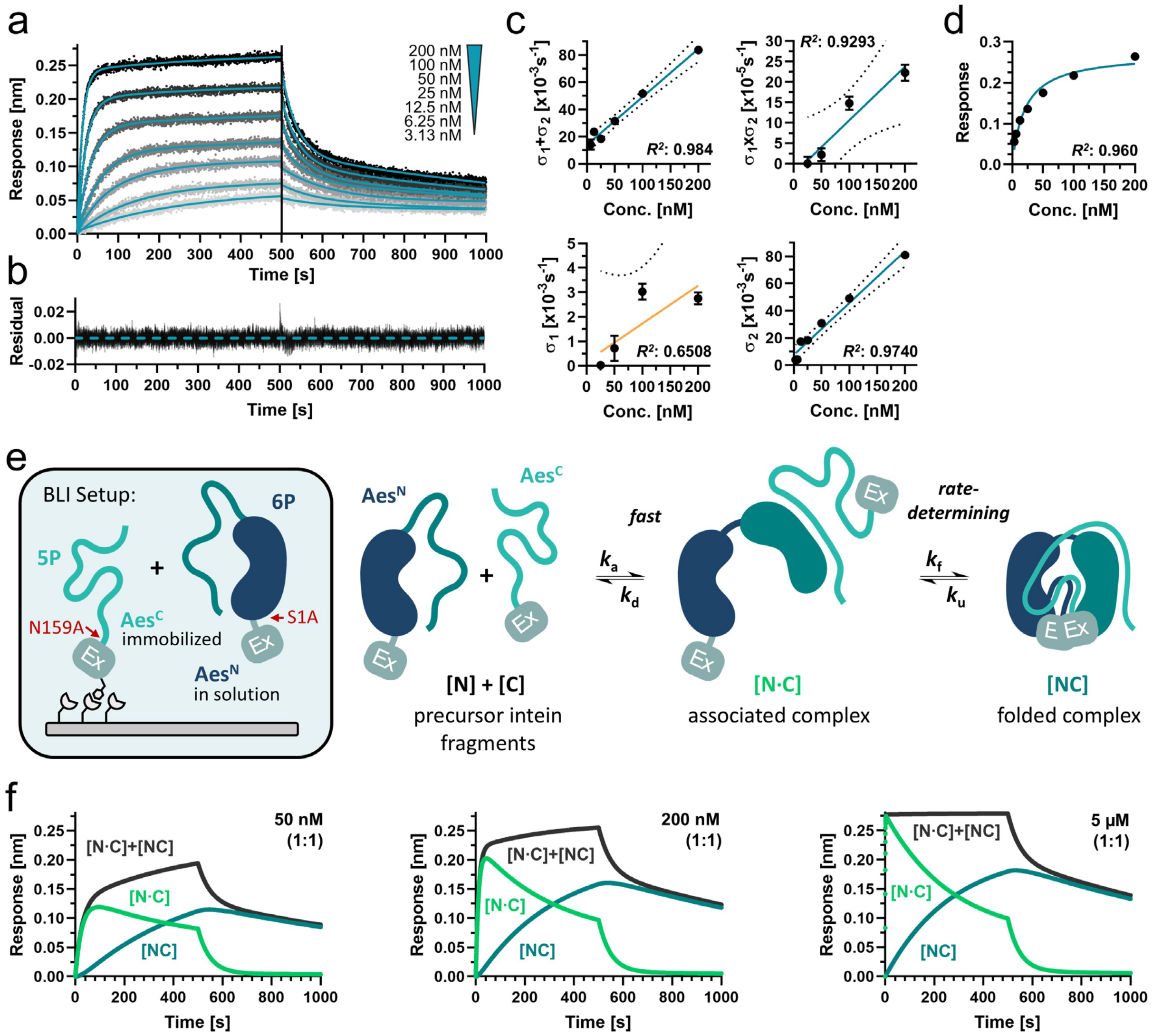


**Supplementary Figure 5** Investigation of the Aes split intein assembly mechanism by biolayer interferometry (BLI). **a** BLI sensorgram using biotinylated Aes$^C$-GFP(N159A) (**5P**) as immobilized ligand on the sensor tip and smt3-(S1A)Aes$^N$ (**6P**) as the analyte at different concentrations. The association and dissociation profiles were fitted by double exponential functions. **b** Residual plot of the BLI profiles to the applied fit. **c** Steady state analysis of the BLI response at 490 s – 500 s plotted against the analyte concentration. The upper graph shows the plot with a linear scale of the analyte concentration while for the lower graph a logarithmic scale was used. **d** Sum of $\sigma_1+\sigma_2$ from the double exponential function plotted against the concentration. The continuous line represents a linear fit. Product of $\sigma_1\sigma_2$ plotted against the concentration, $\sigma_1$ plotted against the concentration and $\sigma_2$ plotted against the concentration (details in the method section). **e** Scheme of the experimental BLI setup and the underlying intein assembly mechanism. **f** Simulation of the underlying binding kinetics of [N·C] and [NC] using the determined binding parameters at different concentrations. Note, that the sum [N·C] + [NC] resembles the BLI output. For panel **a** and **d**, *n* = 7 technical replicates were used measured at different concentrations as indicated. Data are presented normalized to a control without analyte protein **6P**. For panel **c**, data are derived from the fit shown in panel a and presented as mean ± s.d. derived from the previous fit.

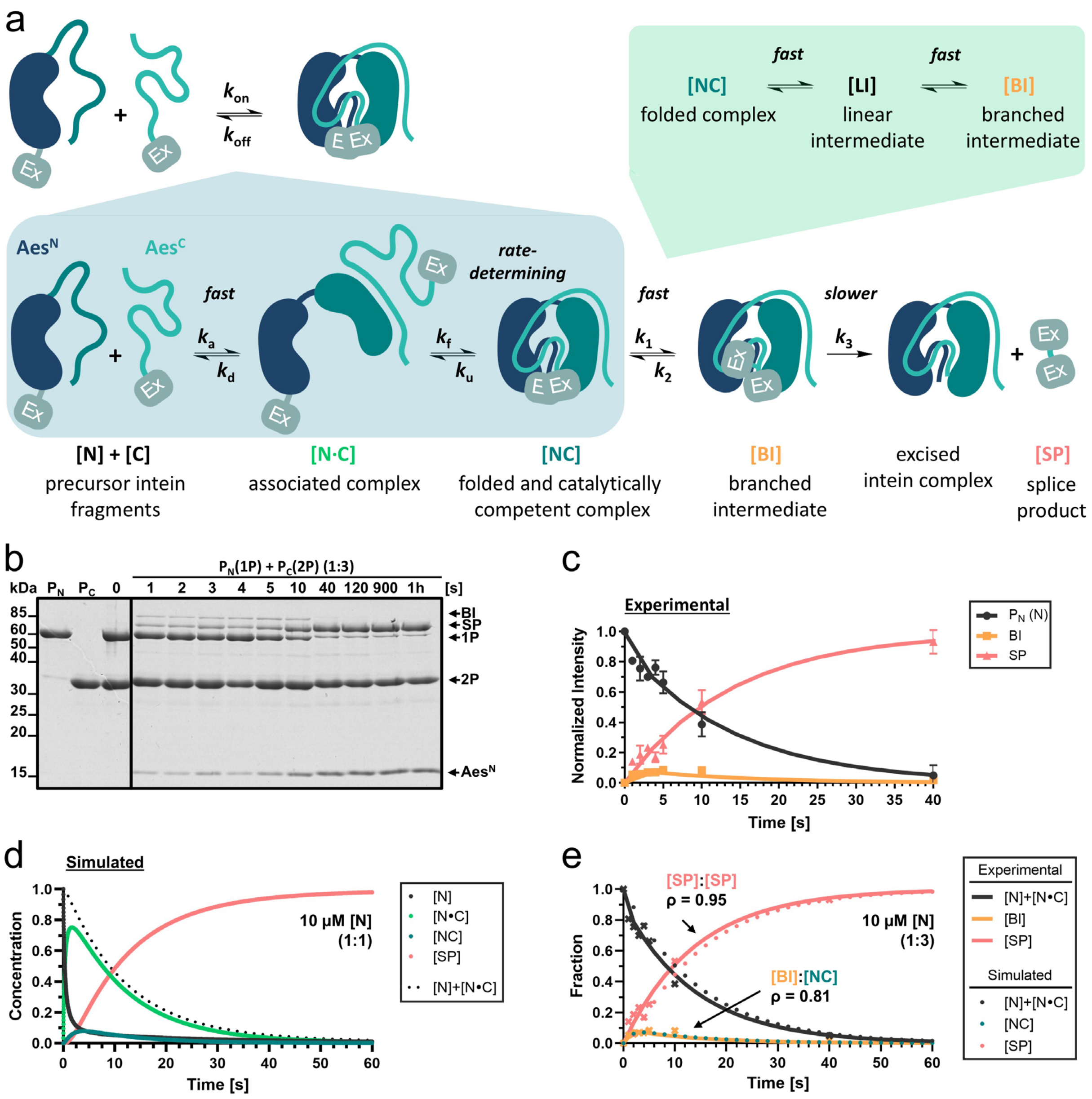


**Supplementary Figure 6** Investigation of the underlying relationship between the Aes assembly and splice mechanism. **a** Scheme of the underlying assembly and simplified thee-state kinetic splicing mechanism. **b** SDS-PAGE analysis of the PTS reaction using monomeric **1P** and **2P** with a three-fold molar excess of **2P** (10 µM **1P** to 30 µM **2P**) at 37°C. **c** Time course of the PTS reaction under pseudo-unimolecular conditions fitted to a three-step kinetic model showing the formation and resolution of the branched intermediate. Only the active fraction (80%) of the precursor protein $P_N$ was considered to determine the formation of the branched intermediate and the splice product. **d** Simulation of the underlying binding and splice kinetics over a time course of 60 s with $Aes^N$ and $Aes^C$ applied in equimolar concentrations (10 µM). **e** Correlation between the simulated binding kinetics and splice parameters and the experimentally determined BI and SP formation using $Aes^C$ in three-fold excess (10 µM $Aes^N$ to 30 µM $Aes^C$). Note, that the measured accumulation of the brached intermediate (BI) resembles the simulated time course of the folded complex [NC]. For panel **c**, *n* = 3 technical replicates. Data are presented as mean band intensity ± s.d. normalized to the molecular weight. For panel **e**, ρ-values were derived from a two-tailed non-parametric Spearman correlation test.

**Supplementary Table 6** Kinetic binding parameters of the Aes123 PolB1 intein.*

| **Binding Parameter** | | **Determined with inactive intein** | **Simulated for active intein** |
|---|---|---|---|
| association rate | $k_a$ | 3.53 ± 0.23 x $10^5$ $M^{-1}s^{-1}$ | |
| dissociation rate | $k_d$ | 1.76 ± 0.13 x $10^{-2}$ $s^{-1}$ | |
| folding rate | $k_f$ | 2.84 ± 1.02 x $10^{-3}$ $s^{-1}$ | 91.13 ± 6.17 x $10^{-3}$ $s^{-1}$ |
| unfolding rate | $k_u$ | 8.73 ± 0.90 x $10^{-4}$ $s^{-1}$ | |
| overall ass. rate | $k_{on}$ | 4.91 ± 1.83 x $10^4$ $M^{-1}s^{-1}$ | 29.59 ± 3.27 x $10^4$ $M^{-1}s^{-1}$ |
| overall diss. rate | $k_{off}$ | 7.51 ± 1.11 x $10^{-4}$ $s^{-1}$ | 7.51 ± 0.32 x $10^{-4}$ $s^{-1}$ |
| dissociation constant | $K_d$ | 11.7 ± 3.0 $M^{-9}$ | 0.48 ± 0.12 $M^{-9}$ |

| **Splice Kinetics** | | **Determined** | **Used in Simulation** |
|---|---|---|---|
| overall splice rate | $k_{total}$ | 91.13 ± 6.17 x $10^{-3}$ $s^{-1}$ | 91.13 ± 6.17 x $10^{-3}$ $s^{-1}$ |
| succinimide resolution | $k_3$ | 0.78 ± 0.13 x $10^{-3}$ $s^{-1}$ | |
| splice rate | $k_{splice}$ | | 0.78 ± 0.13 x $10^{-3}$ $s^{-1}$ |

* Experimentally determined binding parameters by biolayer interferometry using splice inactive constructs and simulated values for the active Aes intein. To consider the influence of active protein *trans*-splicing on the binding parameters, the splice kinetics determined with the monomeric construct **1P** and **3P** (10 µM to 30 µM, $k_{total}$ = 91.13 ± 6.17 x $10^{-3}$ $s^{-1}$) and monomeric **1P** and **2P** (5 µM to 15 µM, 70.04 ± 5.45 x $10^{-3}$ $s^{-1}$) were used.

**Supplementary Note 1** Biophysical and kinetical investigation of the underlying assembly mechanism by biolayer interferometry.

We investigated the underlying binding kinetics of the Aes123 PolB1 intein by biolayer interferometry using biotinylated $Aes^{C}$(N39A)-GFP (**5P**) as immobilized ligand applied to different concentrations of Smt3-(S1A)$Aes^{N}$ (**6P**) as analyte (Supplementary Fig. 5a,b,e). The BLI profiles revealed a biphasic binding behavior which suggests a similar multistep assembly mechanism as described for other split inteins.[23] To determine the binding parameters, we fitted the association and dissociation profiles by double exponential functions and extracted the rate constants by an analytical solution of the biphasic rate equations. Here, the dependencies of the exponents and their sums and products on the analyte concentration (C) were used to confirm the correct model and to determine the rate equations as described by Tiwari *et al.*.[12] The linearity between $\sigma_1 + \sigma_2$ and C verifies that the underlying binding mechanism reflects a linear biphasic reaction. Moreover, the linear relationship of the product $\sigma_1\sigma_2$ to C indicates a binding mechanism as depicted by the two-step conformational change model and confirms the proposed intein assembly mechanism (Supplementary Fig. 5c). The concentration dependent association rate ($k_a$) was obtained from the slope of the sum of the exponents plotted against the concentration with $k_a = 3.53 \pm 0.23 \times 10^5\ M^{-1}s^{-1}$. Based on the two-step conformational change model we calculated a $K_D$ value of 11.7 ± 3.0 nM with an overall $k_{on}$ of $4.91 \pm 1.83 \times 10^4\ M^{-1}s^{-1}$ and a $k_{off}$ of $7.51 \pm 1.11 \times 10^{-4}\ s^{-1}$ (Supplementary Table 6). This high affinity in the low nanomolar range is a typical feature of native split inteins and confirmed by the steady state analysis which assumes a similar $K_D$ value (14 nM) (Supplementary Fig. 5d). Consequently, we simulated binding curves at different concentrations together with the obtained rate constants of the dissociation rate ($k_d = 1.76 \pm 0.13 \times 10^{-2}\ s^{-1}$), the folding rate ($k_f = 2.84 \pm 1.02 \times 10^{-3}\ s^{-1}$) and the unfolding rate ($k_u = 8.73 \pm 0.90 \times 10^{-4}\ s^{-1}$). The simulations resemble the experimental BLI data and clearly identify the folding step as rate determining (Supplementary Fig. 5f).

However, it has to be considered that the affinity measurements were done with the inactivated constructs which probably lead to an underestimation of the real binding kinetics since the equilibrium between folding and unfolding is influenced by the splice rate ($k_{splice}$) (Supplementary Fig. 6a). In a multistep mechanism the rate constants of the rate limiting step defines the overall rate constants. However, the determined folding rate ($k_f$) is slower than the initially obtained overall splice rate ($k_{total}$). We simulated the real binding kinetics at equimolar concentrations (10 µM) by equating the folding rate constant with the previously determined overall splice rate of the PTS reaction ($k_{total} = k_f$). Moreover, a highly time-resolved splice assay was used to determine the single catalytic steps simplified by a three-state kinetic model (Supplementary Fig. 6b,c). We were not able to exactly define $k_1$ and $k_2$ due to the fast reaction rate and low accumulation of the branched intermediate (BI), but we could extract the BI resolution as rate determining step within the splice mechanism with $k_3 = 0.78 \pm 0.13 \times 10^{-3}\ s^{-1}$.

Interestingly, equating $k_3$ as $k_{splice}$ simulates binding and splice kinetics that assume overall splice rates which lie in the realm of the experimentally determined values (Supplementary Fig. 6d). The extrapolated kinetic parameters for the active intein fragments (Supplementary Table 6) suggest a $K_d$ of 0.48 ± 0.12 nM and the corresponding simulated splice product formation even slightly underestimates the experimentally measured product formation (Supplementary Fig. 6e). However, both curves still show a good Spearman correlation coefficient with $\rho = 0.95$ ($P = 0.00035$). Moreover, the simulated appearance of the folded complex strongly adopts the time-resolved accumulation of the branched intermediate with a correlation of $\rho = 0.81$ ($P = 0.011$). This correlation corroborates the fast transesterification into the BI upon fragment folding. This fast conversion ($k_1$) in combination with a fast resolution of the branched oxo-ester intermediate ($k_2$) explains the low BI accumulation and the difficulty in determining these parameters.

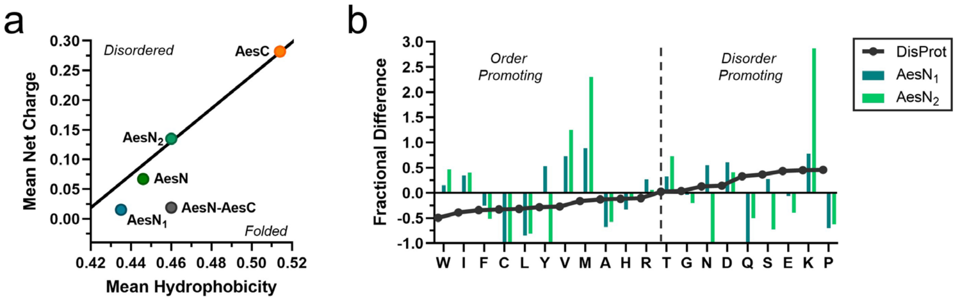


**Supplementary Figure 7** Sequence-based prediction of the folding state by charge-hydrophobicity plot and amino acid composition profiling. **a** Charge-hydrophobicity (CH) plot of the $Aes^{N}$ fragment, the $Aes^{C}$ fragment, the N-terminal lobe of $Aes^{N}$ ($AesN_1$, aa1-68), the C-terminal lobe of $Aes^{N}$ ($AesN_2$, aa69-120) and the fused complex structure ($Aes^{N}$-$Aes^{C}$). A linear boundary (R = 2.785 H −1.151) separates the CH plot into disordered (above) and ordered (below) regions. **b** Comparison of amino acid composition of $AesN_1$, $AesN_2$ and the Disprot 9.6 database (representing known disordered regions) with the general protein database UniProtKB/TrEMBL 2024_04. The bar graph represents the amino acid enrichment (positive) or depletion (negative) relative to TrEMBL. The $AesN_1$ and $AesN_2$ amino acid composition in **b** does not show the typical distribution as known for intrinsically disordered proteins.[24] Rather the amino acid distribution reflects order promoting as well as disorder promoting characteristics. These sequence characteristics accompany the known/putative intein assembly mechanism which requires the same amino acid sequence in disordered and ordered states at different assembly steps. Notably, the overrepresentation of the charged lysine residue in $AesN_2$ indicates a central role in holding the intein fragment structure partially disordered. Note that a high net charge in the $IntN_2$ lobe is conserved among inteins.[23]

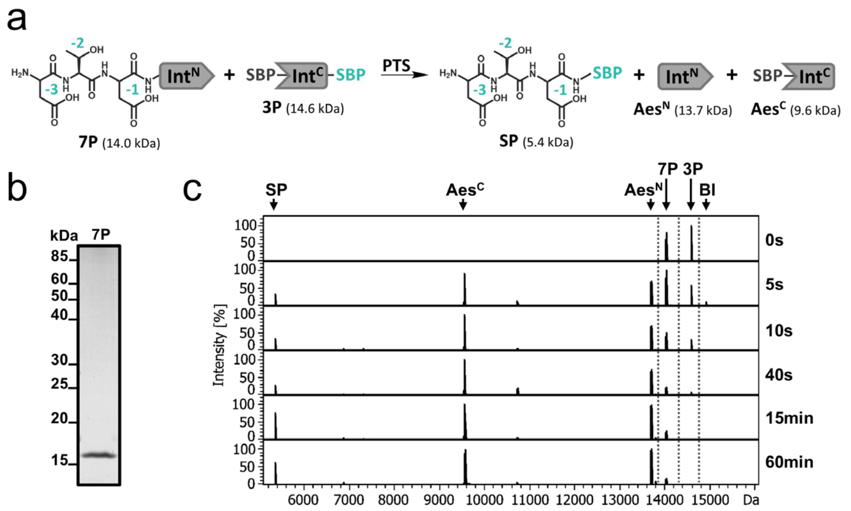


**Supplementary Figure 8** Splice activity of the Aes$^{N}$ precursor with a short N-terminal extein of only 3 residues. **a** Scheme of the PTS reaction. **b** SDS-PAGE analysis of purified monomeric DTD-Aes$^{N}$ (**7P**) (prepared by the monomer isolation protocol). **c** ESI-MS analysis of the PTS reaction using the monomeric species of **7P** in low excess to **3P** (15 µM to 10 µM) at 37°C.

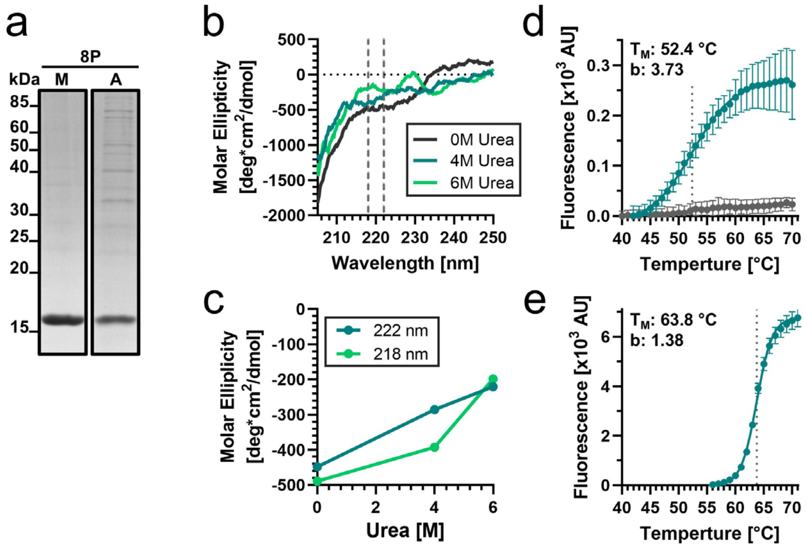


**Supplementary Figure 9** Biophysical investigation of the $\text{Aes}^{\text{N}}$ precursor (additional data to Fig. 3a). **a** SDS-PAGE analysis of the purified monomeric (prepared by the monomer isolation protocol) and aggregated species of DTD-$\text{Aes}^{\text{N}}$(S1A) (**8P**) used for the biophysical analysis. **b** Far UV circular dichroism spectroscopy of 10 µM **8P** containing different concentrations of urea as indicated. **c** Molar ellipticity of **8P** at 218 nm and 222 nm as shown in **b** plotted against urea concentration. **d** Thermal shift assay of DTD-$\text{Aes}^{\text{N}}$(S1A) (**8P**; 0.2 g/L) shown next to a buffer control (gray). The data was fitted to a sigmoidal four-parameter logistic equation to determine the melting temperature ($T_M$) and the folding cooperativity (b; Hill coefficient (no unit)). To this end, the data profile was cut at the highest value (at 70°C) to account for post-peak aggregation of protein-dye complexes leading to quenching of the fluorescence signal. **e** Thermal shift assay of MYIDTD-$\text{Aes}^{\text{N}}$(S1A)-GSH-$\text{Aes}^{\text{C}}$(N159A)-SVYLN (**9P**, 0.5 g/L) fitted to a sigmoidal four-parameter logistic equation. For panel **b**, $n$ = 3 technical replicates. Data are presented as mean molar ellipticity corrected for concentration. For panel **d** and **e**, $n$ = 6 -7 technical replicates. Data are presented as mean ± s.d. normalized to the highest fluorescence intensity.

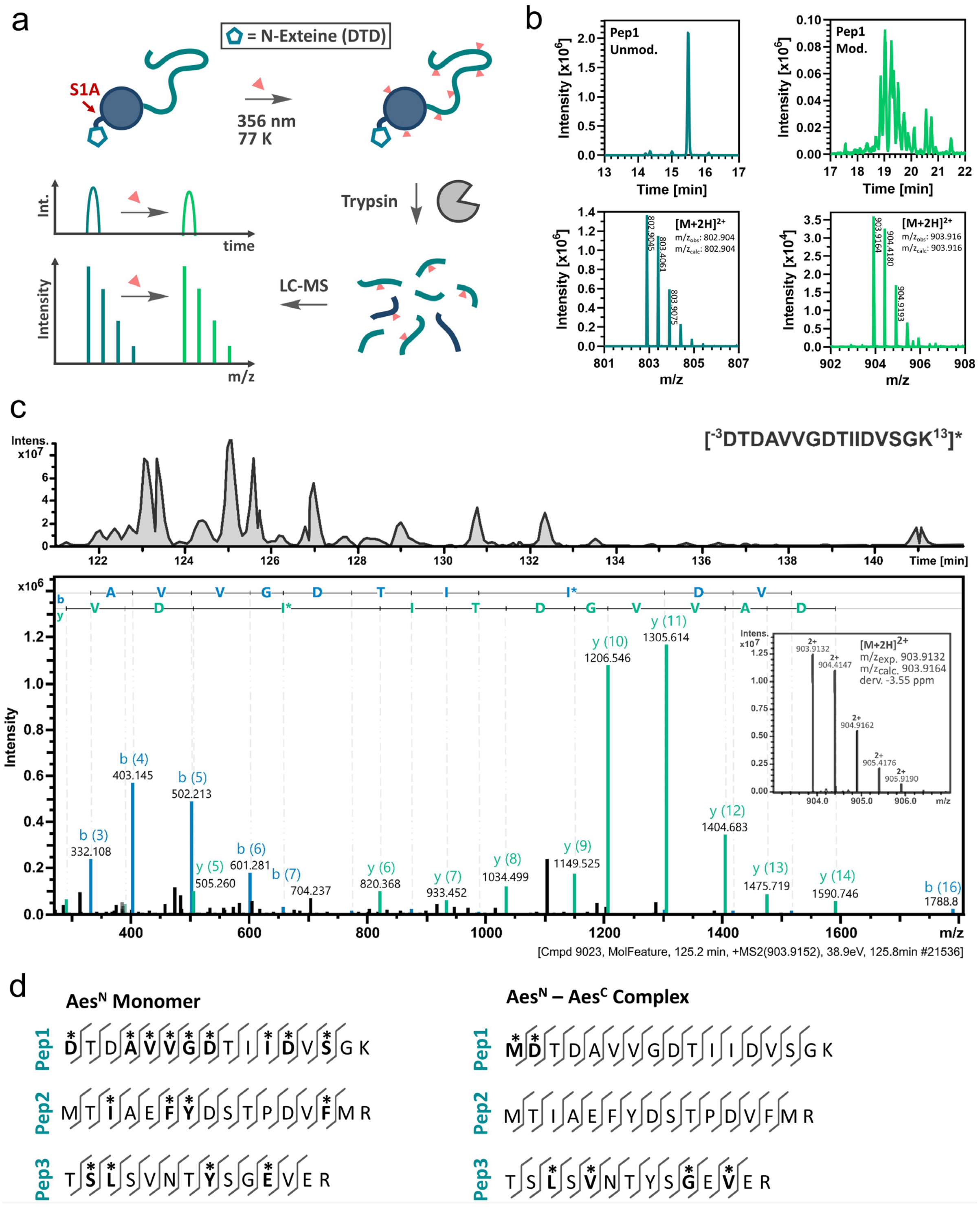


**Supplementary Figure 10** Principle of the photoreactive carbene labeling and LC-MS based footprinting (additional data to Fig. 3c). **a** Scheme of the experimental workflow. **b** Representative extracted ion chromatogram (EIC) after protein trypsinization of unlabeled **Pep1** (blue) and carbene-labeled **Pep1** (green) of the carbene footprinting approach using 10 µM DTD-Aes$^{N}$(S1A) (**8P**). The upper panels show the EIC of unlabeled **Pep1** (blue) and carbene-labeled **Pep1** (green). The lower panels show the MS spectra of unlabeled **Pep1** (blue) and carbene-labeled **Pep1** (green). **c** LC-MS/MS analysis of the carbene-modified and trypsinnated peptides. Shown are representative samples of the non-quantitative tandem MS analysis of carbene-labeled **Pep1.** The upper panel shows the EIC of carbene-labeled **Pep1**, containing various singly labeled species with different retention times. The lower panel shows the MS$^{2}$ spectrum of one such **Pep1** species, which revealed that the carbene-label was inserted at position I8. The inset shows the MS$^{1}$ spectrum of carbene-modified **Pep1**. **d** Summary of the identified carbene modification sites by the non-quantitative tandem MS analysis of **Pep1**, **Pep2** and **Pep3** compared between the monomeric Aes$^{N}$ precursor DTD-Aes$^{N}$(S1A) (**8P**) and the Aes$^{N}$-Aes$^{C}$ complex structure of MDTD-Aes$^{N}$(S1A)-GSH-Aes$^{C}$(N159A)-SVYLN (**10P**). Indicated are amino acids identified as carbene labeled (asterisk).

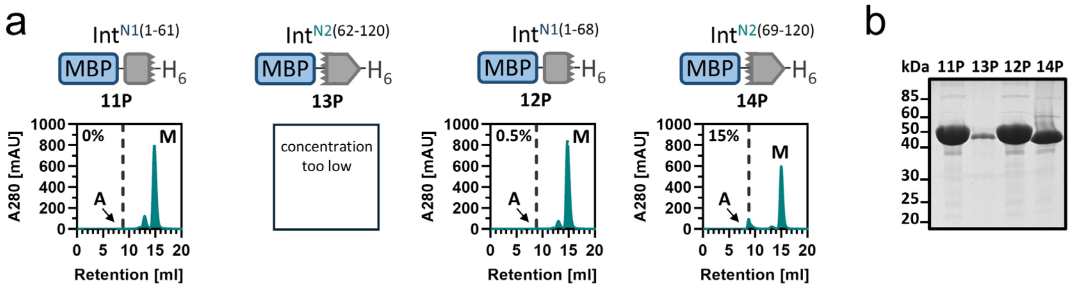


**Supplementary Figure 11** SEC analysis of the single Aes$^N$ fragment segments fused to MBP (additional data to Fig. 3d). **a** SEC-analysis with 150 µM of each indicated protein. The construct **13P** was not tested at these high concentrations due to lower solubility. **b** SDS-PAGE analysis of purified proteins after recombinant protein expression and Ni-NTA affinity chromatography. Note that the proteins were produced under the same expression and purification conditions, leading to different protein yields.

**Supplementary Table 7** Crystallographic data collection and structure refinement.*

| **Data Collection** | PDB ID: 9HTH |
|---|---|
| Space Group | P $2_1$ $2_1$ $2_1$ |
| Unit-cell parameters | |
| a, b, c (Å) | 30.1, 63.0, 162.4 |
| α, β, γ (°) | 90, 90, 90 |
| Resolution range (Å) | 34.13 - 1.38 (1.43 - 1.38) |
| Observed reflections | 827722 (83755) |
| Unique reflections | 64820 (5518) |
| Completeness (%) | 96.27 (85.92) |
| Multiplicity | 12.8 (13.1) |
| I/σ(I) | 16.43 (1.73) |
| $CC_{1/2}$ | 0.999 (0.782) |
| Willson B-factor | 19.07 |
| $R_{meas}$ | 0.077 (1.235) |
| **Refinement** | |
| $R_{work}$, $R_{free}$ | 0.195 / 0.228 |
| Reflections | 62612 |
| Ramachandran favored (%) | 98.53 |
| Ramachandran allowed (%) | 1.47 |
| Ramachandran outliers (%) | 0 |
| RMS bonds (Å) | 0.007 |
| RMS angles (°) | 1.27 |
| Average B-factor | 27.16 |
| macromolecules | 26.7 |
| solvent | 33.7 |
| Number of TLS groups | 15 |

* Values in parentheses refer to the highest-resolution shell.

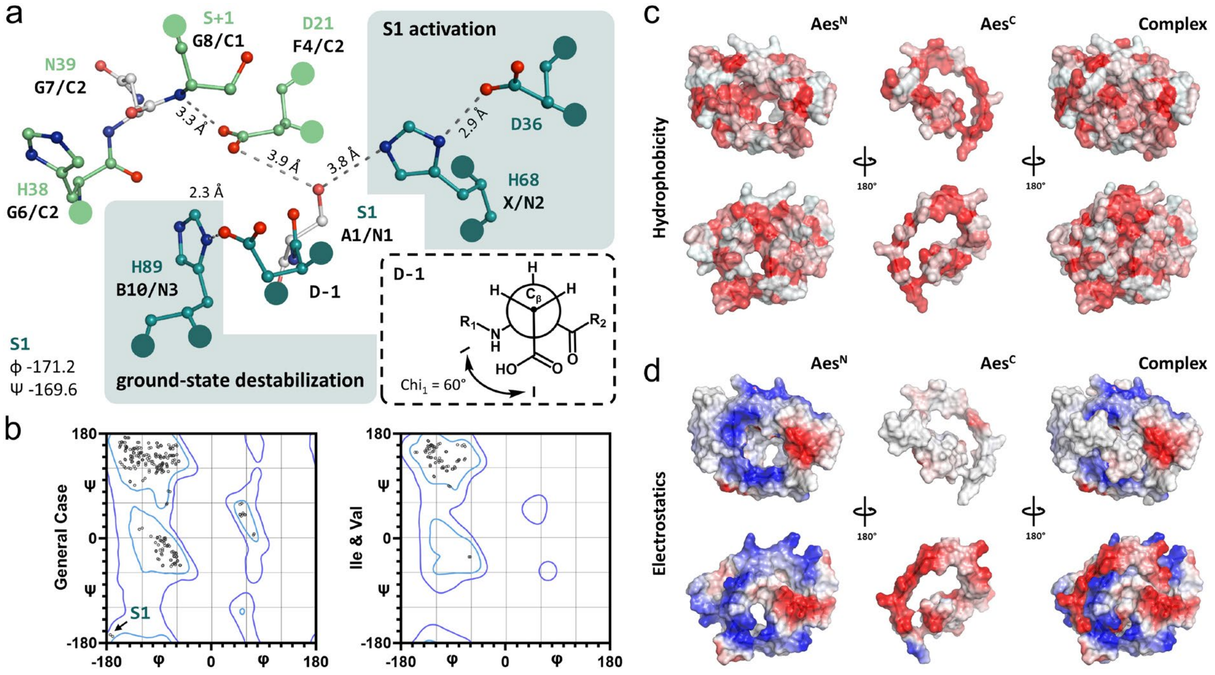


**Supplementary Figure 12** Structural analysis of the catalytic center and the charge/hydrophobicity distribution among the intein fragments using the crystal structure of the Aes123 PolB1 intein. **a** Essential catalytic residues within the active site of the Aes123 PolB1 intein with the residues of the Aes$^{N}$ fragment marked in blue and the Aes$^{C}$ fragment labeled in green. Indicated are the catalytic residues and the corresponding conserved motifs as well as the distance between selected functional groups specified in (Å). The side chains of Ser1 and Asn159 (gray) are remodeled in the shown structure using Pymol as the intein was crystalized with S1A and N159A mutations. The Newman projection illustrates the energetically unfavorable gauche(-) conformation of the Asp-1 side chain. The conformation is defined by an angel of 60° between the Cγ and the amide nitrogen ($R_1$: N-terminal towards Thr-2; $R_2$: C-terminal with the scissile amide bond towards Ser1). **b** Ramachandran plots of the Aes123 PolB1 complex structure generated using MolProbity[25] highlighting the conformational constraints on the S1 backbone. The plot on the left-hand side treats all residues except Val, Ile, Pro and Gly. The plot on the right-hand side treats Val and Ile residues. **c** Surface view of the Aes123 PolB1 intein complex structure and the individual Aes$^{N}$ and Aes$^{C}$ fragments showing the surface hydrophobicity in red scale. **d** Surface view showing the electrostatic properties in red scale (negatively charged) and blue scale (positively charged).

**Supplementary Note 2** Crystal structure of the Aes123 PolB1 intein.

We obtained crystals of the inactivated and fused Aes123 PolB1 precursor construct with 5 extein residues on each flank (construct **9P**) that diffracted to 1.38 Å resolution (Fig. 4c). The asymmetric crystal unit contained two molecules that exhibited only slight differences in electron density. Most notably, while in chain A none of the N-terminal and only three of the C-terminal extein residues were resolved, in chain B four and five residues, respectively, were visible. Overall, the intein adopted the typical horseshoe fold. The artificial GSH-linker between the $Aes^N$ and $Aes^C$ parts was well resolved. Interestingly, the region around residues ~25 to 40 (β3 and β4) showed high structural similarity to the insertion first found at the corresponding region in thermophilic inteins,[26] and later also in the mesophilic cysteine-less PolB16 OarG intein[27] (Supplementary Fig. 13a). Characteristically, the insertion folds into two additional β-sheets that extend the β7-strand at residues Y64-T69 (Fig. 4c,d). The preceding α-helix (α1) is disrupted by P25 and shorter than typically observed in the thermophilic counterparts and the PolB16 OarG intein.

The Aes123 PolB1 intein also contains the recently discovered conserved histidine that represents a signature sequence of cysteine-independent inteins in the newly defined motif NX (Supplementary Fig. 13b).[27] This residue, His68, is positioned close to the side chain of Ser1 (when remodeled back into the crystallized intein with the S1A mutation), thereby further supporting its proposed role in activation of the Ser1 side chain (Supplementary Fig. 12a). Interestingly, together with Asp37 a potential seryl-histidyl-aspartate motif is formed, the most common catalytic triad known from enzymatic studies.[28] This catalytic motif is unique to the Aes123 PolB1 intein within the known intein structures and might explain the high reaction rate of this cysteine-independent intein.

Another notable observation from the crystal structure is an unusual conformation of the first N-terminal extein amino acid. This residue, Asp-1, adopts an allowed but energetically unfavorable gauche-(g-) conformation, defined by an angel of + 60° between Cγ and the amide nitrogen (Supplementary Fig. 12a). The g- conformation appears to be stabilized by a hydrogen-bonding contact between the side chain carboxyl group with the π-nitrogen of the catalytic motif B/N3 histidine His89 (distance of 2.7 Å in chain B). This conformation indicates a functional role of His89 in the distortion of the upstream scissile bond to help facilitate the initial N-O acyl shift. The crystal structure gives hints to slight conformational strains on the scissile peptide bond with torsion angels of φ = -171.2 and ψ = -169.6 in chain B (Supplementary Fig. 12b).

With respect to the interaction of the $Int^N$ with the $Int^C$ fragment, the crystal structure revealed the expected regions of electrostatic complementarity in the folded intein complex while hydrophobic residues are rather uniformly distributed in structure and sequence (Supplementary Fig. 12c,d). The electrostatic complementarity is mostly localized in the intermolecular β-sheet composed of β7 ($Aes^N$) and β13 ($Aes^C$) and was consciously retained during the engineering process. Further, the structural integrity of the folded intein complex and the residue's physicochemical properties were considered for the engineering-based amino acid substitutions using the structural X-ray data.

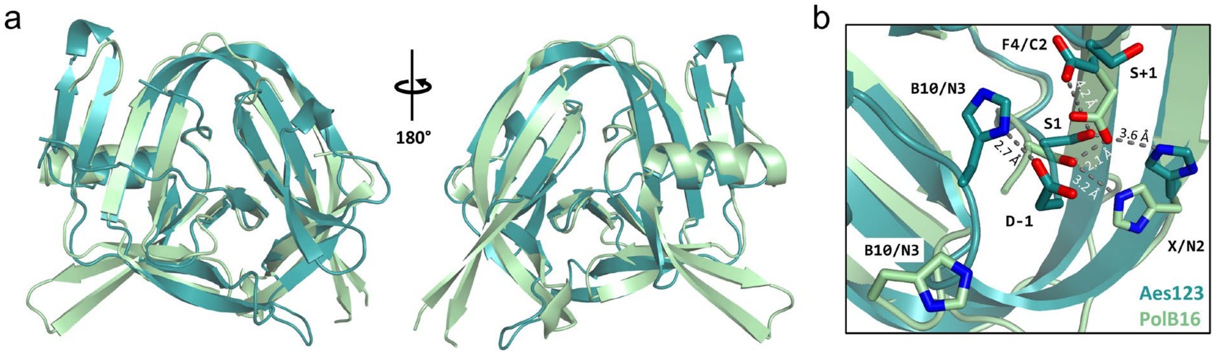


**Supplementary Figure 13** Structural comparison between the cysteine-less Aes123 PolB1 and the PolB16 inteins.[27] **a** Overall structural alignment between both inteins. **b** Structural alignment of the catalytic center of both inteins, highlighting the previously described unusual position of the catalytic B10/N3 histidine in the PolB16 intein (which has to be seen in context with the following amino acid residues that were found disordered in the PolB16 crystal structure),[27] and the previously discovered catalytic block X (N2) histidine, which is conserved among cysteine-independent inteins.[27] Note that Ser1 side chains were modeled into the structures using PyMol, as the double mutants S1A and N159A (N183A for the PolB16 intein) were used for crystallization.

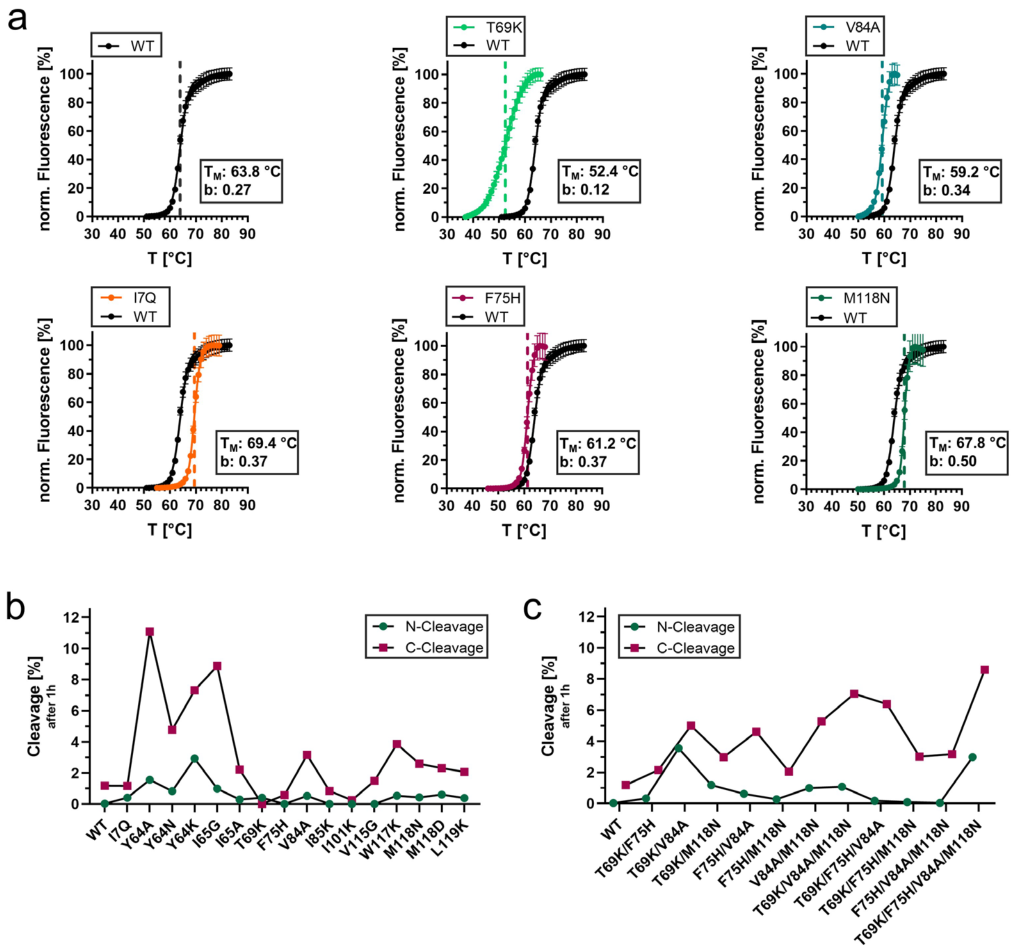


**Supplementary Figure 14** Influence of the aggregation-reducing mutations on the extent of side reactions and on the folding cooperativity to the canonical intein complex structure. **a** Thermal stability of the indicated, selected single mutants compared to the parent $Aes^N$(S1A)-$Aes^C$(N159A) (WT; protein **9P**) fused complex structure. The denaturation profiles were fitted to a sigmoidal four-parameter logistic equation to determine the indicated melting temperatures ($T_M$) and the Hill coefficient (b) of the fit to investigate the cooperativity in defolding. **b** N- and C-cleavage of the single mutants in the PTS reaction after 1 h ($P_N$ and $P_C$ precursors were used at 5 and 15 μM concentration, respectively). **c** N- and C-cleavage of the double, triple and quadruple mutants in the PTS reaction after 1h. For panel **a**, *n* = 5 - 8 technical replicates. Data are presented as mean ± s.d. normalized to the highest fluorescence intensity. For panel **b** and **c**, *n* = 2 technical replicates. Data are presented as mean ± s.d. normalized to the molecular weight.

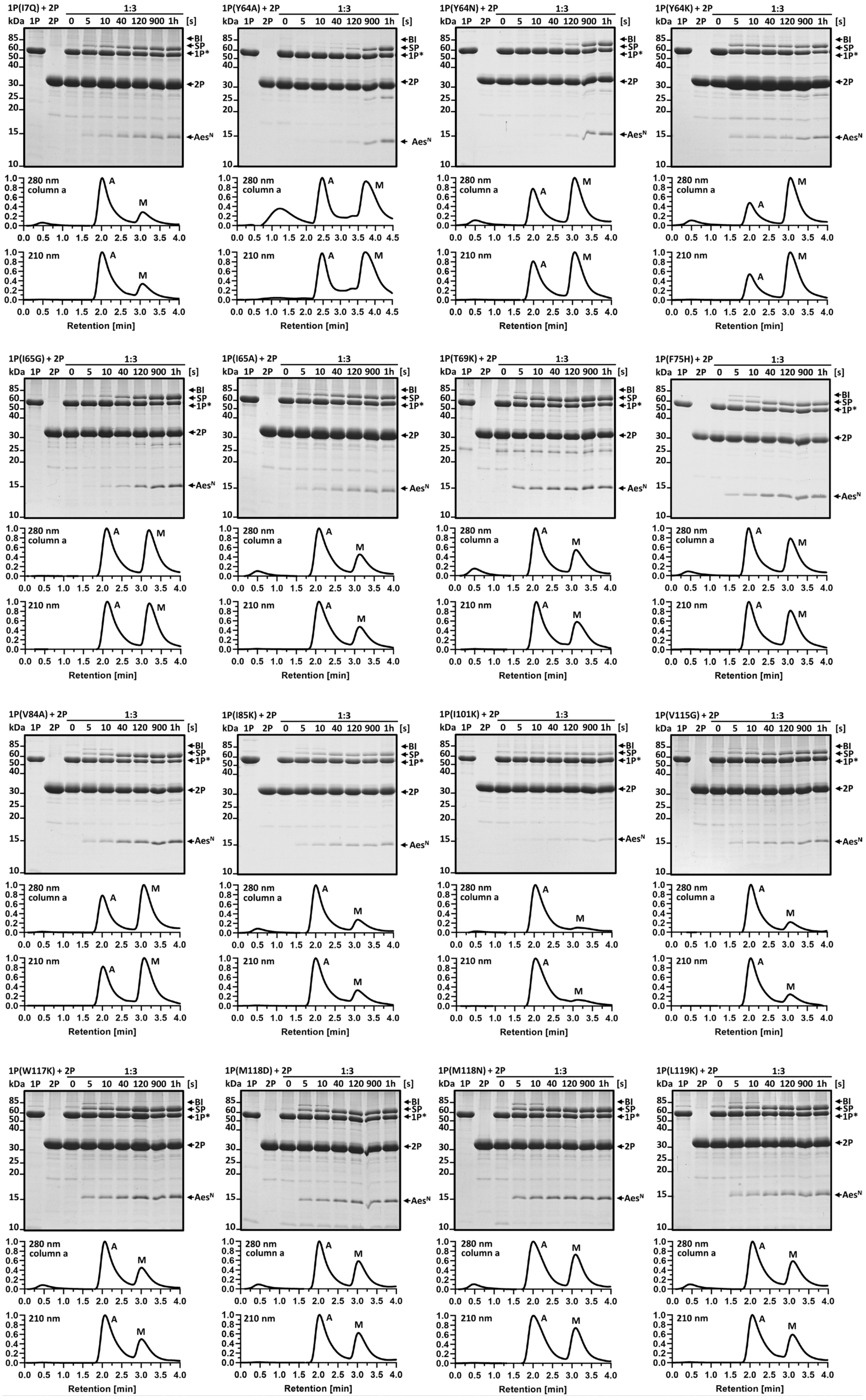


**Supplementary Figure 15** Continued on the next page.

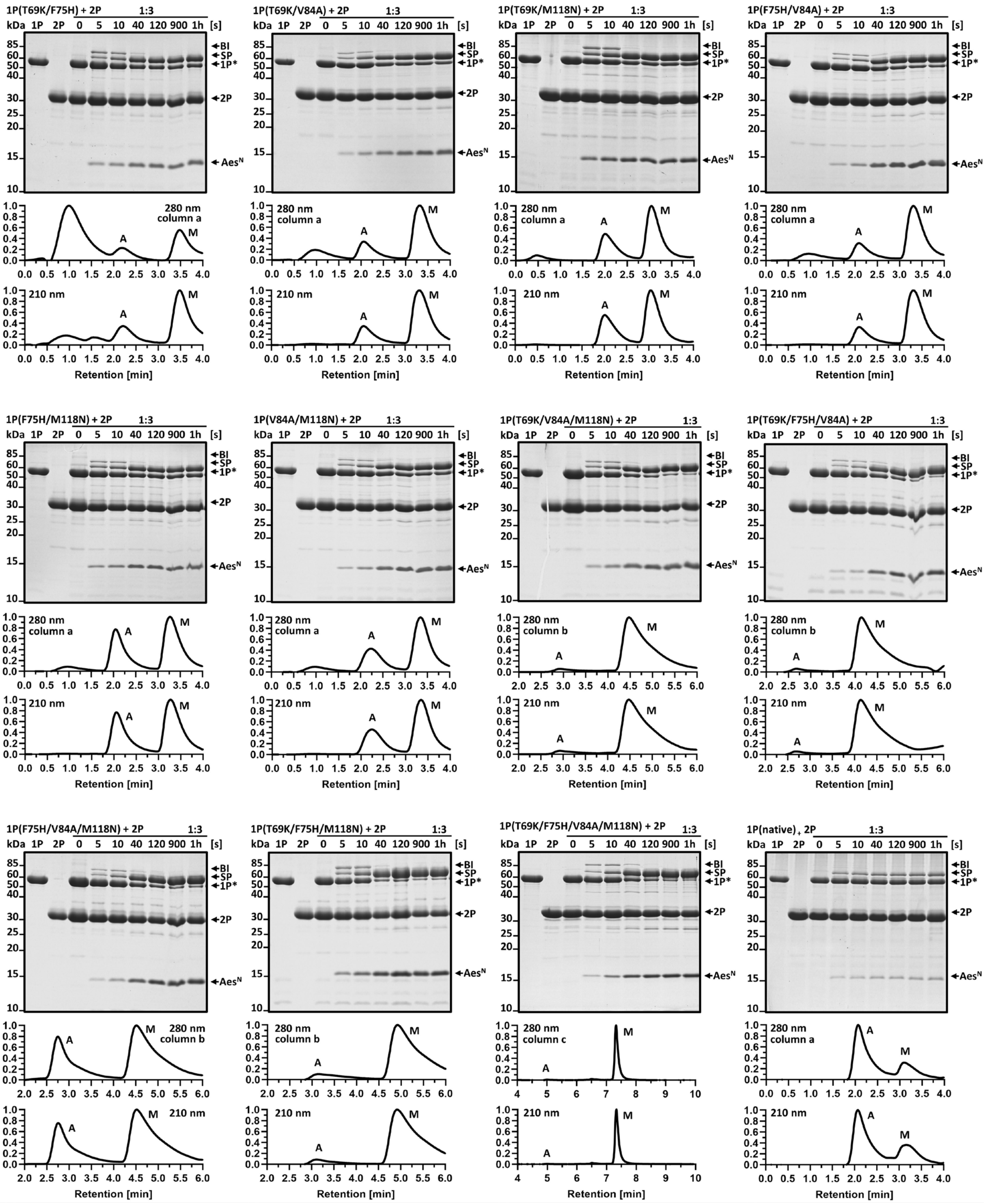


**Supplementary Figure 15** Analysis of the aggregation-reducing mutants of MBP-AesN-$H_6$ (additional data to Fig. 5a). For each mutant protein *trans*-splicing activity was analyzed using Coomassie-stained SDS-PAGE (upper panel) and the monomer/aggregate ratio was determined using analytical SEC and measuring protein absorbance at 280 nm (lower panel). Aggregate (A) and monomeric species (M) are indicated. Note that different SEC columns were used over the course of the project (**columns a**, **b** and **c**). In some cases, an extra signal of unknown origin (marked with asterisk) eluting well ahead of void volume retention time was observed in the 280 nm channel, which, importantly, did not correspond to a protein. For this reason, also the absorbance at 210 nm was monitored and is shown for each sample. PTS reactions were carried out at 37°C and using the indicated mutant of the N-terminal precursor ($P_N$) with a three-fold molar excess of the C-terminal precursor ($P_C$) Aes$^C$-GFP (**2P**) (10 µM and 30 µM, respectively). The three SEC columns used were: **column a**, analytical AdvanceBio SEC 120Å 1.9 µm, 2.1 × 150 mm PEEK pre-packed column (Agilent), operated at a flow rate of 0.1 mL/min; **column b**, analytical AdvanceBio SEC 200Å 1.9 µm, 2.1 × 150 mm PEEK pre-packed column (Agilent), operated at a flow rate of 0.1 mL/min; **column c**: analytical AdvanceBio SEC 200Å 1.9 µm, 4.6 × 300 mm pre-packed column (Agilent), operated at a flow rate of 0.35 mL/min.

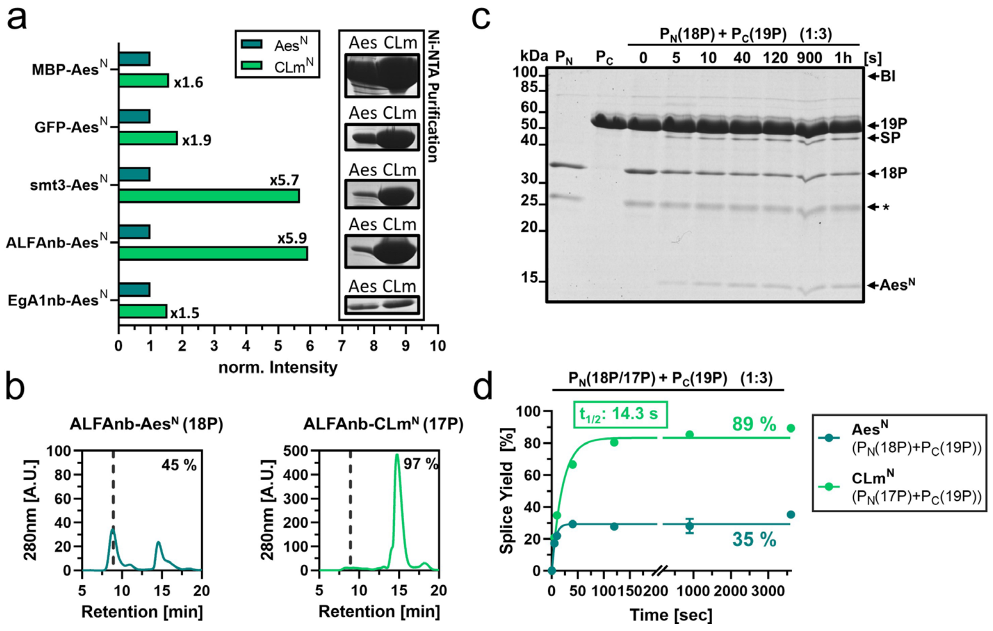


**Supplementary Figure 16** The Aes123 PolB1 triple mutant increases the expression yield and protein purity (additional data to Fig. 6a). **a** Densitometric analysis to compare the yields of various purified Aes$^{N}$ precursor proteins with the corresponding CLm mutants. The intensities were normalized to the Aes$^{N}$ precursor construct of each pair of proteins. **b** SEC-profiles of the construct ALFAnb-Aes$^{N}$-H$_6$ (**18P**) with the native Aes$^{N}$ sequence compared to the CLm mutant ALFAnb-CLm$^{N}$-H$_6$ (**17P**), recorded directly after Ni-NTA affinity purification. Note that the scale of the y-axis was adjusted for a better visualization. **c** SDS-PAGE analysis of the PTS reaction using **18P** together with **19P** used in three-fold molar excess. (5 µM and 15 µM at 37°C). **d** Time course of the PTS reactions (also shown in Fig. 6a,b) of either **18P** (blue) or **17P** (green), both used without monomer separation by SEC-purification, together with H$_6$-Smt3-Aes$^{C}$-GFP (**19P**) in three-fold molar excess (5 µM and 15 µM, respectively). The reactions were analyzed as pseudo-first order reactions and fitted to a one-phase exponential equation. See Fig. 6b for the SDS-PAGE analysis of the reaction performed with **17P** and **19P** under analogous conditions. For panel **d**, *n* = 2 technical replicates. Data are presented as mean ± s.d. normalized to the molecular weight.

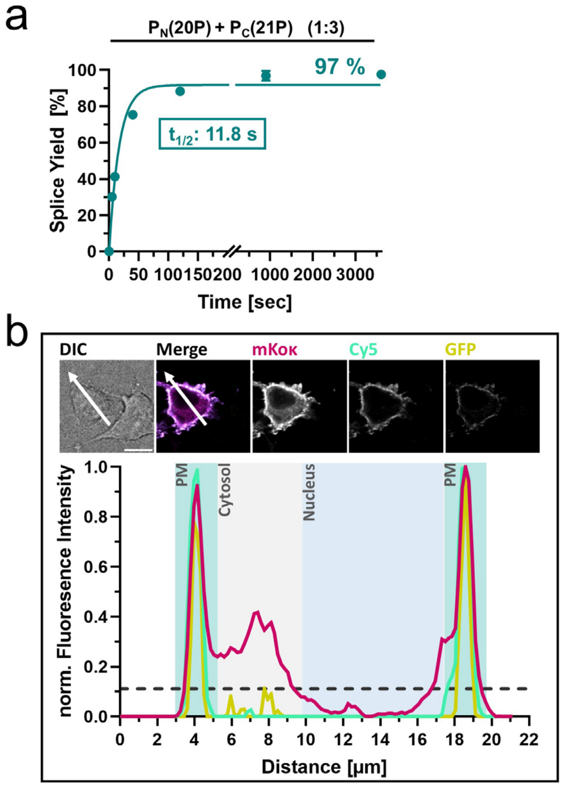


**Supplementary Figure 17** Intein-mediated generation of a bispecific nanobody-dimer for cell surface labeling (additional data to Fig. 6c-f). **a** Time course of the PTS reaction (Fig. 6c,d) at 37°C of EgA1nb-CLm$^{N}$-H$_6$ (**20P**) and H$_6$-CLm$^{C}$-ALFAnb-SBP (**21P**) (5 µM and 15µM, respectively). **b** Spatial analysis of cellular labeling as described in Fig. 6e,f. Shown are CLSM images of transiently transfected HeLa cells (upper panel) and the quantitative analysis of fluorescence intensity (lower panel) across the cell section along the white arrow. HeLa cells were expressing the extracellular domains of the EGF-receptor (EGFR) exposed on the cell surface together with an intracellular fluorescent marker (mKoκ) (Fig. 6e). Cells were incubated with 250 nM Cy5-labeled **SP(20P-21P)** (cyan) for 10 min at 37°C. Afterwards, 50 nM of GFP-ALFAtag (**22P**) (yellow) was added to the cells and incubated for additional 5 min. The scale bar represents 10 µm. The dashed line marks the maximal background fluorescence in the 488-channel (GFP). For panel **a**, *n* = 2 technical replicates. Data are presented as mean ± s.d. normalized to the molecular weight.

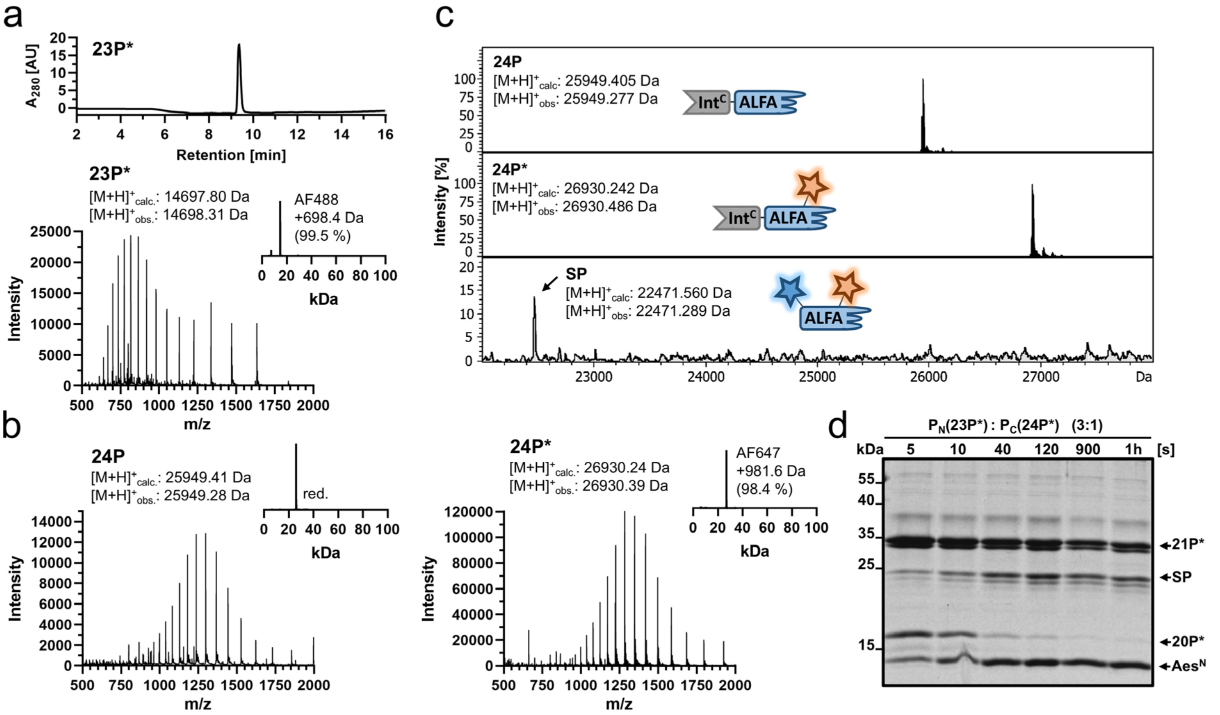


**Supplementary Figure 18** Nanobody functionalization using intein-mediated dual thiol-bioconjugation (additional data to Fig. 6g,h). **a** RP-HPLC and ESI-MS analysis of the used AF488-labeled CAD-Aes$^{N}$ (**23P***). **b** ESI-MS analysis of H$_6$-Aes$^{C}$-ALFAnb(G47C)-SBP (**24P**) before and after labeling by maleimide-AF647 (**24P***). **c** ESI-MS analysis of PTS reaction (shown in Fig. 6g) showing the stepwise dual labeling of the ALFA nanobody of construct H$_6$-Aes$^{C}$-ALFAnb(G47C)-SBP (**24P**) with the AF488-maleimide by thiol bioconjugation of G47C and subsequent labeling with AF647-labeled tripeptide CAD (N-extein of **23P***) by protein *trans*-splicing. **d** SDS-PAGE analysis via Coomassie staining of the PTS reaction between **23P*** and **24P*** at 37°C (proteins at 5 µM and 15 µM, respectively). Note that the observed labeling efficiency differs between ESI-MS (shown in **c**) and SDS-PAGE analysis (shown in **d**) due to partial thioether cleavage, which removes the maleimide label during sample preparation for SDS-PAGE.[29, 30]

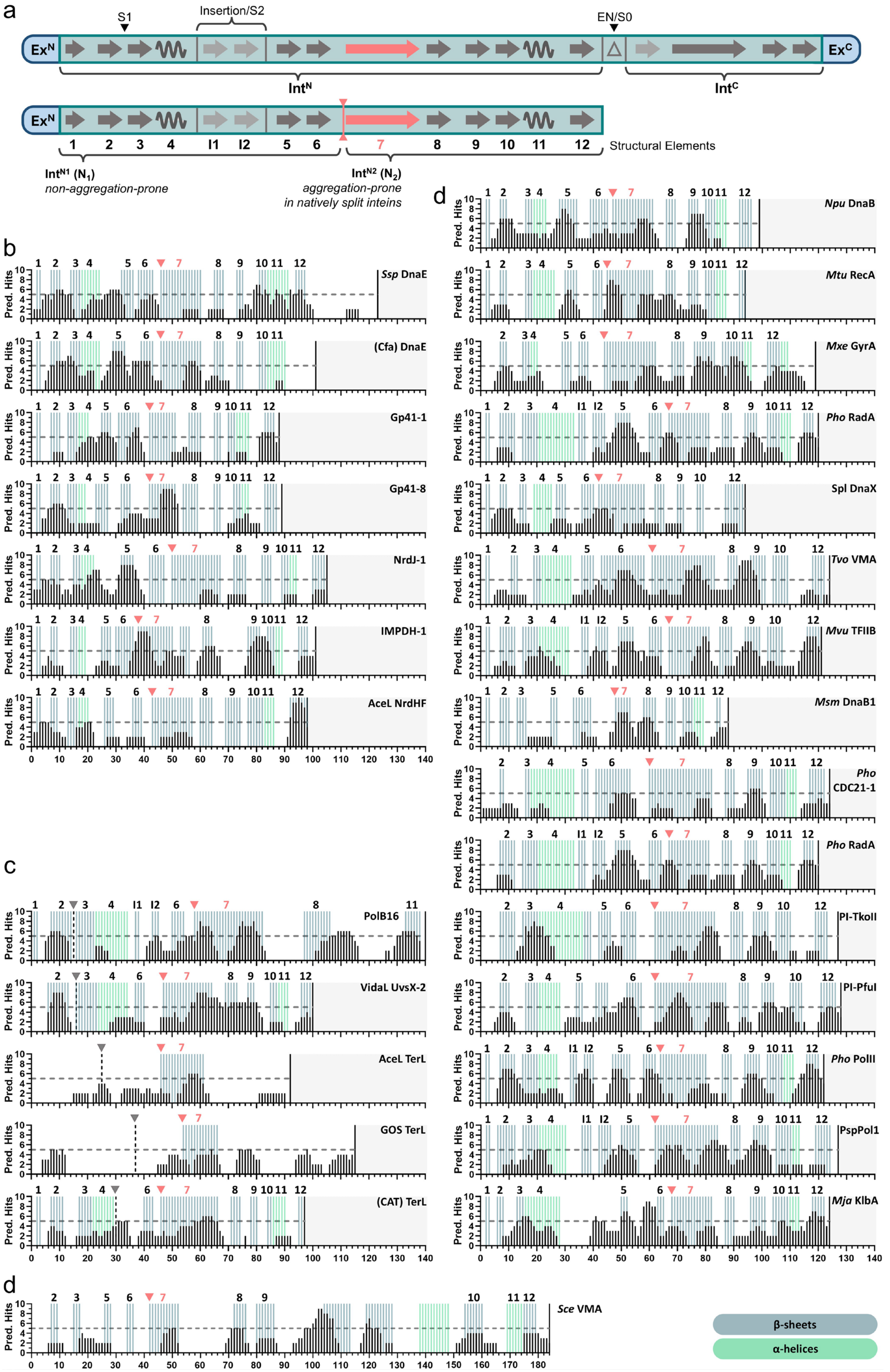
a
S1
Insertion/S2
EN/S0
ExN
ExC
IntN
IntC
Structural Elements
IntN1 (N1)
non-aggregation-prone
IntN2 (N2)
aggregation-prone
in natively split inteins
b
Ssp DnaE
(Cfa) DnaE
Gp41-1
Gp41-8
NrdJ-1
IMPDH-1
AceL NrdHF
c
PolB16
VidaL UvsX-2
AceL TerL
GOS TerL
(CAT) TerL
d
Sce VMA
Npu DnaB
Mtu RecA
Mxe GyrA
Pho RadA
Spl DnaX
Tvo VMA
Mvu TFIIB
Msm DnaB1
Pho CDC21-1
PI-TkoII
PI-PfuI
Pho PolII
PspPol1
Mja KlbA
Pred. Hits
β-sheets
α-helices

**Supplementary Figure 19** *In silico* aggregation-prone site prediction of commonly used inteins. **a** Illustration taken from Figure 7a. Shown are conserved secondary structure elements in the minimal intein horseshoe fold for *cis*-inteins (top) and the $Int^N$ fragment of split inteins (bottom). Additionally, one optional insertion of β-strands I1 & I2 is shown that is also present in the Aes split intein. The split positions corresponding to typically split (EN/S0) and atypically split (S1/S2) inteins are indicated (EN = intervening homing endonuclease domain).[31, 32] The long β-sheet in the $Int^N$ (structural element 7; marked in red) stretches across an axis of two-fold symmetry in the folded intein structure that has previously been used to define the $Int^N$ $N_1$ and $N_2$ lobes.[23] Here, however, the border between $N_1$ and $N_2$ segments was defined at the beginning of β-sheet 7, indicated by the vertical red line (see main text). **b** Sequence analysis by the consensus prediction algorithm AMYLPRED2[9] of the $Int^N$ part of several commonly used natively split inteins. Note that the structural data of Gp41-8, NrdJ-1 and IMPDH-1 inteins were predicted using AlphaFold3 and Phyre$^2$ due to the lack of experimental data causing potential artefacts in the *in silico* analysis. **c** Sequence analysis of the canonical $Int^N$ part of commonly used atypically split inteins. Note that the structural data of AceL TerL and GOS TerL inteins were predicted using AlphaFold3 and Phyre$^2$ due to the lack of experimental data causing potential artefacts in the *in silico* analysis. **d** Sequence analysis of the canonical $Int^N$ part of several commonly used maxi- and mini-inteins.

**Supplementary Table 8** Inteins and the corresponding canonical Int$^{N}$ sequences used to compare the predicted aggregation tendency.

| | Intein | Canonical Int$^{N}$ Sequence | pI |
|---|---|---|---|
| Natively Split Inteins | *Npu* DnaE | CLSYETEILTVEYGLLPIGKIVEKRIECTVYSVDNNGNIYTQPVAQWHDRGEQEVFEYCLEDGSLIRATKDHKFMTVDGQMLPIDEIFERELDLMRVDNLPN | 4.39 |
| | *Ssp* DnaE | CLSFGTEILTVEYGPLPIGKIVSEEINCSVYSVDPEGRVYTQAIAQWHDRGEQEVLEYELEDGSVIRATSDHRFLTTDYQLLAIEEIFARQLDLLTLENIKQTEEALDNHRLPFPLLDAGTIK | 4.36 |
| | *Cfa* DnaE (engineered) | CLSYDTEILTVEYGFLPIGKIVEERIECTVYTVDKNGFVYTQPIAQWHNRGEQEVFEYCLEDGSIIRATKDHKFMTTDGQMLPIDEIFERGLDLKQVDGLP | 4.43 |
| | Gp41-1 | CLDLKTQVQTPQGMKEISNIQVGDLVLSNTGYNEVLNVFPKSKKKSYKITLEDGKEIICSEEHLFPTQTGEMNISGGLKEGMCLYVKE | 5.34 |
| | Gp41-8 | CLSLDTMVVTNGKAIEIRDVKVGDWLESECGPVQVTEVLPIIKQPVFEIVLKSGKKIRVSANHKFPTKDGLKTINSGLKVGDFLRSRAK | 9.47 |
| | NrdJ-1 | CLVGSSEIITRNYGKTTIKEVVEIFDNDKNIQVLAFNTHTDNIEWAPIKAAQLTRPNAELVELEIDTLHGVKTIRCTPDHPVYTKNRGYVRADELTDDDELVVAI | 4.80 |
| | IMPDH-1 | CFVPGTLVNTENGLKKIEEIKVGDKVFSHTGKLQEVVDTLIFDRDEEIISINGIDCTKNHEFYVIDKENANRVNEDNIHLFARWVHAEELDMKKHLLIELE | 4.91 |
| | AceL NrdHF | ALLVGTKVTTKAGDKNIENITLEDEVLQFDMNTKDFSYTNPTKTQKVIRDEIYHFEGAGFDQKVSPNHRMIYEQGGEIKECLAKDFEPSEDKYFIIVE | 4.74 |
| | Aes123 PolB1 | SVVGDTIIDVSGKKMTIAEFYDSTPDVFMRRNDEARDWVKRVGGKTSLSVNTYSGEVERKNINYIMKHTVKKRMFKIKAGGKEVIVTADHSVMVKRDGKIIDVKPTEMKQTDRVVKWMLT | 9.76 |
| Contiguous Inteins (maxi and mini-Inteins) | *Ssp* DnaB | CISGDSLISLASTGKRVSIKDLLDEKDFEIWAINEQTMKLESAKVSRVFCTGKKLVYILKTRLGRTIKATANHRFLTIDGWKRLDELSLKEHIALPR | 9.59 |
| | *Npu* DnaB | CLAGDSLVTLVDSGLQVPIKELVGKSGFAVWALNEATMQLEKAIVSNAFSTGIKPLFTLTTRLGRKIRATGNHKFLTINGWKRLDELTPKEHLALPRNS | 9.80 |
| | *Mtu* RecA | CLAEGTRIFDPVTGTTHRIEDVVDGRKPIHVVAAAKDGTLHARPVVSWFDQGTRDVIGLRIAGGAILWATPDHKVLTEYGWRAAGELRKGDRVA | 8.17 |
| | *Mxe* GyrA | CITGDALVALPEGESVRIADIVPGARPNSDNAIDLKVLDRHGNPVLADRLFHSGEHPVYTVRTVEGLRVTGTANHPLLCLVDVAGVPTLLWKLIDEIKPGDYAVIQRSAFSVDCAGFAR | 5.43 |
| | *Mvu* TFIIB | SVDYSEPIIIKEKGEIKVVKIGELIDEIIKNSKNVRKDGILEIARCKDVEVIAFDSNYKFKFMPVSEVSRHPVSEMFEIVVEGNKKVRVTGSHSVFTVKDNEVVPIRVDDLRVGDILVLAK | 6.14 |
| | *Msm* DnaB1 | ALALDTPLPTPSGWTTMGDVAVGDHLLGPDGEPTRVVADTDVMLGRPCYVVEFSDGTAIVADAQHQWPTEHGVRITANLRAGMHTVVS | 4.59 |
| | *Pho* RadA | CFARDTEVYYENDTVPHMESIEEMYSKYASMNGELPFDNGYAVPLDNVFVYTLDIASGEIKKTRASYIYREKVEKLIEIKLSSGYSLKVTPSHPVLLFRDGLQWVPAAEVKPGDVVVGVR | 4.99 |
| | *Pho* CDC21-1 | AVDYDTEVLLGDGRKRKIGEIVEEAIKKAEKEGKLGRVDDGFYAPINLELYALDVRTLKVRKVKADIAWKRTTPEKMLRIRTKRGREIRVTPTHPFFTLEEGRIKTKKAYELKVGEKIATPREE | 9.69 |
| | *Pho* PolII | CFPGDTRILVQINGTPQRVTLKELYELFDEEHYESMVYVRKKPKVDIKVYSFNPEEGKVVLTDIEEVIKAPATDHLIRFELELGSSFETTVDHPVLVYENGKFVEKRAFEVREGNIIIIIDE | 4.82 |
| | *Spl* Dnax | ALTGDALILSDRGWLRIDDPTLQECRVLSYNESTQQWEWQQVLRWLDQGVRETWKIKTFQTEIKCTGNHLIRTDKGWIKAANITPKMKILSPEI | 8.03 |
| | *Mja* KlbA | ALAYDEPIYLSDGNIINIGEFVDKFFKKYKNSIKKEDNGFGWIDIGNENIYIKSFNKLSLIIEDKRILRVWRKKYSGKLIKITTKNRREITLTHDHPVYISKTGEVLEINAEMVKVGDYIYIPK | 9.28 |
| | PI-PfuI | CIDGKAKIIFENEGEEHLTTMEEMYERYKHLGEFYDEEYNRWGIDVSNVPIYVKSFDPESKRVVKGKVNVIWKYELGKDVTKYEIITNKGTKILTSPWHPFFVLTPDFKIVEKRADELKEGDILIGGM | 5.28 |
| | PI-TkoII | SILPEEWLPVLEEGEVHFVRIGELIDRMMEENAGKVKREGETEVLEVSGLEVPSFNRRTNKAELKRVKALIRHDYSGKVYTIRLKSGRRIKITSGHSLFSVRNGELVEVTGDELKPGDLVAVPRRLE | 6.83 |
| | PspPol-1 | SILPEEWVPLIKNGKVKIFRIGDFVDGLMKANQGKVKKTGDTEVLEVAGIHAFSFDRKSKKARVMAVKAVIRHRYSGNVYRIVLNSGRKITITEGHSLFVYRNGDLVEATGEDVKIGDLLAVPRSVN | 9.88 |
| | *Sce* VMA | CFAKGTNVLMADGSIECIENIEVGNKVMGKDGRPREVIKLPRGRETMYSVVQKSQHRAHKSDSSREVPELLKFTCNATHELVVRTPRSVRRLSRTIKGVEYFEVITFEMGQKKAPDGRIVELVKEVSKSYPISEGPERANELVESYRKASNKAYFEWTIEARDLSLLGSHVRKATYQTYAPILY | 9.32 |
| | *Tvo* VMA | CVSGETPVYLADGKTIKIKDLYSSERKKEDNIVEAGSGEEIIHLKDPIQIYSYVDGTIVRSRSRLLYKGKSSYLVRIETIGGRSVSVTPVHKLFVLTEKGIEEVMASNLKVGDMIAAVAESASE | 6.17 |
| Atypically Split Inteins | VidaL UvsX-2 | CLPKEAVVQIRLTKKGMIEEKKVTVQELRELYLSGEYTIEIDTPDGYQTIGKWFDKGVLSMVRVATATYETVCAFNHMIQLADNTWVQACELDVGVDIQT | 4.75 |
| | PolB-16 | SVHGKTHVFIRSIKNMQEAKIDIKSLYDSLAKKYDVQHKNSYEVIYPKGYEIKVLGNKYVKLVAMSRHKTQKHLVKIVVKSEKTIDSLDPIRQKSLLKKQDEVVVTTDHICMVYNDDHFFENVNAKNLKVGNYVSVYDEA | 9.33 |
| | AceL TerL | CVYGDTMVETEDGKIKIEDLYKRLAMFRTNTNNIKILSPNGFSNFNGIQKVERNLYQHIIFDDDTEIKTSINHPFGKDKILARDVKVGDYLN | 6.11 |
| | GOS TerL | SISQESYINIEVNGKVETIKIGDLYKKLSFNERKFNEMKLPESVVKNNINLKIETPYGFENFYGVNKIKKDKYIHLEFTNGEKLKCSLDHPLSTIDGIVKAKDLDKYTEVYTKFG | 8.51 |
| | CAT TerL (engineered) | CLSGDTMIEILDDDGIIQKISMEDLYQRLAMFKLNTKNIKVLTPSGFKSFSGIQKVYKPFYHHIIFDDGSEIKCSDNHSFGKDKIKASTIKVGDYLQ | 7.00 |

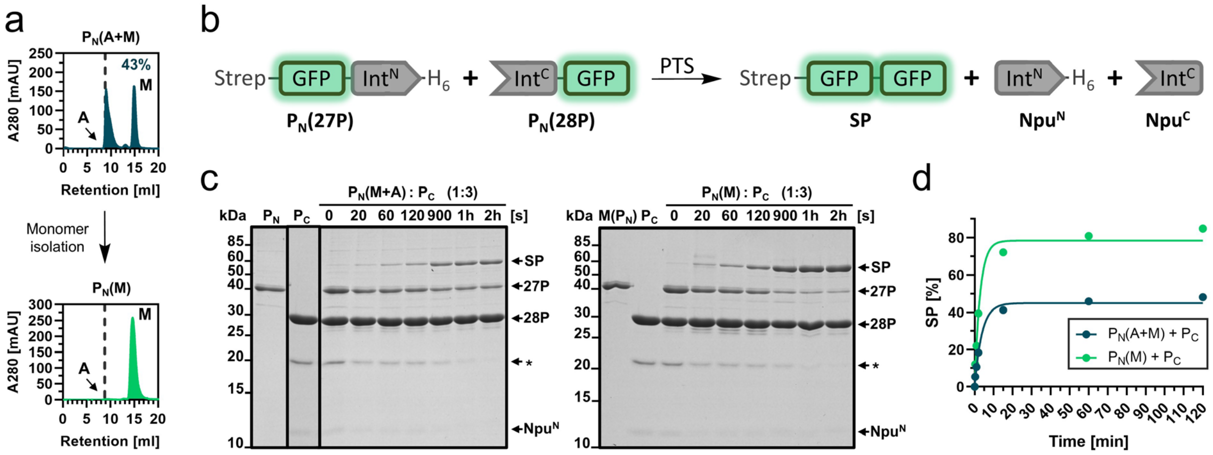


**Supplementary Figure 20** Int$^N$ aggregation into inactive species in the natively split *Npu* DnaE intein. **a** SEC-analysis of **27P** before (upper panel) and after (lower panel) monomer isolation. These two preparations were used for the reactions in **c**. **b** Scheme of the PTS reaction using Strep-GFP-Npu$^N$ (**27P**) and Npu$^C$-GFP (**28P**). **c** Coomassie-stained SDS-PAGE analysis of the PTS reaction (under reducing conditions) with **27P** and **28P** used in three-fold molar excess (3 µM and 9 µM, 37°C) without prior SEC-purification (left panel) and after monomer isolation of **27P** ($P_N$(M(**27P**)) (right panel). **d** Time-course of the PTS reactions in **c** fitted to a one-phase exponential equation. $P_N$(**27P**) = 40.8 kDa, $P_C$(**28P**) = 31.4 kDa, Npu$^N$ = 12.8 kDa, Npu$^C$ = 4.1 kDa, SP = 56.3 kDa. SP = splice product. For panel **d**, *n* = 2 technical replicates. Data are presented as mean ± s.d. normalized to the molecular weight.

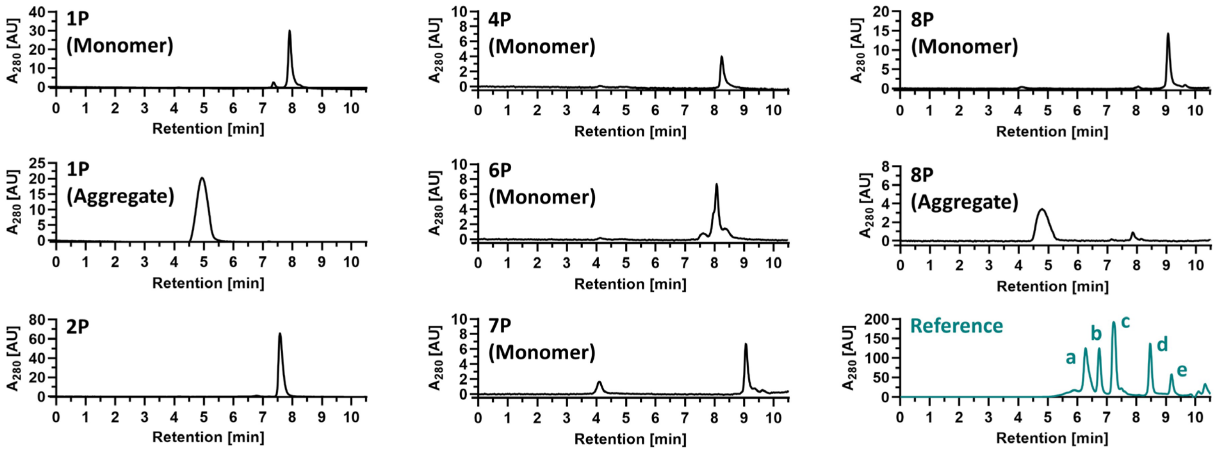


**Supplementary Figure 21** SEC analysis of the purified and isolated monomeric or aggregated proteins used throughout this study. The reference SEC UV-chromatogram indicates the usual retention time of globular proteins between 12 kDa and 200 kDa while the aggregated species show a retention time of 4 – 5.5 min. Proteins were analyzed on AdvanceBio SEC 200Å 1.9 µm, 4.6 × 300 mm pre-packed column (Agilent) at a follow rate of 0.35 mL/min. (a: β-amylase (200 kDa), b: alcohol dehydrogenase (150 kDa), c: bovine serum albumin (66 kDa), d: carbonic anhydrase (29 kDa), e: cytochrome c (12.4 kDa).

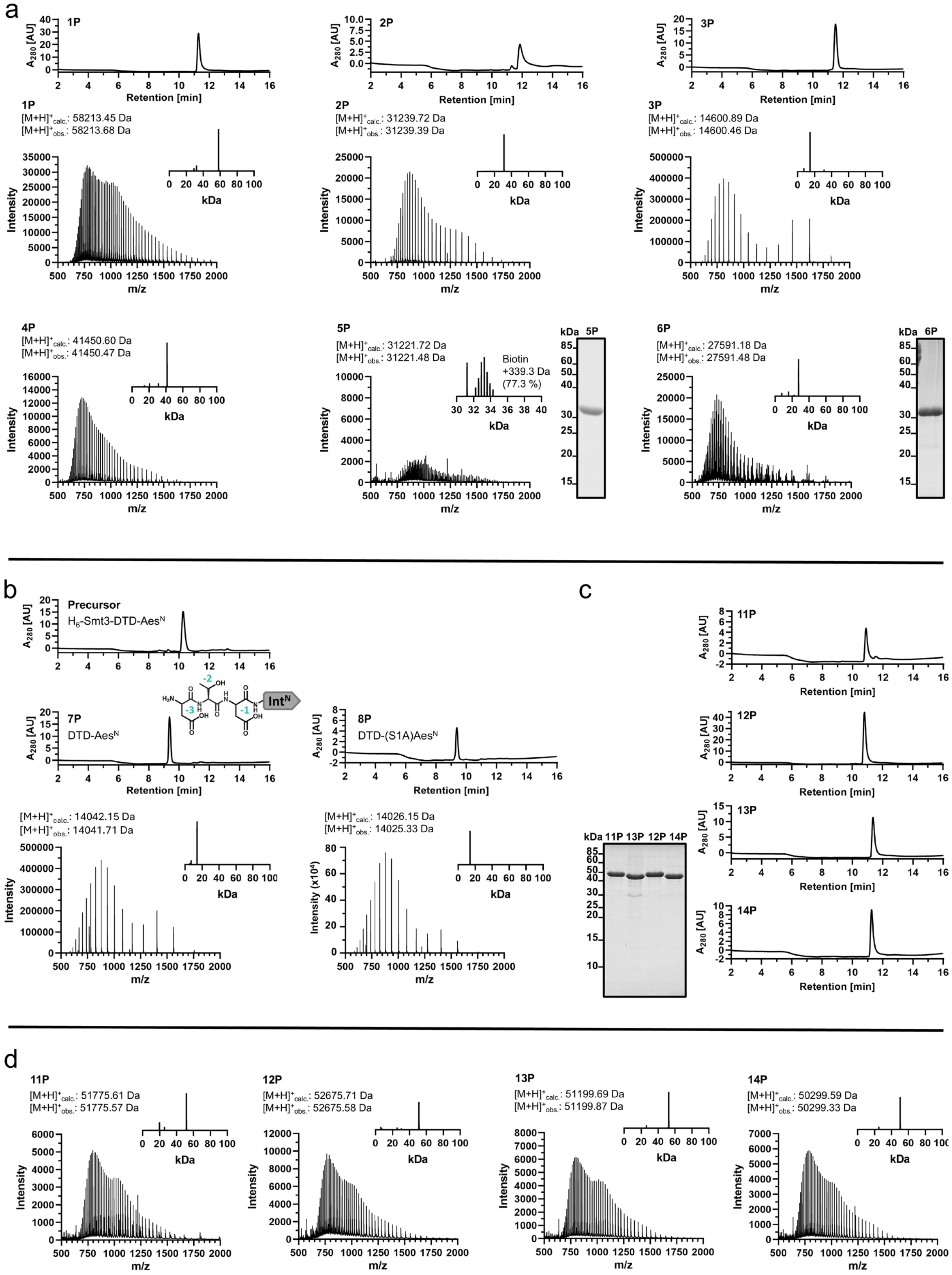


**Supplementary Figure 22** Continued on next page.

e

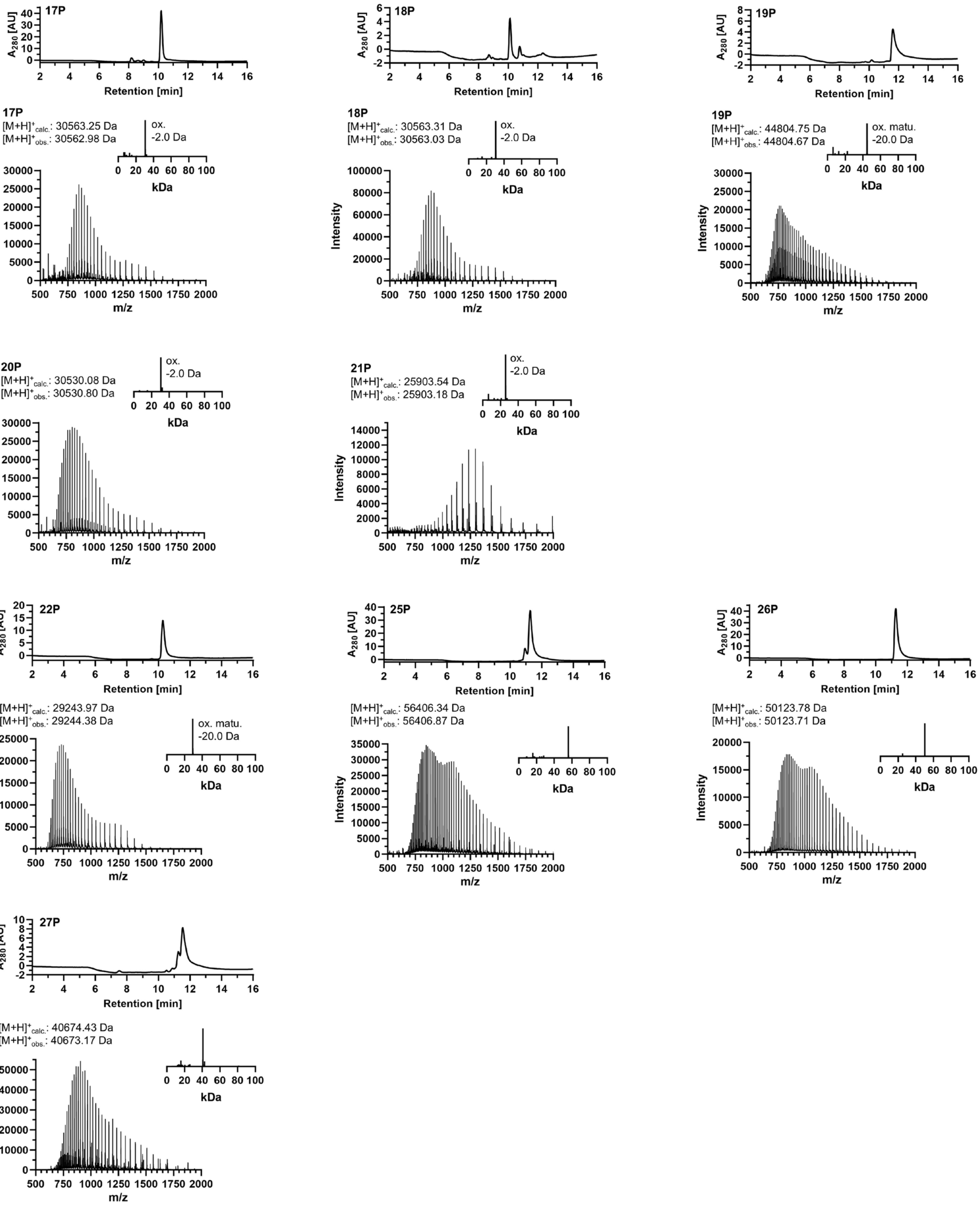


**Supplementary Figure 22** Analytics to the used constructs. **a** RP-HPLC UV-chromatograms (280 nm) to monitor purity of purified constructs **1P**, **2P** and **3P**. ESI-MS analysis of purified constructs **1P**, **2P**, **3P**, **4P**, **5P** (labeled via NHS-C2-Biotin) and **6P**. Shown are the original MS and the deconvoluted spectra (insets). SDS-PAGE analysis to monitor purity of purified constructs **5P** (labeled via NHS-C2-Biotin) and **6P**. **b** RP-HPLC UV-chromatograms to monitor purity of purified constructs **7P**, its precursor construct with a $H_6$-Smt3 tag (exemplary shown) and **8P**. ESI-MS analysis of purified constructs **7P** and **8P**. **c** RP-HPLC UV-chromatograms and SDS-PAGE analysis to monitor purity of purified constructs **11P**, **12P**, **13P** and **14P**. **d** ESI-MS analysis of purified constructs **11P**, **12P**, **13P** and **14P**. (E) RP-HPLC UV-chromatograms to monitor purity of purified constructs **17P, 18P**, **19P**, **22P**, **25P**, **26P** and **27P**. ESI-MS analysis of the used constructs to prove protein identity and redox state of the purified constructs **17P**, **18P**, **19P**, **20P**, **21P**, **SP(20P-21P)-Cy5**, **22P**, **25P**, **26P** and **27P** (ox.: oxidized; matu.: chromophore maturation within GFP).

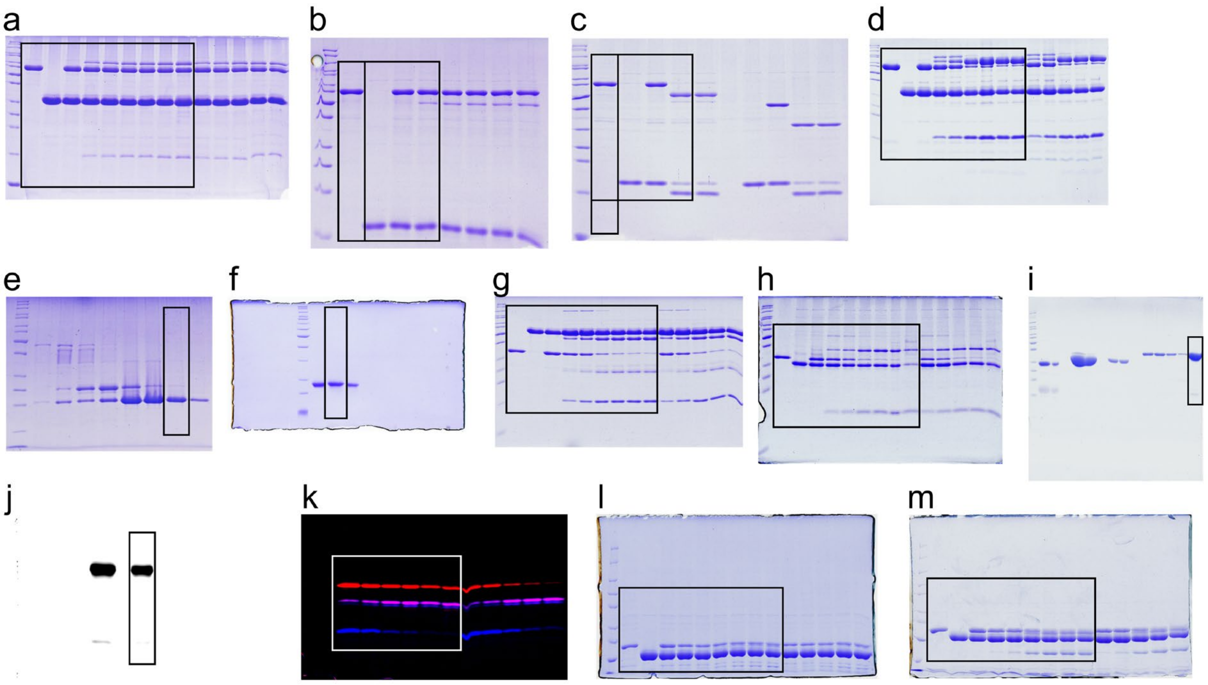


**Supplementary Figure 23** Unprocessed SDS-PAGE images of the figures shown in the main text. The black frame indicates the section used for the figures **a** Figure 1b, **b** Figure 2b,c, **c** Figure 2b,c, **d** Figure 5b, **e** Figure 5d, **f** Figure 5d, **g** Figure 6b, **h** Figure 6d, **i** Figure 6d, **j** Figure 6d, **k** Figure 6h, **l** Figure 7e, **m** Figure 7e.

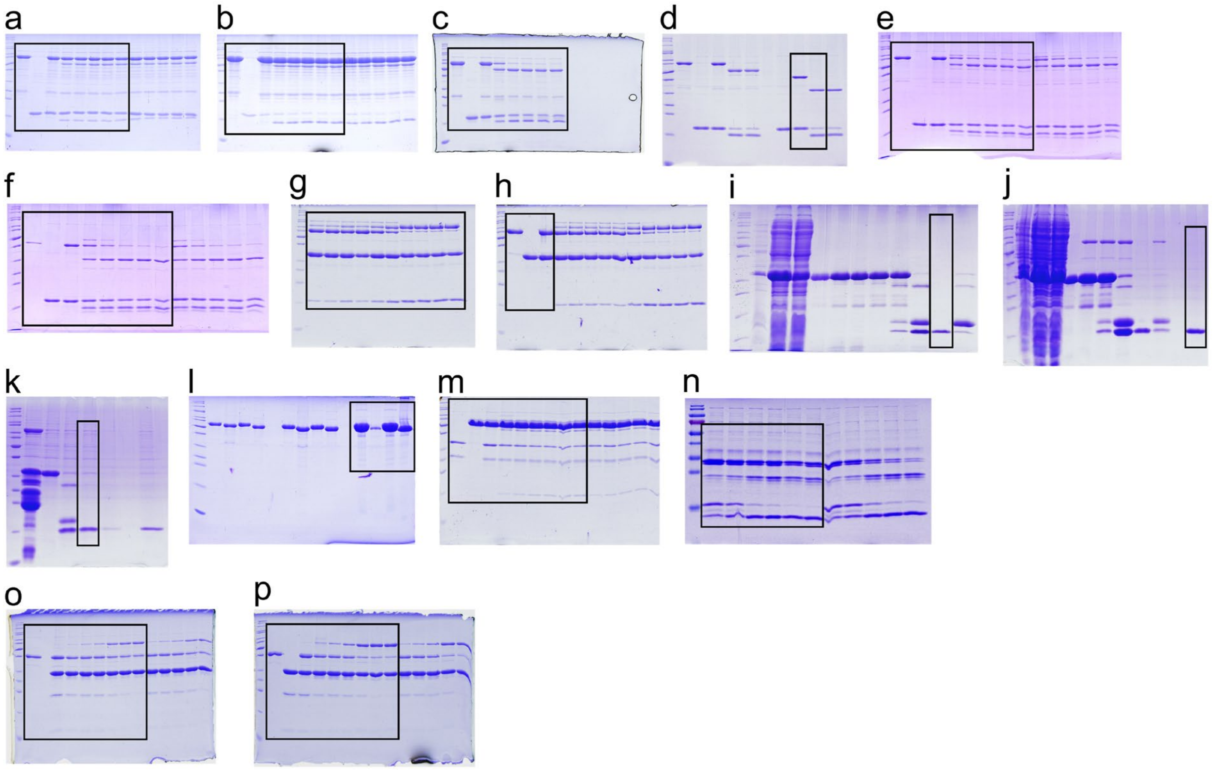


**Supplementary Figure 24** Unprocessed SDS-PAGE images of the supplementary figures. The black frame indicates the section used for the figures **a** Supplementary Fig. 2b, **b** Supplementary Fig. 2c, **c** Supplementary Fig. 3b, **d** Supplementary Fig. 4b, **e** Supplementary Fig. 4d, **f** Supplementary Fig. 4e, **g** Supplementary Fig. 6b, **h** Supplementary Fig. 6b, **i** Supplementary Fig. 8b, **j** Supplementary Fig. 9a, **k** Supplementary Fig. 9a, **l** Supplementary Figure 11b, **m** Supplementary Fig. 16c, **n** Supplementary Fig. 18d, **o** Supplementary Fig. 20c, **p** Supplementary Fig. 20c.
